# The SAMPL6 SAMPLing challenge: Assessing the reliability and efficiency of binding free energy calculations

**DOI:** 10.1101/795005

**Authors:** Andrea Rizzi, Travis Jensen, David R. Slochower, Matteo Aldeghi, Vytautas Gapsys, Dimitris Ntekoumes, Stefano Bosisio, Michail Papadourakis, Niel M. Henriksen, Bert L. de Groot, Zoe Cournia, Alex Dickson, Julien Michel, Michael K. Gilson, Michael R. Shirts, David L. Mobley, John D. Chodera

**Affiliations:** Computational and Systems Biology Program, Sloan Kettering Institute, Memorial Sloan Kettering Cancer Center, New York, NY 10065; Tri-Institutional Training Program in Computational Biology and Medicine, New York, NY 10065; Department of Chemical and Biological Engineering, University of Colorado Boulder, Boulder, CO 80309; Skaggs School of Pharmacy and Pharmaceutical Sciences, University of California, San Diego, La Jolla, CA 92093, USA; Max Planck Institute for Biophysical Chemistry, Computational Biomolecular Dynamics Group, Göttingen, Germany; Biomedical Research Foundation, Academy of Athens, 4 Soranou Ephessiou, 11527 Athens, Greece; EaStCHEM School of Chemistry, University of Edinburgh, David Brewster Road, Edinburgh EH9 3FJ, UK; Atomwise, 717 Market St Suite 800, San Francisco, CA 94103; Department of Biochemistry and Molecular Biology, Michigan State University, East Lansing, MI, USA; Department of Computational Mathematics, Science and Engineering, Michigan State University, East Lansing, MI, USA; Department of Pharmaceutical Sciences and Department of Chemistry, University of California, Irvine, California 92697, USA

## Abstract

Approaches for computing small molecule binding free energies based on molecular simulations are now regularly being employed by academic and industry practitioners to study receptor-ligand systems and prioritize the synthesis of small molecules for ligand design. Given the variety of methods and implementations available, it is natural to ask how the convergence rates and final predictions of these methods compare. In this study, we describe the concept and results for the SAMPL6 SAMPLing challenge, the first challenge from the SAMPL series focusing on the assessment of convergence properties and reproducibility of binding free energy methodologies. We provided parameter files, partial charges, and multiple initial geometries for two octa-acid (OA) and one cucurbit[8]uril (CB8) host-guest systems. Participants submitted binding free energy predictions as a function of the number of force and energy evaluations for seven different alchemical and physical-pathway (i.e., potential of mean force and weighted ensemble of trajectories) methodologies implemented with the GROMACS, AMBER, NAMD, or OpenMM simulation engines. To rank the methods, we developed an efficiency statistic based on bias and variance of the free energy estimates. For the two small OA binders, the free energy estimates computed with alchemical and potential of mean force approaches show relatively similar variance and bias as a function of the number of energy/force evaluations, with the attach-pull-release (APR), GROMACS expanded ensemble, and NAMD double decoupling submissions obtaining the greatest efficiency. The differences between the methods increase when analyzing the CB8-quinine system, where both the guest size and correlation times for system dynamics are greater. For this system, nonequilibrium switching (GROMACS/NS-DS/SB) obtained the overall highest efficiency. Surprisingly, the results suggest that specifying force field parameters and partial charges is insufficient to generally ensure reproducibility, and we observe differences between seemingly converged predictions ranging approximately from 0.3 to 1.0 kcal/mol, even with almost identical simulations parameters and system setup (e.g., Lennard-Jones cutoff, ionic composition). Further work will be required to completely identify the exact source of these discrepancies. Among the conclusions emerging from the data, we found that Hamiltonian replica exchange—while displaying very small variance—can be affected by a slowly-decaying bias that depends on the initial population of the replicas, that bidirectional estimators are significantly more efficient than unidirectional estimators for nonequilibrium free energy calculations for systems considered, and that the Berendsen barostat introduces non-negligible artifacts in expanded ensemble simulations.

## 1 Introduction

Predicting the binding free energy between a receptor and a ligand has attracted a great deal of attention due to its potential to speed up small-molecule drug discovery [1]. Among the methodologies that have been developed to carry out this task, physics-based methods employing classical force fields are starting to be routinely used in drug development projects and demonstrate success in real lead optimization scenarios [2–5]. These technologies are also often employed to obtain mechanistic insights into the physics of binding such as the discovery of binding poses [6] and pathways [7], or attempts at providing intuitive guidance on how to improve ligand binding potency [8]. However, the applicability domain of these models is currently limited to a narrow portion of the accessible chemical space for small molecules, and well-behaved protein-ligand systems that do not undergo significant conformational changes or solvent displacement on timescales larger than a few tens of nanoseconds [9, 10]. For this reason, much work has been directed at benchmarking and improving both the predictive accuracy and efficiency of these computational protocols [11–14]. The computational cost of a method, in particular, is a critical factor that enters the decision-making process both in academia and industry. For example, to achieve maximum impact in drug discovery, methods should achieve high-confidence predictions on a timescale sufficiently short to inform synthetic decisions—with increasingly rapid predictions in principle enabling quicker cycles of idea generation and testing. [2, 9, 10]. More generally, unconverged results and systematic errors can compromise the assessment of the accuracy of a force field through fortuitous cancellation/amplification of error, with immediate consequences on the optimization of free energy protocols and molecular models. Determining which methods are capable of most rapidly reducing the error is thus critical to enable not only prospective studies in drug discovery, but also to carry out meaningful benchmarks and optimize molecular models with useful turnaround times.

### 1.1 Multiple sources contribute to the error of the estimate

In the rest of the work, we refer to the model of the system to include any element affecting the potential energy function we intend to simulate (e.g., force field, charge model, protonation states, ion concentrations). The model, together with the thermodynamic parameters (e.g., temperature, pressure) and the definition of the binding site completely determine the theoretical binding free energy Δ*G*_*θ*_ through the associated ratio of partition functions [15]. The output of a binding free energy method is a statistical estimate of the free energy, a random variable Δ*G*_calc_ = Δ*G_θ_* + *ϵ*, which is an estimate of Δ*G_θ_* up to an error *ϵ* that generally depends on the method itself and the computational cost invested in the calculation. We consider a method to be efficient if it can quickly reduce the standard deviation of Δ*G*_calc_ (i.e., std(Δ*G*_calc_) = std(*ϵ*)) and its bias, which is defined as 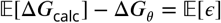, where the expected value is intended over multiple independent executions of the method of the same computational cost.

Assuming a method is exact and correctly implemented, the major source of statistical error is arguably connected to the sampling strategy adopted by the method. Due to the rough potential energetic landscape, short molecular dynamics (MD) or Monte Carlo (MC) simulations (where for proteins, short can still be 100s of ns) can miss entire areas of configurational space that contribute significantly to the partition functions, or have insufficient time to accurately estimate the relative populations of the different free energy basins. This introduces bias into the affinity estimates. Enhanced sampling strategies such as metadynamics [16, 17], replica exchange [18–20], and expanded ensemble [21] methodologies are designed to increase the sampling efficiency along one or a few collective variables (CV), although their effectiveness strongly depends on the choice of the CV. Moreover, even in the limit of infinite sampling, common non-Metropolized sampling strategies such as Verlet integration and Langevin dynamics can introduce systematic bias due to the integration error. While the magnitude of this bias has not been studied extensively in free energy calculations of host-guest or protein-ligand systems, it was shown to be significant in simple systems depending on the size of time step, and choice of integrator [22, 23]. Finally, while many different free energy estimators (e.g., exponential averaging, BAR, MBAR, thermodynamic integration) are provably asymptotically unbiased and consistent, these behaviors break down for finite sample sizes, and their bias and variance decay differently as a function of the number of independent samples [24].

### 1.2 Comparing the efficiency of methods requires eliminating confounding factors

Any simulation parameter altering the potential energy landscape of the end states can alter the energetic barriers between metastable states and change the theoretical binding free energy Δ*G*_*θ*_. The former impact the correlation times of the dynamics and thus the convergence rates of methods, while the latter makes it harder to detect systematic biases introduced by the methodologies. There are several examples in the literature noting differences in binding free energy predictions between different methods, but in which it was impossible to determine whether this was due to other differences in system preparation, insufficient sampling, or shortcomings of the methodology [25–28]. Consequently, it is important to test the methods on the same set of molecular systems, using the same model. The latter, in particular, requires specifying force field parameters and partial charges, but also other components of the simulation, such as ion concentrations and the treatment of long-range interactions (e.g. PME, reaction field, Lennard-Jones cutoff, dispersion correction). Treating long-range interactions equivalently is particularly challenging due to differences in functional forms, implementations, and options supported by the various software packages, including small discrepancies in the value of the Coulomb constant [29, 30]. Establishing a set of simulation settings that minimizes these differences does not prevent systematic bias due to sampling issues, but it makes it possible to detecting by comparing calculations performed with independent methods and/or starting from different initial configurations.

Comparing multiple independent methods on the same set of systems currently requires substantial pooled technical expertise and coordination as well as significant computational resources. Confidently estimating the bias necessitates very long simulations and consensus between methods. Moreover, in the absence of a reliable strategy for uncertainty estimation, multiple independent replicates are vital for a correct ranking of performance of different methods. Previous work investigating the reproducibility of relative alchemical hydration free energy calculations across four molecular packages uncovered various issues and challenges in comparing across simulation packages and resulted in various bug fixes [30]. However, the reproducibility and efficiencies of various simulation-based approaches has not yet been evaluated in the context of binding free energy calculations, which is the focus of this work.

### 1.3 We need robust general strategies to measure the efficiency of binding free energy calculations

While there are generally established ways of measuring the accuracy of free energy calculation protocols with respect to experimental measurements, there is no consensus or standard practice regarding how to measure the efficiency of a method. A study focusing on accuracy of free energy calculations typically ranks different protocols and methodologies using commonly adopted correlation and error statistics describing how well experimental affinities are predicted (e.g. R^2^, MUE, and RMSE) [25, 26, 31–34]. On the other hand, the efficiency of sampling strategies in the context of free energy calculations has been evaluated in many different ways in the past, none of which we found completely adequate for the goal of this challenge.

In some cases, one or more system-specific collective variables associated with a slow degree of freedom can be directly inspected to verify thorough sampling [27, 35, 36]. This strategy requires extensive knowledge of the system and is not generally applicable to arbitrary receptor-ligand systems. Moreover, free energy calculations commonly involve simulating the same system in multiple intermediate states—which are not always physical intermediates—that do not necessarily have the same kinetic properties. Commonly, quantitative comparisons of performance are based on the standard deviation of the free energy estimates after roughly the same computational cost [37–40]. This statistic, however, does not quantify the bias, which is, in general, not negligible. In principle, one can test the methods on a set of molecules composed of quickly converging systems, or the calculations can be run for a very long time in order to increase our confidence in the assumption that the bias has decayed to zero. However, neither of these two scenarios necessarily reflect the performance of the method in a real scenarios, which ordinarily involves complex receptor-ligand systems with long correlation times and simulations of a few nanoseconds per intermediate state. Alternatively, other statistics such as acceptance rate and mean first-passage time have been reported [39–41], but these statistics are method-specific, and not necessarily indicative of the error of the free energy estimate. Another common strategy to assess the efficiency of a method is the visual inspection of the decay of some error metric [42, 43], but this qualitative analysis is not scalable nor statistically quantifiable when the number of methods and systems considered increases. Finally, there is a large body of theoretical work focusing on the efficiency of estimators and protocols in free energy calculations [24, 37, 40, 42, 44, 45], but in many cases, they are difficult to apply to practical scenarios. The results rely on the assumption of independent samples and often focus on the asymptotic regime, both of which are conditions that may not apply in practice.

### 1.4 Objectives of the SAMPL6 SAMPLing challenge

In this work, we present the design and the results of the first round of the community-wide **SAMPLing challenge**. Our goal is to establish a statistical inference framework for the quantitative comparison of the convergence rates of modern free energy methods on a host-guest benchmark set. Moreover, we assess the level of agreement that can be reached by different methods and software packages when provided identical charges, force field parameters, systems, input geometries, and (when possible) simulation parameters. These objectives are distinct from the goal of the traditional SAMPL host-guest accuracy binding challenge, which instead focuses on the prediction of experimental values and ignores the computational cost of methods. Contrary to the accuracy challenge, which accepted data from widely different methods such as docking [46], QM [47] and QM/MM [48, 49] calculations, or movable type [50, 51] predictions, we limited the scope of this first round of the challenge to force field-based methodologies that should provide identical free energy estimates. With this first round, we lay the groundwork for future SAMPLing challenges and publish a protocol that can be used by independent studies that are similar in scope.

## 2 Challenge design

### 2.1 Selection of the three host-guest systems

The host-guest systems used here are drawn from the SAMPL6 host-guest binding challenge [26]. We selected 5-hexenoic acid (OA-G3) and 4-methylpentanoic acid (OA-G6) as guest molecules of the octa-acid host (OA), and quinine (CB8-G3) for the cucurbit[8]uril (CB8) host (***Figure 1***). The three guests that were chosen for the challenge include molecules resembling typical druglike small molecules (i.e. CB8-G3) and fragments thereof (i.e OA-G3/G6). Quinine was an obvious choice for the former category as it is currently recommended as the second-line treatment for malaria by the World Health Organization [52]. Originally, two octa-acid guests with very similar structures were purposely included to make them easily amenable to relative free energy calculations. However, we did not receive any submission utilizing relative free energy calculations.

**Figure 1.**
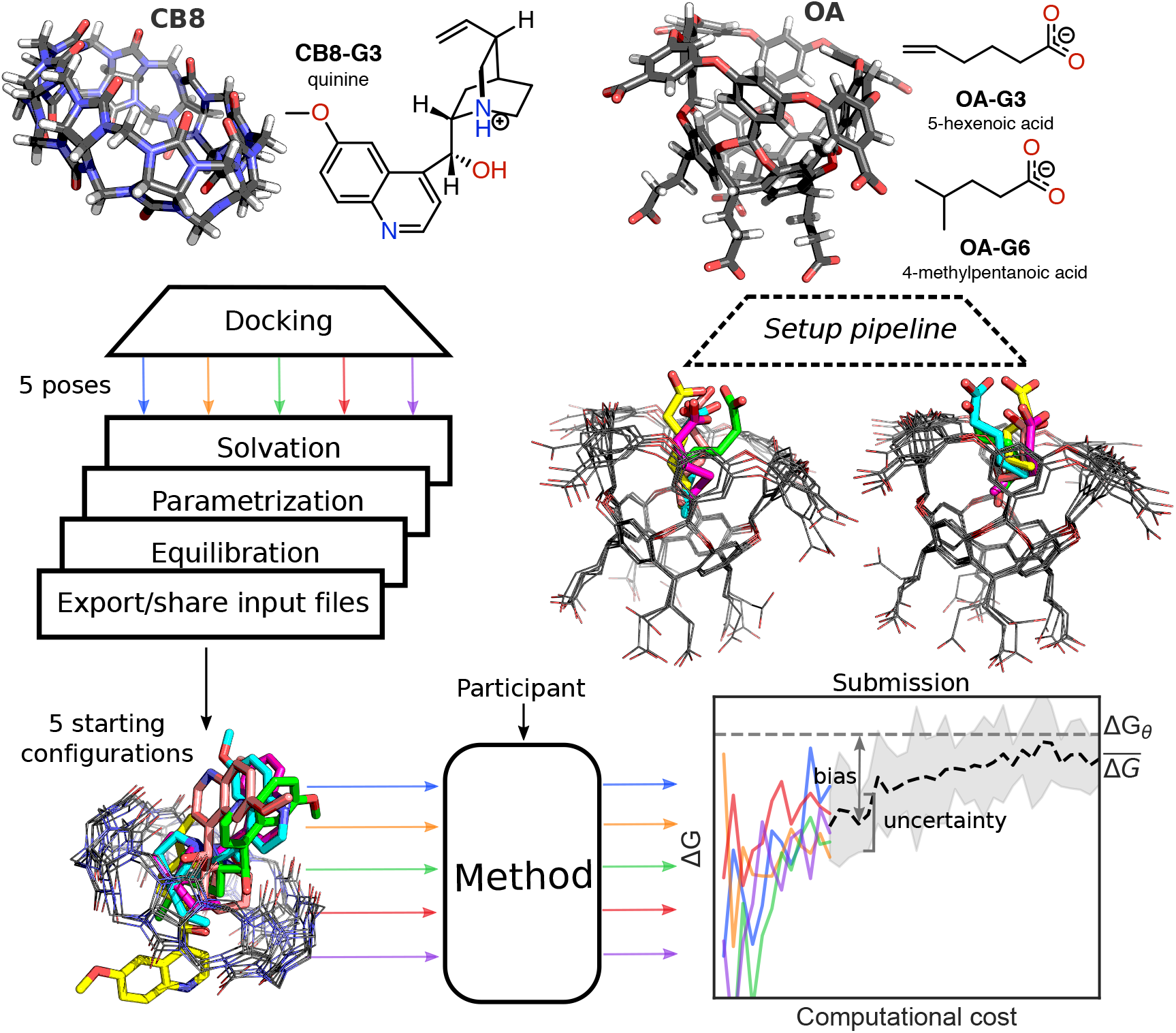
Challenge overview and initial conformations of the host-guest systems featured in the SAMPLing challenge. The three-dimensional structures of the two hosts (i.e. CB8 and OA) are shown with carbon atoms represented in black, oxygens in red, nitrogens in blue, and hydrogens in white. Both the two-dimensional chemical structures of the guest molecules and the three-dimensional structures of the hosts entering the SAMPLing challenge are shown in the protonation state used for the molecular simulations. We generated five different initial conformations for each of the three host-guest pairs through docking, followed by a short equilibration with Langevin dynamics. The three-dimensional structure overlays of the five conformations for CB8-G3, OA-G3, and OA-G6 are shown from left to right in the figure with the guests’ carbon atoms colored by conformation. Participants used the resulting input files to run their methods in five replicates and submitted the free energy trajectories as a function of the computational cost. We analyzed the submissions in terms of uncertainty of the mean binding free energy 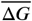 estimate and its bias with respect to the asymptotic free energy Δ*G*_*θ*_.

Both supramolecular hosts have been extensively described in the literature [11, 53–56] and featured in previous rounds of the host-guest binding SAMPL challenge [25, 57, 58]. From the perspective of assessment of binding free energy methodologies, host-guest systems serve as attractive alternatives to protein-ligand systems as they generally do not undergo large conformational reorganizations and have limited number of atoms, which helps the exploration of larger timescales and reducing the uncertainty of the binding affinity estimates. At the same time, this class of systems provides several well-understood challenges for standard simulation techniques. Hosts in the cucurbituril and octa-acid families have been found to bind ions and undergo wetting/dewetting processes governed by timescales on the order of a few nanoseconds [59, 60]. Moreover, the symmetry of CB8 and OA results in multiple equivalent (and often kinetically-separated) binding modes that have to be sampled appropriately or accounted for by applying a correction term [61]. Finally, ligands with net charges can introduce artifacts in alchemical free energy calculations when Ewald methods are used to model long-range electrostatic interactions. There are several approaches for eliminating these errors, but disagreements about the optimal strategy persist [62–65].

### 2.2 Challenge overview

As illustrated in ***Figure 1***, we asked the participants to run five replicate free energy calculations for each of the three host-guest systems using predetermined force field and simulation parameters and starting from five different conformations that we made available in a GitHub repository (https://github.com/samplchallenges/SAMPL6/tree/master/host_guest/SAMPLing) in the form of input files compatible with common molecular simulation packages (i.e., AMBER, CHARMM, DESMOND, GROMACS, LAMMPS, and OpenMM). Participants were asked to submit binding free energy estimates and, optionally, associated uncertainty estimates as a function of the computational cost of their methodologies. More specifically, the submitted data was required to report 100 free energy estimates computed at regular intervals using the first 1%, …, 100% of the samples, which was defined as the amount of samples collected after 1%, …, 100% of the combined total number of force and energy evaluations performed for the calculation.

To rank the performance of methods, we used a measure of efficiency developed in this work (described in the next section) based on estimates of bias and uncertainty of the predictions obtained from the replicate data. To facilitate the analysis, participants were asked to run the same number of force and energy evaluations for all the five replicate calculations of the same system, although the total number of force and energy evaluations could be different for different systems and different methods. Besides the total number of force and energy evaluations, the submissions included also wall-clock time and, optionally, total CPU/GPU time for each replicate as measures of the computational cost. However, due to the significant differences in the hardware employed to run the simulations, this information was not considered for the purpose of comparing the performance of different methods.

### 2.3 Development of an efficiency statistic for free energy methods

In order to rank performance of methods using standard statistical inference tools, we developed a statistic that captures our meaning of efficiency. Unlike what standardly used in the literature (see Section 1.3), we require a measure of the (in)efficiency of a free energy methodology that can simultaneously *(1)* take into account both bias and variance of the free energy estimate, *(2)* summarize the performance of a method over a range of computational costs of interest, *(3)* easily be computed without previous system-specific knowledge (e.g. knowledge of the slowest degrees of freedom).

#### Mean error as an inefficiency statistic

In this section, we propose a measure of efficiency of method X based on the time-averaged root mean square error (RMSE) of the bidning free energy predicted by method X, Δ*G_X_*, with respect to the theoretical binding free energy determined by the model, Δ*G_θ_*

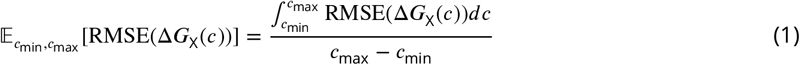

where [*c*_min_, *c*_max_] is the range of computational cost of interest, and

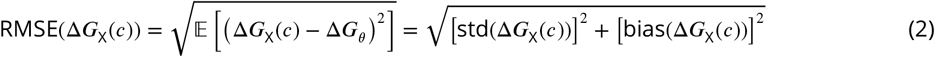

where the expected value, standard deviation, and bias functions are intended over all possible realizations (i.e. replicates) of the free energy calculation after investing a computational cost *c*. This metric satisfies all our requirements. Given the large differences in hardware among the submissions, we chose to measure the computational cost in number of force/energy evaluations rather than CPU or wall-clock time.

More generally, we can consider the *mean error*

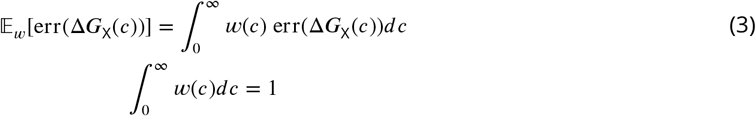

where the normalized weight function *w*(*c*) can be chosen to limit the average over a finite range of *c* (i.e. setting *w*(*c*) = 0 outside some interval), or based on the uncertainty of the estimate of the error statistic err, or also to satisfy other constraints such as the inclination of investing *c* to obtain a free energy prediction within a workflow. In the analysis, we always chose a uniform weight function as in (Eq. 1), but we also report the statistics computed using the standard deviation and absolute bias error functions

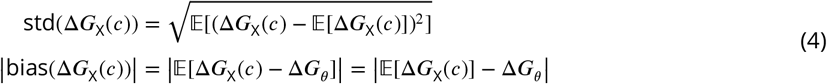

#### The relative efficiency is a robust statistic when data span different ranges of computational cost

The mean error of two methods is sensitive to the interval [*c*_min_*, c*_max_] considered, and thus it can be directly compared only if computed for the same interval of computational cost (see Appendix 1 and SI Figure 4 in the supporting information). However, the calculations submitted by participants have very different lengths, and computing the statistic on the largest range of computational cost shared by all methods would mean discarding between 50% and 75% of the data points for most submissions.

Instead, if we have free energy trajectories from a collection of methods A, B, … spanning different ranges of *c*, but there is one method Z for which we have data covering the whole range, we can compute the *relative efficiency* of all methodologies with respect to Z starting from the ratio of the mean errors

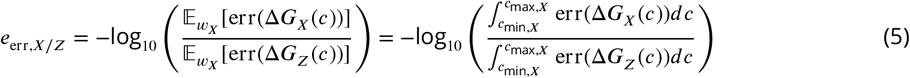

where err is std, bias, or RMSE, X = A, B, …, and the weight function *w*_*X*_ is uniform on the interval [*c*_*min,X*_, *c*_*max,X*_] covered by the data available for method X. The base 10 logarithm ensures *ϵ*_err*,X*/*z*_ = −*e*_err,*z*/*X*_ and facilitates interpretation of the statistic: A relative efficiency *e*_*X*/*z*_ of +1 (−1) means that the total error of X is one order of magnitude smaller (greater) than the total error of Z over the same range of computational cost. We call this the relative *eiciency* of method X as it increases inversely proportional to its mean error. Note that the mean error of Z entering the definition is computed with the same weight function (i.e. over the same interval), which cancels out with the numerator to leave the ratio of the error function areas.

If the methods error decay proportionally to the same function of *c*, the relative efficiency in Eq. (5) is robust to the range of computational cost considered (see Appendix 1 in the supporting information for details). In practice, the statistic seem to be relatively robust to differences in computational cost ranges for most methods (SI Figure 5) with fluctuations that are within the statistical uncertainty of the estimates (SI Figure 6). We thus use the relative efficiency to compare and rank the performance of the methods entering the challenge.

### 2.4 File preparation and information available to participants

The protocol used to prepare the input files is described in the Detailed Methods section. Briefly, for each host-guest system, five different binding poses were selected among the top-scoring predictions of OpenEye’s FRED rigid docking facility [66, 67]. Any docked pose whose guest coordinates had a root mean square deviation (RMSD) less than 0.5 Å with respect to any of the previously accepted docked poses was discarded. This process generated a set of reasonable bound structures with RMSD between any pair of binding poses ranging between 0.72-2.58 Å for CB8-G3 and 1.33-2.01 Å for OA-G3. We then parametrized the systems with AM1-BCC charges [68, 69] and GAFF [70] after solvation in TIP3P [71] water molecules with Na+ and Cl- ions added to neutralize the host-guest net charge and reach a 150 mM ionic strength for CB8 and 60 mM for OA-G3/G6. Finally, we relaxed each replicate with 1 ns of Langevin dynamics to obtain the initial conformations shown in ***Figure 1***. The five conformations of each host-guest pair generally differ both in their positioning within the symmetric binding site and torsion angles. In particular, all rotatable bonds in the guests adopt at least two different dihedral conformations, with the exception of the bonds connecting the carbon in position 4 in OA-G6 to the two methyl groups, and the two carbon-carbon rotatable bonds composing the secondary alcohol linkage connecting the quinoline moiety and the quinuclidine ring of CB8. The input files for different simulation programs were generated and validated with InterMol. Similarly to what was found in [29], the potential energies computed with different packages for the same structures were generally within 1 kJ/mol from each other, except for those computed with AMBER and CHARMM, which differed by about 2–4 kJ/mol from the others. These results were obtained after tampering with the default settings to make the options as similar as possible. Slightly different Coulomb constants are responsible for approximately 70% of the discrepancies, with AMBER and CHARMM adopting values that are furthest away from each other. The remaining 30% is explained by differences in Lennard-Jones cutoff schemes and PME implementations. The contribution from these differences to binding free energy is not trivial predict, but it is expected to be negligible with respect to statistical error and mostly cancel out at the end states of the thermodynamic cycle. The insensitivity to the Coulomb constant definition and PME parameters was confirmed for Hamiltonian replica exchange calculation with the OA-G3 system (see SI Table 1). A detailed breakdown of the energy components in the different packages can be found at https://github.com/samplchallenges/SAMPL6/tree/master/host_guest/SAMPLing. The input files were uploaded to the public GitHub repository together with details on the setup protocol and general instructions about the challenge (https://github.com/samplchallenges/SAMPL6/blob/master/SAMPLing_instructions.md). The instructions also included the recommended values for the simulation parameters known to affect the theoretical binding free energy (e.g., temperature, pressure, Lennard-Jones cutoff, Particle Mesh Ewald settings) in order to minimize factors that could confound the analysis of systematic differences in free energy predictions between methods.

### 2.5 Timeline and organization

Initially, the SAMPL6 SAMPLing Challenge was designed as a blind challenge with deadline Jan 19, 2018. This round included data for the methods referred to below as OpenMM/HREX, GROMACS/EE, OpenMM/SOMD, and OpenMM/REVO. However, OpenMM/SOMD and OpenMM/REVO submissions were affected by two trivial bugs in the calculation setup and the analysis respectively that were corrected after the deadline. Moreover, initial disagreement between OpenMM/HREX and GROMACS/EE, which were originally designated to serve as reference calculations to determine eventual systematic biases arising from methodological issues, prompted us to perform additional calculations. For these reasons, and to further increase the opportunities for learning, we elected to extend the study to more methodologies after the initial results of the calculations were made public and to focus the analysis on the non-blind calculations.

## 3 Results

### 3.1 Overview of free energy methodologies entering the challenge

Seven different free energy methodologies based on alchemical or physical binding pathways and implemented using AMBER [72], GROMACS [73], NAMD [74], or OpenMM [75] entered the challenge. Four of these (referred to in the following as GROMACS/EE, NAMD/BAR, OpenMM/HREX, and OpenMM/SOMD) used the double decoupling methodology [15], and mainly differ in the enhanced sampling strategies and protocols employed. The other three submissions are based on the potential of mean force (AMBER/APR), alchemical nonequilibrium switching (GROMACS/NS-DS/SB), or weighted ensemble (OpenMM/REVO) frameworks. All of the entries computed standard free energies of binding with respect to a standard concentration of 1 M.

In this section, we give a brief overview of the participating free energy methodologies, focusing on their main differences. More details about the methodologies and protocols can be found in Detailed Methods section and in the method description within the submission files available on the public repository at https://github.com/samplchallenges/SAMPL6/tree/master/host_guest/Analysis/Submissions/SAMPLing. Detailed accounts of the results obtained by OpenMM/SOMD and OpenMM/REVO have also been published separately [76, 77] along with detailed accounts of the methodologies they employed.

Importantly, in spite of the focus of this challenge on reproducibility and the best efforts of the organizers and participants, small differences in the model, and thus in the theoretical asymptotic free energy of each method, were introduced in the calculations. This was mostly due to fundamental differences in methodologies and software packages. A brief summary of the main differences affecting the models is included at the end of the section.

#### Double decoupling

The challenge entries with identifier OpenMM/HREX, GROMACS/EE, NAMD/BAR, and OpenMM/SOMD are based on the double decoupling framework[15] for alchemical absolute free energy calculations, which is arguably the most common approach for current absolute alchemical free energy calculations. All three methodologies estimated free energies and their uncertainties using the multistate Bennet acceptance ratio (MBAR) estimator [78] after decorrelating the data, but they differ mainly in the enhanced sampling strategy (or lack thereof) used to collect the data and details of the protocol employed.

OpenMM/HREX used Hamiltonian replica exchange (HREX) [20] to enhance the sampling as implemented in the YANK package [79, 80]. The protocol was based on the thermodynamic cycle in SI Figure 12. Guest charges were annihilated (i.e. intramolecular electrostatic interactions were turned off) before decoupling soft-core Lennard-Jones interactions [81] (i.e. intramolecular interactions were preserved during the alchemical transformation) between host and guest. Since all guests had a net charge, a randomly selected counterion of opposite charge was decoupled with the guest to maintain box neutrality during the alchemical transformation. A harmonic restraint between the centers of mass of host and guest was kept active throughout the calculation to prevent the guest to escape the binding site, and the end-points of the thermodynamic cycles were reweighted to remove the bias introduced by the restraint in the bound state by substituting the harmonic restraint potential to a square well potential. Each iteration of the algorithm was composed of Langevin dynamics augmented by Monte Carlo rigid translation and rotation of the guest and by a Hamiltonian global exchange step (i.e. the exchange was not limited to neighbor states) using the Gibbs sampling approach [82]. The pressure was controlled by a Monte Carlo barostat.

GROMACS/EE employed the weighted expanded ensemble (EE) enhanced sampling strategy [21]. The calculation was performed in the NVT ensemble, and comprised two separate stages, referred to as equilibration and production. During equilibration, the Wang-Landau algorithm [83, 84] was used to adaptively converge to a set of expanded ensemble weights that were then used and kept fixed in the production stage. The data generated using the Wang-Landau algorithm is out-of-equilibrium and non-stationary data, so only the samples generated in the production phase were used for the estimation of the free energy through MBAR, which requires equilibrium samples. The equilibration stage was carried out only for a single replicate, and the same equilibrated weights were used to initialize the other four calculations. We analyzed two separate submissions, identified as GROMACS/EE and GROMACS/EE-fullequil, which differ exclusively in whether the computational cost of the equilibration is “amortized” among the 5 replicas (i.e. the cost is added to each replicate after dividing it by 5) or added fully to each of the 5 replicates respectively. The alchemical protocol uses 20 states to annihilate the electrostatic interactions followed by 20 states to annihilate Lennard-Jones. Two restraints attached to the center of mass of host and guest were used in the complex phase: A flat-bottom restraint, which was kept activated throughout the calculation, and a harmonic restraint that was activated during the annihilation of the Lennard-Jones interactions to rigidify the guest in the decoupled state. The Rocklin charge [63] correction was used to remove the effect of the artifacts introduced by alchemically decoupling a molecule with a net charge. The correction amounted to −0.0219 and −0.0302 kcal/mol for OA-G3 and OA-G6 respectively.

OpenMM/SOMD used the implementation in Sire/OpenMM6.3 [75, 85]. The protocol used 24 intermediate thermodynamic states for CB8-G3 and 21 states for OA-G3/G6 that were simulated independently (i.e. without enhanced sampling methods) with a velocity Verlet integrator and a 2 femtosecond time-step for 20 ns each and a Monte Carlo barostat. Unlike the other submissions, which constrained only bonds involving hydrogen atoms, here all bonds were constrained to their equilibrium values in the host and guest molecules. The temperature was controlled with an Andersen thermostat [86] set at a collision frequency of 10 ps^−1^, and pressure control was achieved with a Monte Carlo Barostat and isotropic box scaling moves were attempted every 25 time steps. In the complex leg of the calculation, a flat-bottom distance restraint between one atom of the guest and four atoms of the host was kept active throughout the calculation. This is the only submission using a generalization of the Barker-Watts reaction field [87, 88] to model long-range electrostatic interactions instead of Particle Mesh Ewald. Reaction field models usually require larger cutoffs to be accurate for relatively large systems due to the assumption that everything beyond the cutoff can be modeled as a uniform dielectric solvent. Consequently, a 12 Å cutoff was used both for Coulomb and Lennard-Jones interactions instead of the 10 Å cutoff employed by the other methods.

Finally, NAMD/BAR calculations were based on the implementation in NAMD 2.12 [74]. In this case as well, the intermediate states were simulated independently with no enhanced sampling strategy and a flat-bottom restraint was used in the complex phase of the calculation. However, 32 *λ* states were used in which the Lennard-Jones interactions were decoupled in equidistant windows between 0 and 1, and the charges were turned off simultaneously over the *λ* values 0–0.9 for CB8-G3 and 0–0.5 for OA-G3 and OA-G6. The second schedule was the result of a protocol optimization to work around an issue in which convergence was impaired by a sodium ion binding tightly the carboxylic group of the OA guests in earlier pilot calculations. A non-interacting particle having the same charge as the guest was created during the annihilation of the Coulomb interactions to maintain the charge neutrality of the box. [65, 89]. The system was propagated with Langevin dynamics using a Nosé–Hoover barostat to control the pressure [65, 89]. Free energy estimates and uncertainties were computed with the BAR estimator.

#### Nonequilibrium alchemical calculations

In GROMACS/NS-DS/SB, the binding free energies were predicted with alchemical nonequilibrium switching calculations using a strategy referred to previously as double-system/single-box [90]. In this approach, two copies of the guest are simulated in the same box, one of which is restrained to the binding site of the host by a set of restraints as described by Boresch [91]. In addition, a harmonic positional restraint is applied to each of the guest molecules to keep them at a distance of 25 Å from one another. The first guest is decoupled simultaneously with the coupling of the second guest in order to keep the net charge of the box neutral during the alchemical transformation. For each replicate, the calculation was carried out first by collecting equilibrium samples from the two endpoints of the transformation. A total of 50 frames were extracted from each equilibrium simulation at an interval of 400 ps, and each snapshot was used to seed a rapid nonequilibrium alchemical transformation of a fixed duration of 500 ps in both directions. For CB8-G3, a second protocol, here referred to as GROMACS/NS-DS/SB-long, was also applied in which 100 snapshots were extracted from each equilibrium simulation at an interval of 200 ps, and each nonequilibrium trajectory had a duration of 2000 ps. Ten independent calculations were run for each of the 5 initial conformations, and a bi-directional estimator BAR, based on Crook’s fluctuation theorem [92], was used to estimate the binding free energy after pooling all work values from all the independent runs. The uncertainty of Δ*G* for each initial conformation was instead estimated by computing the standard error from the ten independent free energy estimates. Because this approach required two copies of the guest and a box large enough to sample distances between host and guest of 25 Å, the complexes were re-solvated. The force field parameters were taken from the challenge input files. However, both with CB8-G3 and OA-G3/G6, the ion concentration was set to 100 mM, which is different than the reference input files. Unfortunately, we realized this after the calculations were already completed.

#### Potential of mean force

AMBER/APR followed the attach-pull-release (APR) [93, 94] methodology to build a potential of mean force profile along a predetermined path of unbinding. The method was implemented in the pAPRika software package based on AMBER [72]. Briefly, the method is divided into three stages. In the “attach” stage, the guest in the binding pocket is gradually rigidified and oriented with respect to the pulling direction in 14 intermediate states through the use of 3 restraints. An additional 46 umbrella sampling windows were used to pull the host and guest apart to a distance of 18 Å. A final semi-analytical correction was applied to compute the cost of releasing the restraints and obtain the binding free energy at standard concentration. The analysis was carried out using thermodynamic integration, and the uncertainties were determined using an approach based on blocking and bootstrap analysis. As in the case of GROMACS/NS-DS/SB, the method required larger solvation boxes than the cubic ones provided by the challenge organizers, in order to reach sufficiently large distances between host and guest. Therefore, the initial five complex conformations were re-solvated in an orthorhombic box, elongated in the pulling direction, of TIP3P waters with Na+ and Cl- ions. The resulting ionic strength differed from the provided files by about 2–5 mM, but the force field parameters were identical.

#### Weighted ensemble of trajectories

The OpenMM/REVO method predicted binding and unbinding kinetic rates with a particular weighted ensemble approach named reweighting of ensembles by variation optimization [77, 95] (REVO) as implemented in the wepy package (https://github.com/ADicksonLab/wepy) using OpenMM [75]. The calculation was carried out by maintaining a set of 48 independent walkers generating MD trajectories starting from bound and unbound states, the latter defined with a distance between host and guest above 10 Å. At each cycle of the algorithm, some of the walkers are cloned or merged in order to maximize a measure of trajectory variation given by the weighted sum of all-to-all distances between walkers. For unbinding trajectories, the distance between two walkers was defined as the RMSD of the system coordinates after aligning the host, while rebinding trajectories used a measure of distance based on the RMSD with respect to the reference unbound starting structure. The *k*_on_ and *k*_off_ rates were estimated directly from the weights of the “reactive” unbinding and rebinding trajectories, and the free energy of binding was computed from the ratio of the rates.

#### Summary of main differences in setups and models

While force field parameters and charges were identical in all calculations, there are small differences among the models used by the different methods. The challenge instructions suggested the settings for simulation parameters that are traditionally not included in parameter files. In particular, most calculations were performed at a temperature and pressure of 298.15 K and 1 atm respectively, using particle mesh Ewald (PME) [96] with a cutoff of 10 Å, and employing a Lennard-Jones cutoff of 10 Å with a switching function between 9 Å and 10 Å. Because of methodological and technical reasons, however, not all simulations were run using these settings. In particular, AMBER does not support switching function so AMBER/APR used a 9 Å truncated cutoff instead, and OpenMM/SOMD supports only reaction field for the treatment of long-range electrostatic interactions. Moreover, even when the suggested settings were used, software packages differ in the supported options and parameter values such as PME mesh spacing and spline order, or the exact functional form of the Lennard-Jones switching function. In addition, all the bonds in OpenMM/SOMD were constrained to their equilibrium value, while all the other calculations constrained only the bonds involving hydrogen. Finally, the APR and NS-DS/SB methodologies required a larger solvated box than the cubic one provided by the organizers. Host and guests were thus re-solvated, and while the force field parameters and charges were preserved, the resulting ion concentrations in the box were slightly different from the original files.

### 3.2 Converged estimates and identical force field parameters do not ensure agreement among methods

#### Absolute free energy calculations can converge to sub-kcal/mol uncertainties in host-guest systems

The final predictions of the submitted methods are shown in ***Table 1***, ***Figure 2***, and SI Figure 7 in terms of the average binding free energy of the five replicate calculations with 95% t-based confidence intervals. With the exception of OpenMM/REVO, the five independent replicate calculations of each method starting from different initial conformations are always within 0.1–0.4 kcal/mol for OA-G3, and 0.1–0.6 kcal/mol for OA-G6 (see also SI Table 3). All methods achieved this level of convergence for the two octa-acid systems in less than 400 · 10^6^ force/energy evaluations (i.e. the equivalent of 800 ns of aggregate MD simulations with a 2 fs integration time step) that can be parallelized over more than 40 processes in all methods with the exception of GROMACS expanded ensemble (see Discussion for more details on parallelization). The agreement between replicates of the same method is generally worse for CB8-G3. Nevertheless, all CB8-G3 predictions of OpenMM/HREX and GROMACS/NS-DS/SB-long are within 0.4 kcal/mol after 2000 · 10^6^ force/energy evaluations (i.e. the equivalent of 4 *μ*s of MD with a 2 fs time step), which suggests that absolute free energy calculations can indeed achieve convergence for this class of systems in reasonable time given widely available computational resources.

**Table 1.**
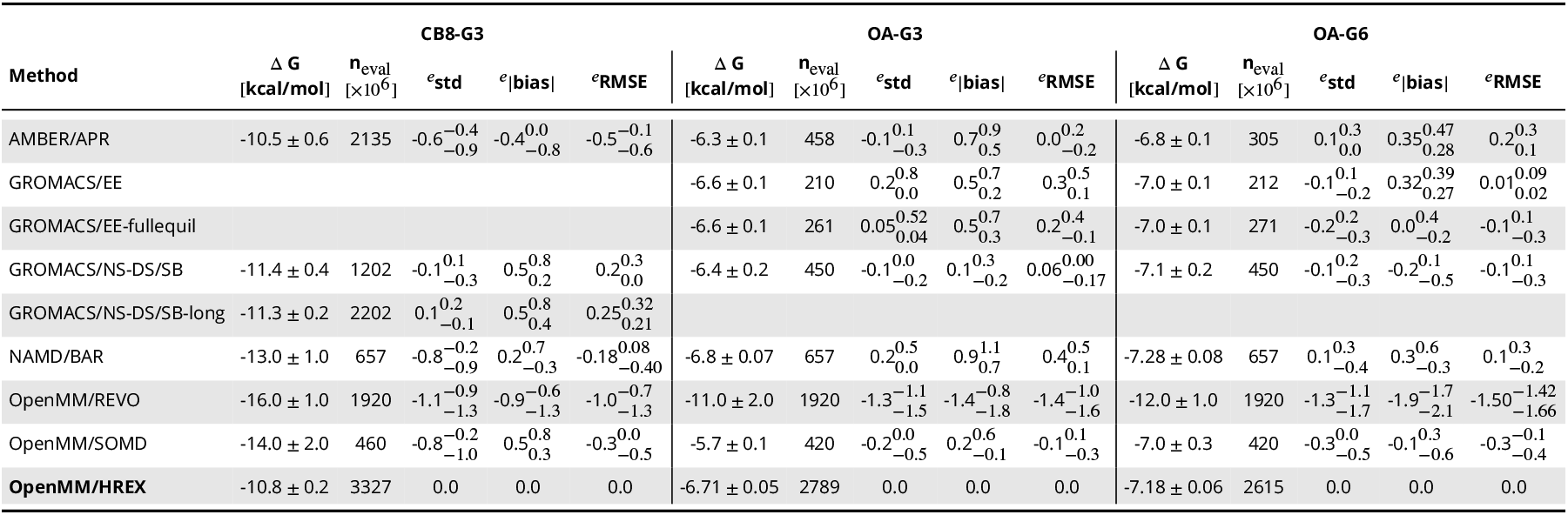
Average binding free energy predictions, computational cost, and relative efficiencies of all methods. Final average binding free energy predictions in kcal/mol computed from the five independent replicate calculations with 95% t-based confidence intervals. The computational cost is reported in millions of force and energy evaluations per replicate calculation. Relative efficiencies of a method *X* are reported with respect to OpenMM/HREX as *e*_err,*X*/OpenMM/HREX_ as defined by Eq. (5). The lower and upper bound of the 95% confidence intervals bootstrap estimates for the relative efficiencies are reported as subscript and superscript respectively.

**Figure 2.**
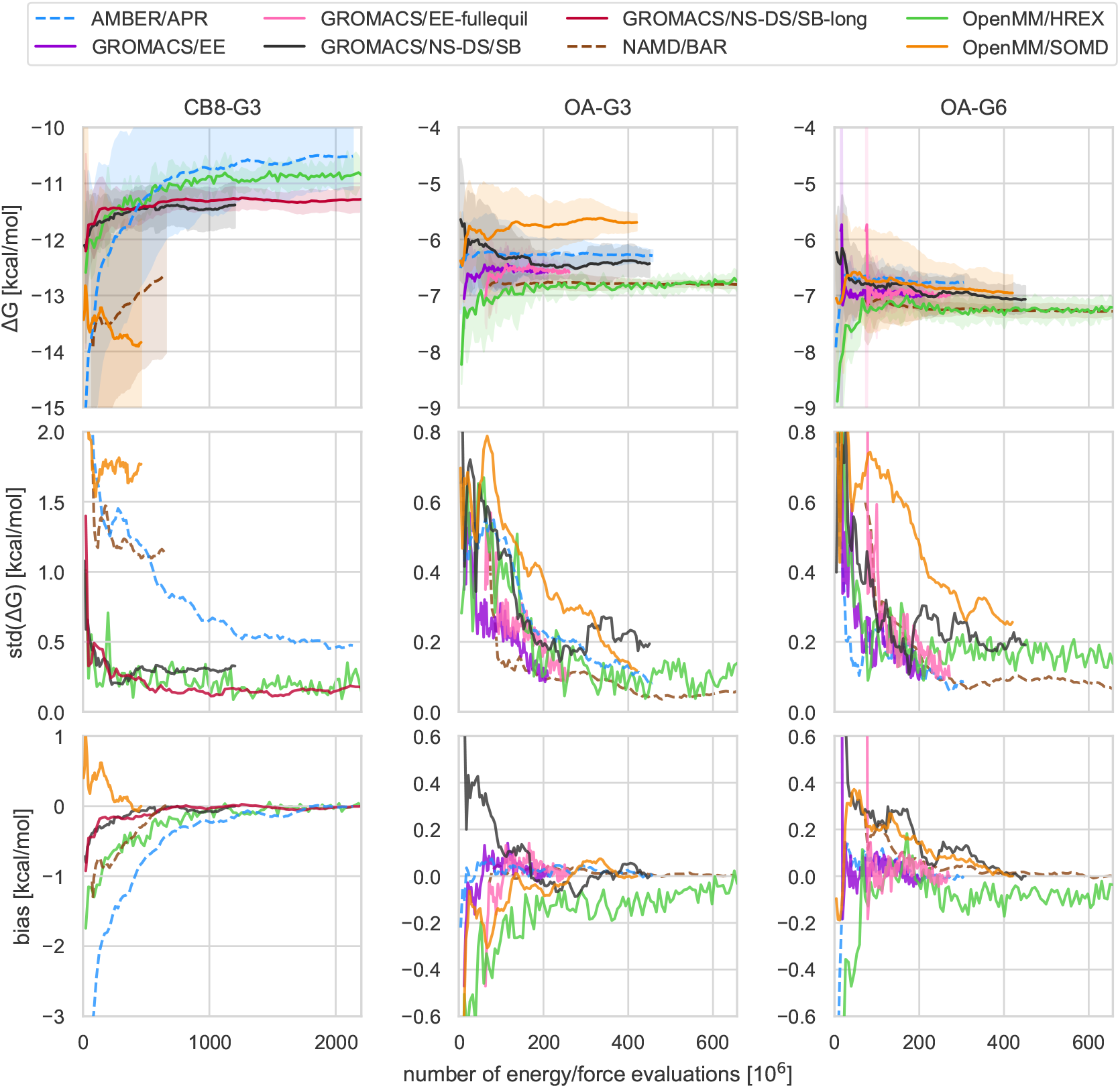
Mean free energy, standard deviation, and bias as a function of computational cost. The trajectories and shaded areas in the top row represent the mean binding free energies and 95% t-based confidence intervals computed from the 5 replicate predictions for CB8-G3 (left column), OA-G3 (center), and OA-G6 (right) for all submissions, excluding OpenMM/REVO. The same plot including OpenMM/REVO can be found in SI Figure 7. The second and third rows show the standard deviation and bias, respectively, as a function of the computational effort. Given the differences in the simulation parameters between different methods, the finite-time bias is estimated assuming the theoretical binding free energy of the calculation to be the final value of its mean free energy. This means that the bias eventually goes to zero, but also that the bias can be underestimated if the simulation is not converged.

#### Identical force field parameters and charges do not guarantee agreement among methods

Although the predictions of different methods are roughly within 1 kcal/mol, the methods sometimes yield statistically distinguishable free energies. For example, OpenMM/REVO tended towards significantly more negative binding free energies than those predicted by the other methods by about 5-6 kcal/mol, and the final predictions of OpenMM/SOMD for OA-G3 were between 0.5 and 1.0 kcal/mol more positive than the other alchemical and PMF methods. NAMD/BAR and OpenMM/SOMD also generally obtained very negative binding free energies for CB8-G3, but in these two cases, the large statistical uncertainty suggests that the calculations are not close to convergence (i.e. the replicate calculations do not agree). This could be a reflection of the smaller number of energy evaluations used for these submissions (see ***Table 1***). AMBER/APR also obtained free energy predictions for OA-G3 and OA-G6 that are significantly different than the predictions from OpenMM/HREX, GROMACS/EE, and NAMD/BAR by 0.2-0.5 kcal/mol. Finally, GROMACS/NS-DS/SB-long and AMBER/APR differ in their predictions for CB8-G3 by 0.8 ± 0.6 kcal/mol.

#### The origin of the discrepancies between free energy predictions is unclear

In several cases, the interpretation of these results is confounded by differences in simulation parameters and setups. For example, without more data, it is impossible to distinguish whether the systematic bias observed in OpenMM/SOMD is due to sampling issues or the use of reaction field instead of PME or a Lennard-Jones cutoff of 12 Å instead of 10 Å. Multiple explanations are also possible for the other observed discrepancies. Firstly, simulation engines generally differ in the implementation details of the long-range treatment strategies. For example, AMBER does not support switched Lennard-Jones cutoff as the AMBER family of force fields was fit with a truncated cutoff. As a consequence, APR calculations were run using a truncated 9 Å cutoff. In principle, the default values and the algorithms used to determine parameters such as the PME grid spacing and error tolerance can also have an impact on the free energies. Secondly, discrepancies may arise from small differences in the model. Specifically, in order to allow for sufficiently great distances between host and guest in the unbound state, the solvation boxes for APR and NS-DS/SB were regenerated and have a slightly different ionic strength, which is known to affect the binding free energy of host-guest systems. Finally, even for these relatively simple systems, differences in sampling, such as those arising from unsurmounted energetic barriers and different numerical integration schemes, could have affected the convergence of the calculations and introduced non-negligible biases respectively.

We investigated most of these hypotheses focusing on APR and HREX, which showed systematic and statistically distinguishable differences of 0.3–0.4 kcal/mol in the final free energies for all systems. The choice of focusing on these two methods was mainly due to technical feasibility as we considered it possible to run further HREX calculations after minimizing the differences in setups and other simulation parameters. However, switching to a truncated 9 Å caused the HREX calculations to increase even further the discrepancies from 0.4 ± 0.1 to 0.7 ± 0.1, while the HREX calculations resulted insensitive to differences in PME parameters, ionic strength, integrator discretization, Coulomb constant, and restraint employed. Detailed results of the sensitivity analysis of HREX can be found in Appendix 2. Although other explanations exist, it is possible that the observed discrepancies between AMBER/APR and OpenMM/HREX are the results of subtle differences or bugs in the software packages, or of an area of relevant configurational space that is systematically undersampled, which was found to be a problem in host-guest systems both with umbrella sampling [97] and alchemical approaches [98]. A version of APR implemented with OpenMM is close to be completed and might prove useful in determining whether the differences are caused by the methods or the simulation package.

Further work will be required to establish the exact source of the persistent deviation between seemingly well-converged calculations.

### 3.3 Bias and variance of free energy estimates can vary greatly with methods and protocols

We estimated standard deviation, bias, and RMSE relative efficiencies for all methods and built bias-corrected and accelerated (BCa) bootstrap [99] 95% confidence intervals (see also Detailed Methods for details). We used the total combined number of force and energy evaluations to measure the computational cost, and OpenMM/HREX was used as a reference for the calculation of the relative efficiencies because it was the longest calculation and could thus provide free energy estimates for all the computational cost intervals required to estimate the statistics. The resulting relative efficiencies with confidence intervals are represented in ***Table 1***.

#### The methods displayed system-dependent performance

Overall, no method emerged as a superior choice in all three systems, but double decoupling, potential of mean force, and nonequilibrium switching all proved to be solid approaches to obtained precise binding free energy estimates for the host-guest systems considered. Indeed, GROMACS/NS-DS/SB (nonequilibrium switching with double-system/single box), NAMD/BAR (double decoupling), and AMBER/APR (potential of mean force) obtained the greatest RMSD efficiency for CB8-G3, OA-G3, and OA-G6 respectively. In general, however, all methods showed larger uncertainty and slower convergence for CB8-G3 than for OA-G3/G6 (Figure 2), and the differences among the methods’ performance, which were relatively small for the two octa-acid systems, increased for CB8-G3. For example, with GROMACS/EE, it was not possible to equilibrate the expanded ensemble weights within the same time used for OA-G3/G6. Moreover, OpenMM/SOMD and NAMD/BAR replicate calculations could not converge the average free energy to uncertainties below 1 kcal/mol, and OpenMM/HREX and AMBER/APR displayed a significant and slowly decaying bias. Contrarily, GROMACS/NS-DS/SB, which generally obtained a slightly negative relative efficiency in OA-G3/G6, performed significantly better than any other methods with CB8-G3 and obtained variance similar to OpenMM/HREX but smaller total bias.

#### Enhanced-sampling strategies can increase convergence rates in systems with long correlation times

The four double decoupling methods performed similarly for the two octa-acid systems, while differences in performance widened with CB8-G3, which featured the largest guest molecule in the set and generally proved to be more challenging for free energy methods than OA-G3/G6. OpenMM/HREX obtained much smaller uncertainties and bias with CB8-G3 than both OpenMM/SOMD and NAMD/BAR, whose replicates seem far from converging to a single prediction. Looking at the individual replicate free energy trajectories for CB8-G3 (SI Figure 9), one notices that both OpenMM/SOMD and NAMD/BAR produced a few relatively flat trajectories that differ by 3-4 kcal/mol. Further OpenMM/SOMD repeats suggest that the replicate disagreement is not determined by the initial conformations, and it is more likely caused by long mixing times of the system (SI Table 5). The difference in performance with respect to OpenMM/HREX for CB8-G3 might then be explained by the Hamiltonian replica exchange strategy, which is in agreement with previous studies on cucurbit[7]uril [100]. On the other hand, NAMD/BAR and GROMACS/EE obtained the greatest relative efficiencies for OA-G3/G6, and, while their difference in efficiency is not statistically significant, it is worth noticing that NAMD/BAR did not employ enhanced sampling methodologies. This suggests that the impact of enhanced sampling strategies based on Hamiltonian exchange might be significant in absolute free energy calculations only for transformations and systems with long correlation times.

Nonequilibrium switching trajectories (the NS protocol) also seemed to be effective in working around problematic energetic barriers in CB8-G3 associated with the alchemical transformation. In particular, NS-DS/SB-long, which used longer nonequilibrium switching trajectories, slightly improved the efficiency of the method in CB8-G3. This suggests that collecting fewer nonequilibrium switching trajectories to achieve a narrower nonequilibrium work distribution can be advantageous in some regimes.

As a final note, NAMD/BAR generally obtained a greater efficiency than OpenMM/SOMD in OA-G3/G6, which also did not use any enhanced sampling approach. It is unclear whether this difference is due to the number of intermediate states (32 for NAMD/BAR, 21 for OpenMM/SOMD), the initial equilibration of 2 ns performed by NAMD/BAR, or the long-range electrostatics model (PME for NAMD/BAR and reaction field for OpenMM/SOMD). It is clear, however, that two different but reasonable protocols can result in very different efficiencies. As a confirmation of this, the NAMD/BAR submission for OA-G3/G6 used an optimized *λ* schedule turning off charges linearly between *λ* values 0.0–0.5 rather than 0.0–0.9 as done in the first batch of calculations. The new *λ* schedule considerably improved the convergence over the original protocol, which was causing long mixing times due to sodium ions binding tightly the carboxylic group of the OA guests.

#### Equilibrating expanded ensemble weights can increase efficiency when running replicates

In the two octa-acid systems, OpenMM/HREX and GROMACS/EE-fullequil achieved similar efficiencies, although the latter obtained a better absolute bias relative efficiency with OA-G3. GROMACS/EE obtained, however, a greater RMSE relative efficiency when the cost of equilibrating the expanded ensemble weights is amortized over the five replicate calculations. This strategy is thus attractive when precise uncertainty estimates through replicate calculations are required. These observations, however, are limited to the two OA systems as the expanded ensemble weights equilibration stage did not converge in sufficient time for CB8-G3. Finally, we note that differences in the details of the protocols between GROMACS/EE and OpenMM/HREX may explain the greater efficiency of the former.

In the expanded ensemble strategy, the weights attempt to bias the probability of jumping from a state to another in order to sample all intermediate states equally. In the presence of bottlenecks, this helps to reduce the round trip time along the alchemical *λ* variable, which in turn can help reducing correlation times of the sampled binding poses in the bound state. Moreover, while OpenMM/HREX decoupled a counterion of opposite charge to the guest to maintain the neutrality of the simulation box, GROMACS/EE corrected for Coulomb finite-size effects arising with PME using an analytical correction [63]. While the approach decoupling the counterion does not introduce approximations, the process of discharging an ion is accompanied by solvent reorganization, which could impact the statistical efficiency of the calculation. Finally, GROMACS/EE annihilated Lennard-Jones (LJ) interactions (i.e. intra-molecular LJ forces were turned off in the decoupled state) while OpenMM/HREX decoupled them (i.e. intra-molecular LJ interactions were left untouched). The choice of decoupling versus annihilating has two effects on convergence, and these may work in opposite directions. On one hand, annihilating the LJ could increase the thermodynamic length of the transformation, which was found to be directly connected to the minimum theoretical variance of the free energy estimate [40]. On the other hand, annihilation of internal LJ interactions might remove some energy barriers separating metastable states, which could help reducing correlation times.

#### Estimating binding free energies via estimation of binding kinetics was an order of magnitude less efficient than predicting binding free energies directly

OpenMM/REVO employed a dramatically different approach for free energy prediction, calculating estimates of the binding kinetics through direct sampling of the binding and unbinding processes. The free energies obtained using the ratio of the binding and unbinding rates had larger uncertainties and showed a significant systematic bias with respect to other methodologies, although the ranking of the compounds agrees with the other submissions. The slow unbinding process may be responsible for the large variance and bias observed in REVO. Indeed, REVO calculations collected a total of 1.92 *μ*s per system per replicate, which should allow obtaining reasonably robust statistics for the binding process, whose mean first passage time (MFPT) estimated by the method for the three systems was between 36±6 and 150±50 ns [77]. On the other hand, the MFPT estimates for the unbinding process yielded by the method were 6±4 *μ*s for OA-G3, 2.1±0.5 s for OA-G6, and 800±200 s for CB8-G3, which is significantly beyond the reach of the data accumulated for the prediction, and suggests that further simulation is required to obtain a better estimate of k_off_ and Δ*G*. Another possible element that may have affected the asymptotic free energies is the size of the simulation box, which was relatively small for this type of calculation and made it difficult to sample long distances between host and guest in the unbound state, which can artificially lower the unbinding rate. Despite the smaller efficiency in predicting the binding free energy, this method was the only one among the submissions capable of providing information on the kinetics of binding.

### 3.4 Unidirectional nonequilibrium work estimators can be heavily biased and statistically unstable

We verified how the choice of the estimator can impact the convergence of the free energy estimate in nonequilibrium switching calculations. In particular, besides the bi-directional BAR estimates discussed above (GROMACS/NS-DS/SB and GROMACS/NS-DS/SB-long), we computed binding free energies of the host-guest systems using uni-directional estimator based on Jarzynski’s equality [103] in both forward and reverse directions and the estimator presented in [102], which is based on Jarzynski’s equality and the assumption of normality of the nonequilibrium work distribution. No extra simulation was run to obtain these new estimates. Rather, the same nonequilibrium data produced by the GROMACS/NS-DS/SB and GROMACS/NS-DS/SB-long protocols were re-analyzed using the unidirectional estimators. Their associated computational cost was halved to account for the fact that the method required to generate only nonequilibrium switching trajectories in one direction. As can be seen in ***Figure 3*** and in SI Table 3, the efficiency of unidirectional estimators is significantly smaller than one obtained with BAR in all cases but GROMACS/NS-Jarz-F for OA-G3, where the sign of the RMSE relative efficiency is not statistically significant. In particular, the estimator based on the Gaussian approximation of the work distribution can be significantly unstable for both the forward (e.g. CB8-G3) and the reverse (e.g. OA-G3) directions. This may be due to the Gaussian estimator’s linear dependency on the work variance, which makes its free energy estimate sensitive to rare events that do not affect Jarzynski’s estimator. For example, the average free energy profile obtained for OA-G3 with the Gaussian estimator in the reverse direction (i.e. Gaussian-Reverse) displays a “saw-like” pattern with large and sudden jumps in the average free energy that are due to single rare events with large work dissipation which substantially increase the variance of the work distribution (SI Figure 10). The work variance subsequently gradually decreases when more regular events are introduced. Moreover, all unidirectional estimates for CB8-G3 are significantly biased, and none of them agree with the bidirectional estimates within statistical uncertainty. In general, this data suggests that collecting nonequilibrium switching trajectories in both directions is worth the cost of generating samples from the equilibrium distributions at both endpoints of the alchemical transformations.

**Figure 3.**
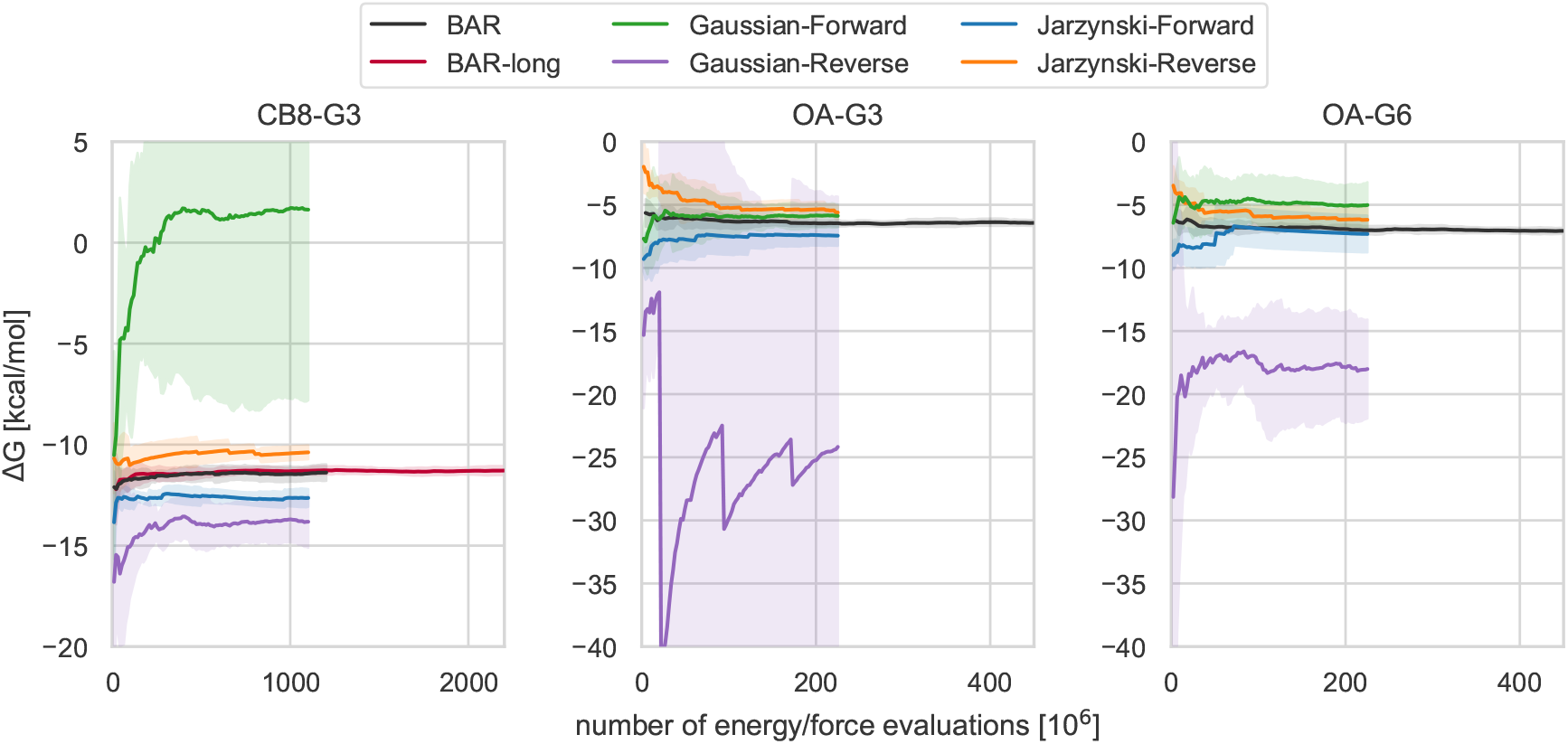
Comparison of bidirectional and unidirectional free energy estimators of the same nonequilibrium work switching data. Average free energy estimates obtained by different estimators from the same nonequilibrium work data collected for CB8-G3 (left), OA-G3 (center), and OA-G6 (right) as a function of the number of energy/force evaluations. The average and the 95% t-based confidence interval (shaded areas) are computed from the 5 replicate calculations. BAR and BAR-long correspond to the GROMACS/NS-DS/SB and GROMACS/NS-DS/SB-long submissions in ***Figure 2***, and utilize the bidirectional Bennett acceptance ratio estimator based on the Crooks fluctuation theorem [101]. Jarzynski-Forward/Reverse are the free energy estimates computed through unidirectional estimators derived from the Jarzynski equality using only the nonequilibrium work values accumulated in the forward/reverse direction respectively. The Gaussian-Forward/Reverse trajectories are based on the Crooks fluctuation theorem and the assumption of normality of the forward/reverse nonequilibrium work distribution, as described in [102]. Unidirectional estimators can introduce significant instabilities and bias in the estimates.

### 3.5 The Berendsen barostat introduces artifacts in expanded ensemble calculations

Initially, the GROMACS/EE free energy calculations were performed in the NPT ensemble, but these converged to different binding free energies than the reference OpenMM/HREX calculations performed with YANK. In order to understand the origin of this discrepancy, we looked into the differences in the protocols adopted by the two methods that could have affected the asymptotic binding free energies. In particular, we examined the robustness of the reweighting step used by YANK at the end points to remove the bias introduced by the harmonic restraint (see also Detailed methods section), the sensitivity of the calculations to the PME parameters (i.e. FFT grid, error tolerance, and spline order), and the barostat employed.

After verifying that the reweighting step and the PME parameters did not impact significantly the free energies predicted by the two methods (SI Figure 2 and SI Table 6), we investigated the effect of the barostat on the asymptotic binding free energy. OpenMM used Metropolis-Hastings Monte Carlo molecular scaling barostat [104, 105] while GROMACS a continuous scaling (or Berendsen) barostat [106]. Because of implementation issues, only the Berendsen barostat was compatible with both expanded ensemble simulations and bond constraints at the time simulations were run. It is known that the Berendsen barostat does not give the correct volume distribution [107, 108], but in most cases, expectations of variables relatively uncorrelated to the volume fluctuations, such as energy derivatives in alchemical variables, might be expected to be essentially unaffected. We thus re-ran both methods in NVT, first with different and then identical PME parameters. If the NVT calculation is run at the average NPT volume, we expect the NVT and NPT binding free energy predictions to be essentially identical as, in the thermodynamic limit, d*G* = d*A* + d(*pV*), where *G* and *A* are the Gibbs (NPT) and Helmholtz (NVT) free energies respectively, and we expect 1 atm·Δ*V̅*, where *V̅* is the change in volume on binding, to be negligible. The box vectors used for the NVT calculations were selected from the OpenMM/HREX NPT trajectories in order to obtain the volume closest to the average NPT volume. The changes introduced by the different PME parameters were not statistically significant (SI Table 6), but we found that the discrepancies between the methods vanished without the barostats. In particular, OpenMM/HREX yielded free energies identical to those obtained at NPT, whereas the expanded ensemble predictions for OA-G3 decreased by 0.6 kcal/mol, suggesting that the Berendsen barostat was responsible for generating artifacts in the simulation.

To obtain further insight, we performed molecular dynamics simulations of OA-G3 at 1 atm and 100 atm in NPT using the GROMACS Berendsen barostat and the OpenMM Monte Carlo barostat. We found that the Berendsen barostat generated volume distributions with much smaller fluctuations and slightly different means than the MC barostat. At 1 atm, the mean of the Berendsen and MC barostat distributions are 80.250 ± 0.006 nm^3^ and 80.286 ± 0.004 nm^3^ respectively (errors here are two times the standard error of the mean). In contrast to the MC barostat, reweighting the distribution generated by the Berendsen barostat at 1 atm with the weight *e*^*β*(100atm−1atm)*V*^ fails to recover the 100 atm distribution (***Figure 4***), which confirms that the Berendsen barostat did not sample correctly the expected volume fluctuations in the NPT ensemble. Moreover, the volume distribution sampled in the bound state by the Berendsen barostat during the expanded ensemble calculations is quite different from that obtained through simple MD simulations, with thicker right tails and mean 80.298 ± 0.008 nm^3^. The apparent shift to the right is consistent with the volume expansion observed in the neighbor intermediate states during the expanded ensemble calculations (SI Figure 8), which suggests that the artifacts might be introduced by the random walk along states. In principle, we expect the difference in binding free energy due to the different barostats to be approximately *p*(Δ*V̅* _MC_ − Δ*V̅* _B_), where Δ*V̅* _MC/B_ is the change in volume on binding from according to the MC or Berendsen barostat, as indicated. However, because the mean volume for the Berendsen and MC barostats are different even for the simple MD simulation, it is not completely clear whether a difference in free energy would still be present without the expanded ensemble algorithm. In fact, the mean bound state volume obtained by the Berendsen barostat during the expanded ensemble calculation is closer to the MC mean volume than the one obtained with MD. Further free energy calculations using the Berendsen barostat but independent *λ* windows might be helpful in clarifying this issue.

**Figure 4.**
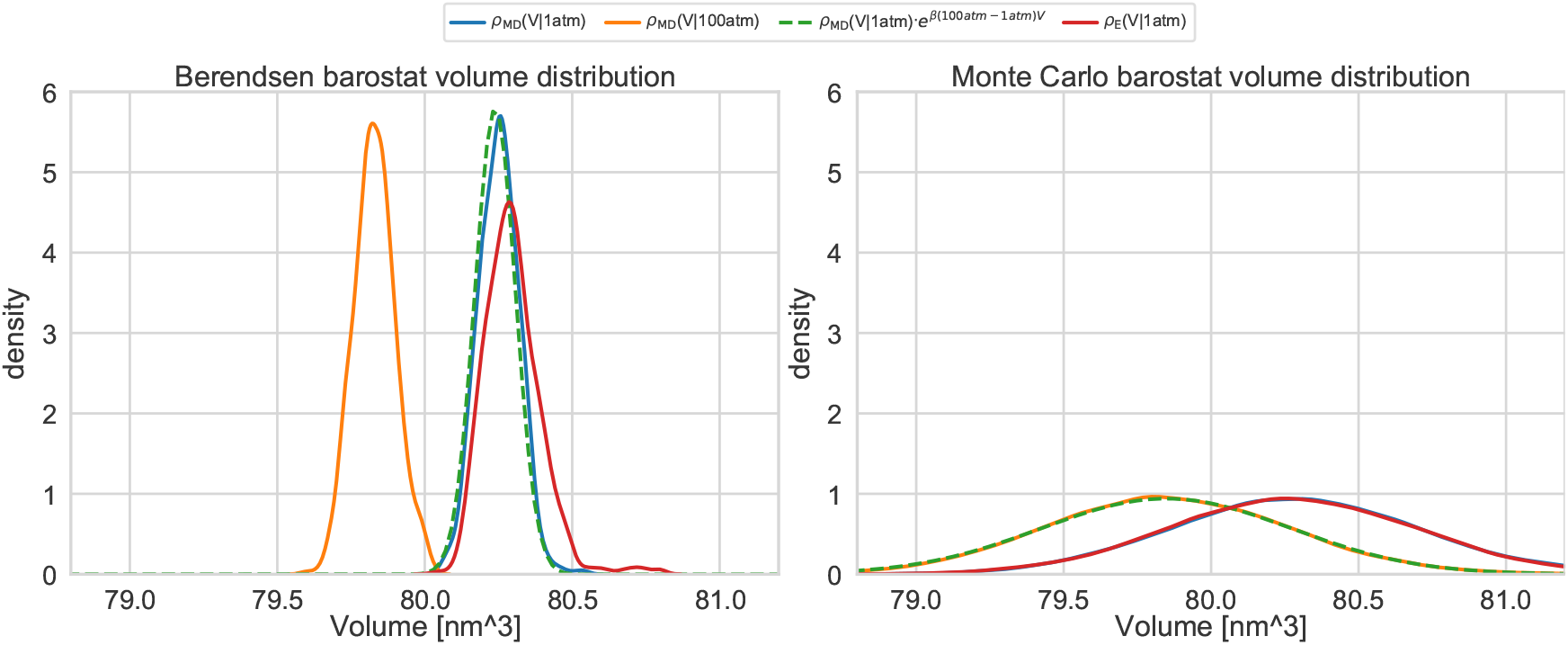
OA-G3 volume distribution, restraint radius distributions, and binding free energy dependency on the binding site definition. Box volume empirical distributions obtained by NPT simulations using the Monte Carlo barostat implemented in OpenMM (right) and the Berendsen barostat implemented in GROMACS (left) at 298 K. The continuous blue (*ρ*_MD_(V|1atm)) and orange (*ρ*_MD_(V|100atm)) lines represent Gaussian kernel density estimates of volume distributions sampled with simple molecular dynamics at a constant pressure of 1 atm and 100 atm respectively. The green distribution is obtained by reweighting *ρ*_MD_(V|1 atm) to 100 atm. The red densities (*ρ*_MD_(V|1 atm)) represent the volume distribution sampled in the bound state by the enhanced sampling algorithm (i.e., expanded ensemble for the Berendsen barostat and HREX for the Monte Carlo barostat). The expected distribution is predicted correctly only from the volumes sampled using the Monte Carlo barostat, while the Berendsen barostat samples distributions of similar mean but much smaller fluctuations. Moreover, the expanded ensemble algorithm introduce artifacts in the volumes sampled by the Berendsen barostat.

### 3.6 Estimators of the free energy variance based on correlation analysis can underestimate the uncertainty

Since participants also submitted uncertainty estimates for each of the five replicate calculations, we were able to verify how accurately the different uncertainty estimators could reproduce the true standard deviation of the Δ*G* estimates, here referred to as std(Δ*G*), from a single run. OpenMM/HREX, GROMACS/EE, and SOMD estimated the single-replicate uncertainties from the asymptotic variance estimator of MBAR after decorrelating the potential based on estimates of the integrated autocorrelation time. AMBER/APR instead used blocking analysis to compute the mean and standard error of dU/d*λ* in each window. These statistics were then used to generate 1000 bootstrapped splines, and the uncertainty was determined by computing the standard deviation of the free energies from the thermodynamic integration of the bootstrapped splines. Finally, GROMACS/NS-DS/SB estimated the uncertainties by running an ensemble of 10 independent non-equilibrium switching calculations for each of the 5 replicate calculations and computing their standard deviations. We built *ŝ*(Δ*G*), our best estimate of std(Δ*G*), with 95% confidence intervals for each method by computing the standard deviation of the five replicated free energy predictions. Under the assumption of normally-distributed Δ*G*, *ŝ*(Δ*G*) is distributed according to *ŝ*(Δ*G*) ~ *χ*_*N*−1_std(Δ*G*)/(*N* − 1), where *N* = 5 is the number of replicates [109], which makes it trivial to build confidence intervals around *ŝ*(Δ*G*).

Under this statistical analysis, the single-replicate trajectories of most methods are within the confidence interval of *ŝ*(Δ*G*) (SI Figure 9). In particular, the standard deviations of the single GROMACS/NS-DS/SB replicate calculations generally agree within statistical uncertainty to our best estimate. This is probably expected as both are based on independent calculations. The AMBER/APR uncertainty estimates based on bootstrapping also agree well with the replicate-based estimate, especially in the final part of the trajectory. We note, however, that the MBAR standard deviation estimate based on autocorrelation analysis statistically underestimates *ŝ*(Δ*G*) in OpenMM/SOMD, and, in general, it shows a marked tendency to be on the lower end of the confidence interval also in OpenMM/HREX and GROMACS/EE. These observations are consistent with those of a prior comparison of the autocorrelation and blocking analysis methods [94]. Similarly, the BAR standard deviation in the NAMD/BAR submission did well for the two octa acids, but the uncertainty was significantly underestimated for the CB8-G3, in which the true standard deviation was on the order of 1.2 kcal/mol. Curiously, the MBAR uncertainties are almost identical across the five replicates in all three submissions using them and for all systems. This is in contrast not only to bootstrap- and replicate-based methods but also to the BAR uncertainty estimates submitted by NAMD/BAR, which seem to yield estimates that are more sensitive to differences in the single free energy trajectories.

In order to verify if the performance of the MBAR uncertainties was due to an inadequate decorrelation of the samples, we analyzed again the HREX data after raising the interval used for subsampling from approximately 2.8 ps to 5, 10, 20, 50, 100 and 200 ps. In this case, the equilibration time, and thus the number of initial iterations discarded, was determined as two times the statistical inefficiency. As SI Figure 11 shows, setting the statistical inefficiency to 5 ps is sufficient for the single-replicate uncertainty to fall within the best estimate confidence interval, and arguably, the agreement becomes slightly better with greater values of statistical inefficiency. However, the single-replicate uncertainties are still almost identical across the five replicates even for the estimates obtained with statistical inefficiency set at 200 ps, in which, due to the limited number of samples, the individual free energy trajectories are quite different and show very different errors. Thus, while the error computed through autocorrelation analysis is within statistical uncertainty of the standard deviation, the estimates seem insensitive to the particular realization of the free energy trajectory.

### 3.7 The initial bias of HREX is explained by the starting population of the replicas

#### The initial conformation can bias the free energy in systems with long correlation times

In all three host-guest systems, we noticed that the OpenMM/HREX free energy trajectories were significantly biased at the beginning of the calculation. The problem was particularly evident for the CB8-G3 system, for which the performance of methods was generally poorer, and a lot of computational effort was required for the bias to decay in comparison to OA-G3 and OA-G6. ***Figure 5*** shows that the initial bias of CB8-G3 gradually disappears when an increasing amount of data from the initial portion of the calculation is ignored during the analysis. This suggests the initial conditions to be the cause of the bias. This becomes apparent when realizing that the HREX free energy trajectory in ***Figure 5*** observed after discarding 2000 iterations can be interpreted as from HREX calculations starting from different initial conditions. What is peculiar about this equilibration process is the consistent sign of the observed bias(i.e. 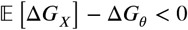), which remains negative even after several thousands iterations are removed (1000 iterations corresponding to the equivalent of 131 ns of aggregate simulation from all replicas). The same trend is observed both for OA-G3 and OA-G6, although the correlation times governing the equilibration process appear much smaller in these two cases than with CB8-G3.

**Figure 5.**
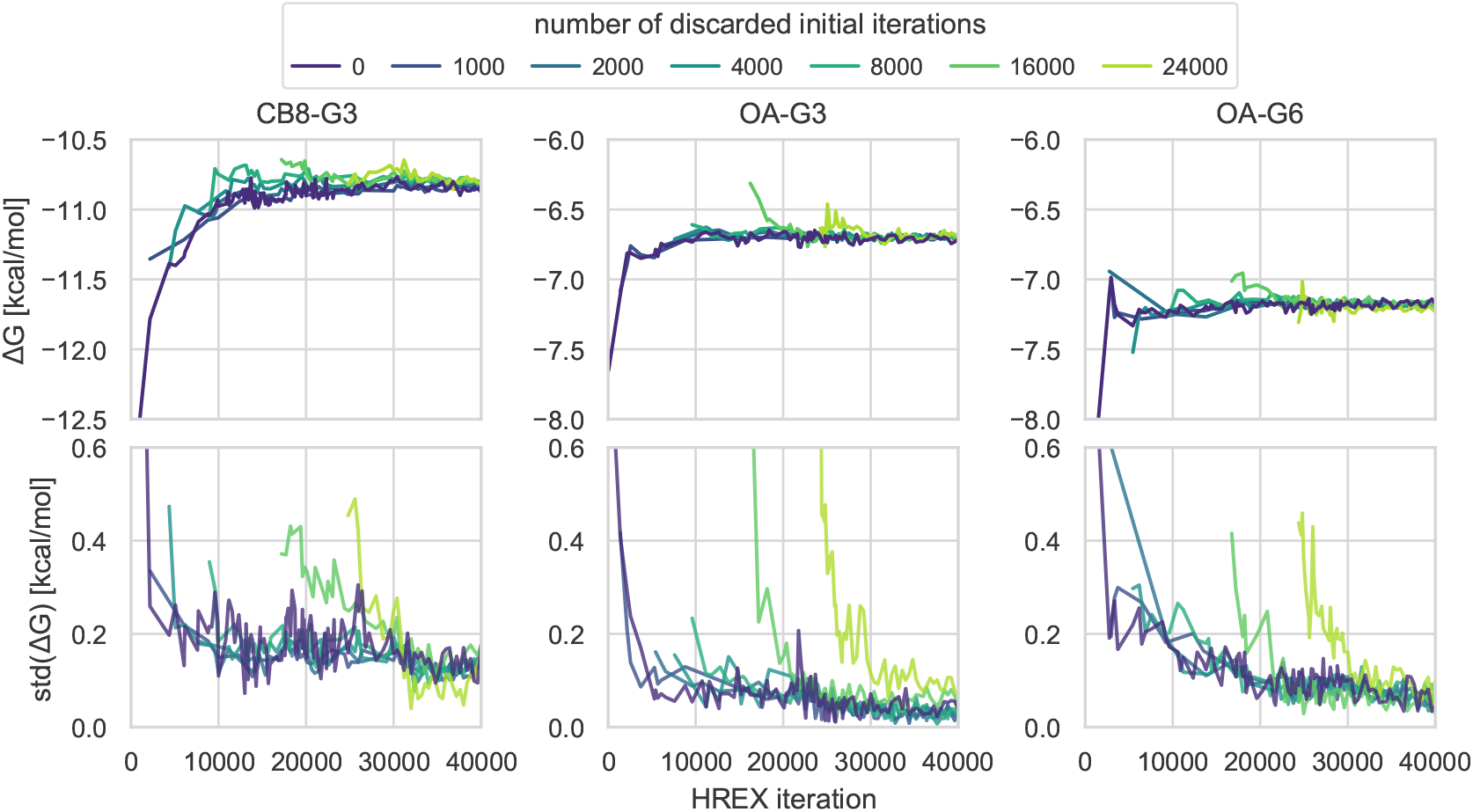
Initiating the HREX calculation from a single conformation introduces significant bias that slowly relaxes as the system reaches equilibrium. Mean (top row) and standard deviation (bottom row) of the five replicate free energy trajectories as a function of the simulation length computed after discarding an increasing number of initial iterations going from 1000 (purple) to 24000 (light green) for the three host-guest systems. The trajectories are plotted starting from the last discarded iteration. The initial bias is consistently negative, and it decays faster in OA-G3/G6 than in CB8-G3, in which correlation times are longer. Ignoring the beginning of the trajectory removes the bias.

#### Initializing all replicas with a bound structure might be the cause of the negative sign of the bias

Decomposing the free energy in terms of contributions from complex and solvent legs of the HREX calculation shows that the finite-time bias is entirely attributable to the complex phase (SI Figure 13). As it is common to do with multiple-replica methodologies, all HREX replicas were seeded with the same initial conformation, which, for the complex phase, was obtained by equilibrating the docked structures for 1 ns in the bound state.

The so-obtained initial structure is representative of the bound state, and we expect it to decorrelate quickly in the decoupled state thanks to the missing steric barriers and the Monte Carlo rotations and translations performed by YANK. On the other hand, the intermediate states might require a long time to relax the initial conformation, during which the generated samples will be closer to the bound state distribution than if they had been sampled from the intermediate states equilibrium distribution. Under these conditions, the free energy estimator will predict the bound state to have a lower negative free energy. A detailed explanation of this last fact can be found in Appendix 3 in the supporting information.

An alternative explanation for the negative sign of the bias relies on the increase in entropy that often accompany the transformation from the bound to the decoupled state. This is usually attributed to the larger phase space available to receptor and ligand and to solvent reorganization [110], and, in this instance, it is confirmed by the entropy/enthalpy decomposition of the predicted free energy (SI Figure 14). The hypothesis relies on the assumption that the larger phase space available in the decoupled state would require thorough sampling to be estimated correctly, which would be impossible at the beginning of the calculation when the estimate would be computed from a small number of correlated samples. As a result, the difference in entropy between the end states would initially be underestimated, and the binding free energy would become more positive as the number of samples enables a more precise prediction. However, this hypothesis seems unlikely, at least in this case, as it does not explain why ignoring the initial part of the calculation would result in an unbiased estimate since the beginning of the free energy trajectory would still be based on an equivalently small number of samples. The large fluctuations of the estimated entropy and potential energy trajectories, which are in the range of 10-20 kcal/mol (SI Figure 14) against a bias of less than 2 kcal/mol, hinder the direct verification of the two hypotheses, but further investigation of the cause and sistematicity of the negative bias across different receptor-ligand systems is currently ongoing.

#### Relevance for other methods

While, for reason of data availability, we focused on HREX here, it should be noted that, in principle, this is not a problem confined to the HREX methodology, and most free energy trajectories generated by alchemical methods show an initial upward trend in all three host-guest systems that may be due to one of these two explanations. In fact, the bias of HREX in CB8-G3 seems to decay faster than other multiple-replica double decoupling methods (i.e., NAMD/BAR and OpenMM/SOMD), whose free energy estimates are still significantly more negative when compared to more converged estimates (e.g., APR, HREX, NS-DS/SB) at the same computational cost (***Figure 2***). This is consistent with our hypothesis as the enhanced sampling strategy should help reducing correlation times of the intermediate states as well. Indeed, while we could not identify a specific physical collective variable responsible for the slow decorrelation of the intermediate states, the correlation time of the replica state index is consistent with the bias decay time in CB8-G3 and OA-G3/G6 (***Figure 6***).

**Figure 6.**
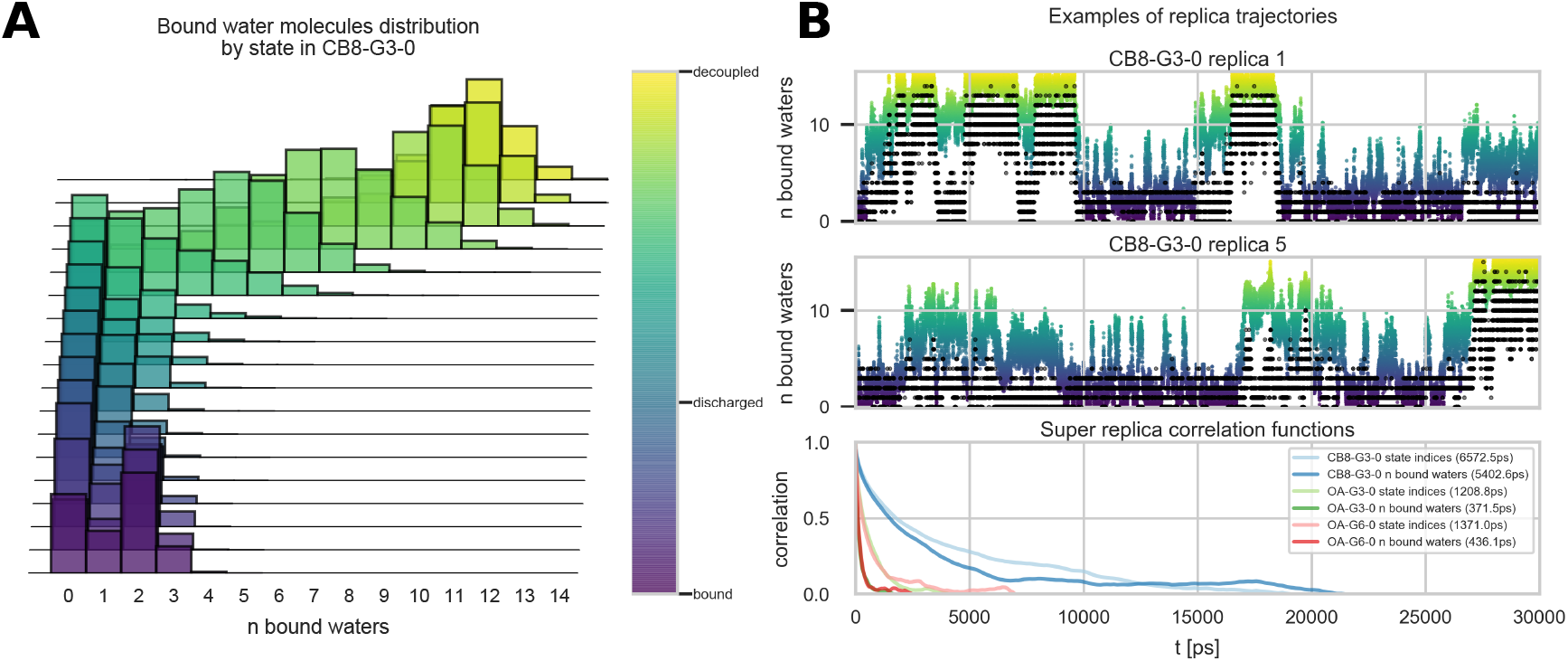
Bound water molecules induce metastability in HREX replicas with CB8-G3. **(A)** Histograms of the number of bound water by thermodynamic state. The color maps the progression of the alchemical protocol from the bound state (purple) to the discharged state (blue), where all the charges are turned off but Lennard-Jones interactions are still active, and decoupled state (yellow). The number of bound waters has a peaked distribution around 0-2 for most of the alchemical protocol, and it rapidly shifts to the right in the near-decoupled state. **(B)** Superposition of the trajectories of the number of bound waters and the state index for replica 1 and 5 of the OpenMM/HREX calculation for CB8-G3-0 (top) and autocorrelation function computed from the time series of the number of bound waters (dark colors) and replica state indices (light colors) for CB8-G3-0 (blue), OA-G3-0 (green), and OA-G6-0 (red) (bottom). Each autocorrelation function was computed as the average of the correlation functions estimated for each replica trajectory [39, 111]. Replicas remain stuck in the near-decoupled states for several nanoseconds. CB8-G3 exhibits much longer correlation times for both time series than the two OA systems.

The data suggest that cheap methods for the determination of sensible initial conformations for the intermediate states may improve considerably the efficiency of HREX in systems with long correlation times. Moreover, a better trade-off between bias and variance in the final estimate could be achieved with better strategies for automatic equilibration detection or by reducing the number of intermediate states (69 for the complex and 62 for the solvent in the CB8-G3 HREX calculations), which directly impact the total number of energy evaluations spent equilibrating the replicas.

### 3.8 Water binding/unbinding in CB8-G3 might contribute to long correlation times in HREX

In order to get insights into the origin of the large uncertainties generally obtained by the double decoupling submissions for the CB8-G3 system, we analyzed the correlation times of various collective variables (CV) in the complex phase of the OpenMM/HREX calculations. ***Figure 6*** shows that the number of waters in the binding site of CB8-G3 is metastable and correlate with the state index of the replicas (where each replica of the Hamiltonian replica exchange calculation can explore multiple states). The number of bound waters was computed by counting the water molecules with at least one atom within the convex hull of the heavy atoms of CB8. The metastability along replica trajectories depicted in ***Figure 6B*** is connected to a rapid shift towards greater numbers of the distribution of bound waters near the decoupled state (***Figure 6A***). This contrasts with the discharging step, where the only evident change is a change in the mode of the bound water histogram from 2 to 0. The shift in mode is consistent with the observed distribution of restrained distance between host and guest (SI Figure 2), which suggests that the guest tends to crawl into the hydrophobic binding site in the discharged state to compensate for the loss of the polar interactions with water. Histograms of the number of bound waters for OA-G3 and OA-G6 (SI Figure 16) show similar features to that of CB8-G3, but the mean number of bound waters in the decoupled state is smaller (i.e. 4.84 water molecules) due to the smaller volume of the octa-acid binding site. Moreover, the statistical inefficiency computed from the correlation function of the state index, which was previously found to correlate well with the uncertainty of free energy estimates in Harmiltonian replica exchange calculations [39], is about five times smaller for OA-G3/G6 (1208.8 ps and 1371.0 ps) than for CB-G3 (6572.3 ps). This is consistent with the slower convergence generally observed for the latter set of calculations.

While these results prove only the existence of correlation between the metastabilities in the number of bound waters and the state indices along a replica trajectory in the CB8-G3 calculations, it is plausible to hypothesize that water molecules displaced by the quinine when the Lennard-Jones interactions are re-coupled, alongside eventual steric clashes with the host binding site, might contribute significantly to hindering the replica exchange step with obvious negative effects on the ability of the HREX algorithm to enhance sampling. This is consistent with the faster replica exchange mixing observed for OA-G3/G6 as coupling the guest would have to displace a smaller number of bound waters than CB8-G3 due to the smaller volume of the guests. No other CV we analyzed had statistical inefficiencies on the same order of magnitude as those observed for the bias decay time shown in ***Figure 5***. In particular, both the host-guest distance restrained by the harmonic potential and the distance between the alchemically-decoupled counterion and the guest seem to decorrelate quickly along replica trajectories, with estimated statistical inefficiencies never exceeding 50 ps. Possibly, an increased number of intermediate states close to the decoupled state might enhance the replica exchange acceptance rates for CB8-G3 and reduce the statistical inefficiency of the state index.

### 3.9 Methods generally overestimated the host-guest binding free energies with respect to experimental measurements

Accuracy with respect to experiments was not the focus of this study, but the input files for the challenge were created using a quite typical setup, and it is thus interesting to compare the converged predictions to the corresponding experimental data collected for the accuracy host-guest challenge [26, 112, 113]. The ITC measurements yielded binding free energies of −6.45 +−0.06 kcal/mol for CB8-G3, −5.18 +−0.02 kcal/mol for OA-G3, and −4.97 +−0.02 for OA-G6. In comparison, the well-converged computational results were more negative on average by −4.4, −1.2, and −2.1 kcal/mol respectively, in line with what was observed for other methods employing the GAFF force field in the SAMPL6 host-guest accuracy challenge [26]. It should be noted that the ionic strengths of SAMPLing systems (i.e., 150 mM for CB8-G3 and 60 mM for OA-G3/G6) were slightly higher than in experimental conditions (estimated to be 57.8 mM for CB8-G3 and 41.25 mM for OA-G3/G6) used for the host-guest binding challenge, and previous evidence revealed the host-guest binding free energies to be sensitive to concentration and composition of the ions. In a recent SOMD calculations performed for the SAMPL6 accuracy challenge, removing the ions modeling ionic strength of the experimental buffer (i.e. going from 150 mM for CB8-G3 and 60 mM OA-G3/G6 to 0 mM) caused the Δ*G* prediction to shift by −4.87 ± 2.42, 1.37 ± 0.50, and 1.48 ± 0.48 for CB8-G3, OA-G3, and OA-G6 respectively (computed as the average of three runs ± standard error of the mean) [76]. In particular, the estimated binding free energy for OA-G3 obtained without buffer ions agreed with the experimental measurement within uncertainty. It is unlikely for the ion concentrations to be the sole responsible for the overestimated binding affinities. The sign of the shift for CB8-G3 described above is not consistent with the hypothesis, and a negative mean error was very consistent across GAFF submissions employing different buffer models. Nevertheless, the order of magnitude of these shifts suggests that ionic strengths cannot be neglected.

## 4 Discussion

### 4.1 Disagreements between methodologies impact force field development and evaluation

In many cases, methods obtained statistically indistinguishable predictions with very high precision. The agreement between methodologies is quite good for OA-G6, where essentially all estimates are within 0.4 kcal/mol. On the other hand, despite the focus of the study on reproducibility, some of the methods yielded predictions that significantly deviated from each other by about 0.3 to 1.0 kcal/mol. This directly raises a problem with force field evaluation and development since it implies that the accuracy afforded by a given set of forcefield parameters (and thus the value of the loss function used for their training) can, in practice, be affected significantly by the software package, methodological choices, and/or details of simulation that are considered to have negligible impact on the predictions (e.g., switched vs truncated cutoff, treatment of long-range interactions, ion concentrations). Trivially, this also implies that we should not expect a force field to maintain its accuracy when using simulation settings that differ from those used during fitting.

Similar observations were made in previous work in different contexts. In a reproducibility study involving four different implementations of relative hydration free energy calculations, the authors found in many cases statistically significant ΔΔ*G* differences on the order of 0.2 kcal/mol [30]. Systematic differences of the same order of magnitude were detected in a recent study comparing Monte Carlo and Molecular Dynamics sampling for binding free energy calculations [27], although, in this case, differences in water models and periodic boundary conditions might confound the analysis.

### 4.2 Bias is critical when comparing the efficiency of different methodologies

The results show that quantifying not only the variance but also the bias of a binding free energy method is important to draw a complete picture of the efficiency of a method. The bias of the free energy predictions varied substantially depending on the method and the system, and for calculations that are short with respect to the correlation times, the bias can be greater or have the same order of magnitude of the variance. For example, in CB8-G3, NS-DS/SB-long obtained a greater RMSE efficiency than HREX in spite of the similar variance because the bias of OpenMM/HREX for CB8-G3 remained non-negligible for a substantial portion of the calculation. This suggests that looking at the variance of the free energy estimate alone is insufficient to capture the efficiency of a method, and the RMSE relative to the asymptotic binding free energy prediction should be favored as the main statistic used in studies focusing on exploring and testing methodological improvements.

Estimating the RMSE and bias is a more complicated problem than estimating the variance as it requires the value of asymptotic free energy given by the model and thus to ascertain that the calculation has converged. Visual inspection of the free energy trajectory is useful, but it can be misleading. Besides the presence of unexplored relevant areas of configurational space, the noise in the trajectory can hide very slow decays (see YANK calculation in CB8-G3). More recommendations about how to detect convergence issues can be found in [114, 115].

On the other hand, a focus on quantifying the efficiency of free energy calculations in terms of RMSE could increase the attention paid to convergence issues as well as incentivize the creation of reference datasets that could provide asymptotic free energies associated to specific input files without always requiring long and expensive calculations. The latter would particularly benefit the field when the efficiency of a method would need to be evaluated only for very short protocols (e.g. overnight predictions). This is, however, conditional on identifying the source of the discrepancies between the predictions of different methods and an asymptotic value can be agreed upon in the first place.

### 4.3 Multiple replicates are one route to avoiding underestimating the uncertainty

MBAR uncertainties and bootstrap uncertainties built with the blocking method were in most cases able to estimate the standard deviation of the free energy prediction within confidence interval. Nevertheless, when sampling is governed by rare events and systematically misses relevant areas of conformational space, data from a single trajectory simply cannot contain sufficient information to estimate the uncertainty accurately. An example is given by the CB8-G3 calculations performed by OpenMM/SOMD and NAMD/BAR, for which the uncertainty estimates were underestimated by more than 1 kcal/mol. In these cases, replicate calculations starting from independent conformations can offer a solution to or compensate for the problem. Relaxed docked conformations can be a viable method to generate the independent conformations, although this is not, in general, an easy task and multiple short replicates starting from the same or very similar initial conformations can still cause the uncertainty to be underestimated. Moreover, given a limited amount of computational resources, the number of replicate calculations should not be large enough to prevent sampling of all the relevant time scales, which are strongly system-dependent.

In addition to a more accurate estimate of the free energy estimate, it has been argued that predictions computed from an ensemble of independent calculations lead to more robust estimates [32, 116]. In agreement with these results, the simple average of the five independent free energies is surprisingly robust even when the single-replicate predictions do not agree quite well (SI Figures 9,11).

### 4.4 Shortcomings of the analysis and lesson learned for future studies

#### The bias estimation strategy favors short and unconverged calculations

Originally, the calculations run by the organizers (i.e., OpenMM/HREX and GROMACS/EE) were meant to provide a reference estimate of the asymptotic free energy of the model that we could use to detect and estimate systematic biases. However, because of the differences in setups and treatment of long-range interactions adopted in the different submissions, this type of analysis was not possible. Instead, we estimated the asymptotic free energy for each methodology as the average binding free energy of the 5 replicates after 100% of the computational cost. As a consequence, the bias is generally underestimated, and long calculations and converged results are thus generally penalized in the calculation of the efficiency statistic. Some of these differences could be minimized by picking settings to which most software packages and methods will be able to adhere. For example, providing systems solvated in both cubic and elongated orthorhombic boxes, and running reference calculations for both of them, could lower the barrier for PMF calculations to enter the challenge without re-solvating the reference files. Moreover, using a truncated cutoff instead of a switched cutoff could help as AMBER does not support switched cutoffs and different simulation packages could use slightly different switching functions. Also, providing template input configuration files for common simulation packages that encapsulate other settings such as PME parameters could reduce the risk of running several methods with different settings.

#### The number of force evaluations can miss important information about the computational cost

In this work, we have focused the analysis on the number of energy/force evaluations as a measure of the methods’ computational cost. In general, this is a very practical and fair measure of the cost of a method. For example, unlike wall-clock or CPU time, it does not depend on hardware and the particular implementation, which is compatible with the objective of this challenge in detecting fundamental differences in efficiency between algorithms. Thus, even though implementation details might affect wall-clock/GPU time dramatically, methods with a comparable number of energy/force evaluations might eventually be able to be put on equal footing given enough developer time if it seemed warranted. Moreover, this measure treats both molecular dynamics and Monte Carlo strategies equally, which would not be possible if the cost was measured, for example, in terms of simulation time (e.g., nanoseconds of simulation).

However, the number of force/energy evaluations can miss important details. It is insensitive to the system size, and it assumes that the computational cost of all other components of the calculation is negligible. Furthermore, while some sampling schemes require multiple evaluations of the Hamiltonian, often it is not necessary to compute it in its entirety. For example, in multiple time scale MD and Monte Carlo moves involving a reduced number of degrees of freedom, one only needs to compute a subset of pairwise interactions. HREX requires the evaluation of multiple Hamiltonian at the same coordinates, but only the parts of the Hamiltonian that change between intermediate state needs to be evaluated multiple times. When the algorithms and setups differ, this may become important to take into account. For example, double decoupling methods assigned the same computational cost to each time step of the complex and solvent stages of the calculation, while REVO, APR, and NS-DS/SB ran only in one stage using a box of the same or greater size of the complex so that one force evaluation for the latter methods on average is practically more expensive than a force evaluation for double decoupling.

In future challenges, it might be useful to collect another simple but more precise measure of the computational cost of a method based on a scaled version of the number of energy/force evaluations, with the scaling factor depending on the number of particles that enters the evaluation. Moreover, instead of requesting exactly 100 free energy estimates for each replicate, requesting free energy estimates that are roughly equally spaced by a predetermined number of force/energy evaluations could make it simpler to perform direct comparisons between all methods without requiring the comparison to a reference calculation.

#### A larger and more varied test set is necessary to obtain a more comprehensive picture of the methods’ efficiency

This first round of the challenge was created as a component of the SAMPL6 host-guest challenge, and we created a minimal test set including both fragment-like and drug-like compounds. We believe this was a beneficial decision. Fragment-like guests that converged relatively quickly such as OA-G3/G6 proved very useful to debug systematic differences between methods while most of the methods problems or strengths were unveiled from the calculations targeting CB8-G3, which has a greater size and generally proved to be more challenging for free energy methods than the two octa-acid guests.

Expanding the test set to include one trivial system and a few more challenging systems could increase the potential for learning and provide a more complete picture of the problems to address and the domain of applicability of the different methods, especially as different approaches may have different strengths and weaknesses. For example, HREX and EE could be less effective at improving convergence for systems with a single dominant binding mode. On the other hand, systems with a buried binding pocket that remains dry in both the holo and apo states could be less problematic for HREX and EE, which are challenged by wetting/dewetting processes that occur at the “almost decoupled” state. At the same time, physical-pathway methods such as APR and REVO might be less effective for receptor-ligand systems with buried binding pockets, as an efficient unbinding path could require large reorganization of the receptor that might be difficult to determine or sample.

For systems that are easier to converge, it might also be possible to increase the number of replicates from five. The increased statistical power could be particularly helpful to resolve differences between methods in efficiency, in estimated binding free energy predictions, and for the analysis of the uncertainty estimates (e.g. blocking, bootstrap, and correlation analysis) since the standard deviation of the binding free energy estimated from five replicates have large variance, which makes it hard to draw statistically significant conclusions. For bigger systems, this may not be practical, but the number of replicates does not necessarily have to be the same for all the tested systems.

Finally, we point out that the selection of systems for such convergence studies is not limited by the lack of experimental data or a chemical synthesis route, and one is free to craft an optimal test system.

### 4.5 Parallelization considerations

The analysis above does not account for the differences in the intrinsic levels of parallelization of the different methods, but almost all methods can be completely or almost trivially parallelized over up to 40 parallel processing units with the given protocols. APR, NAMB/BAR, and SOMD protocols use respectively 60, 64, and 40 windows, the last two numbers to be divided equally between complex and solvent stages. Each HREX calculation ran more than 100 MC/MD parallel simulations, although the exchange step provides a bottleneck for the simulation. Similarly, the protocol used for the REVO methodology employs 48 independent walkers that can be run in parallel throughout the calculation, with a bottleneck occurring at the cloning/merging stage of the adaptive algorithm. NS-DS/SB protocol used 10 independent equilibrium simulations for each end state (i.e. bound and unbound states) that generate frames used to spawn nonequilibrium switching trajectories in both directions. New NS trajectories can be started as soon as new equilibrium samples are generated. Thus, because the nonequilibrium trajectory duration in this protocol is greater than the interval between two equilibrium frame, the calculation can in principle have at least 40 independent simulations running in parallel. The EE protocol submitted for this work is an exception as it does not use a parallelization scheme, although maintaining and coordinating multiple independent expanded ensemble chains is in principle possible [117].

Nevertheless, all calculations can also be trivially parallelized over the molecules in the set and over eventual independent replicate calculations. Under perfect parallelization, or in the presence of negligible bottlenecks, the relative efficiency is insensitive to the number of parallel processing units so we expect the analysis in this work can be informative also in many common scenarios involving parallel computing systems. However, these results should be careful re-interpreted in the presence of massively parallel computational systems, in which the number of processing units does not provide a fundamental bottleneck. For example, a large number of GPUs could be exploited better with protocols simulating many intermediate states that can be simulated in parallel, such as those used by HREX and APR.

### 4.6 Relevance and future plans for relative free energy calculations

Unfortunately, we did not receive any relative free energy submission for this round of the challenge. However, the data reported here has implications for relative calculations as well. Given that enhanced sampling strategies based on Hamiltonian exchange had little or no impact on efficiency for the octa-acid systems, we expect a relative calculation to be significantly more efficient than two absolute calculations in computing a ΔΔ*G* value for the simple OA-G3 to OA-G6 transformation that we set up. For the same reason, we would expect enhanced and non-enhanced relative methods to perform similarly for the OA-G3 to OA-G6 transformation. On the contrary, a relative transformation involving a system with long correlation times such as CB8-G3 might benefit more from enhanced sampling strategies and be less sensitive to the initial bound conformation. Finally, while cancellation of error might help, we expect to observe discrepancies between different packages and/or methods also for relative calculations as, with the exception of OpenMM/HREX and AMBER/APR, the ΔΔ*G* between methods does not appear to be systematic.

In future rounds of the challenge, we are interested in probing the boundaries of applicability of this technology, particularly in the presence of ligands or alchemical transformations requiring exploration of multiple, kinetically-separated binding modes. For these cases, state-of-the-art methods in both absolute and relative calculations often rely on scaling selected Coulomb, Lennard-Jones, and/or torsional terms of the Hamiltonian to lower the energetic barriers between relevant conformations [35, 118] Typically, the optimal choice for the range of scaling factors and the subset of the systems to enhance is very system-dependent, not known *a priori*, and essentially determined by a trade-off between shortening mixing times and simulating extra intermediate states sharing poor overlap with the end states of the transformation. In this sense, absolute methods such as HREX and EE bring this trade-off to an extreme by turning off completely receptor-ligand electrostatics and sterics interactions. This enables dramatic changes in binding pose, such as the upside-down flip of CB8-G3, at the cost of introducing states with poor overlap with the end states, although without usually modifying torsions or receptor atoms that would reduce the overlap even further. Again, careful selection of the receptor-ligand systems will be fundamental to determine under which conditions protocols favoring sampling or statistical efficiency would result in faster convergence.

## 5 Conclusions

We have presented the results of the first round of the SAMPLing challenge from the SAMPL challenge series. The design and execution of the challenge made apparent the need for a measure of efficiency for free energy calculations capable of capturing both bias and uncertainty of the finite-length free energy estimates and summarizing the performance of a method over a range of computational costs. The analysis framework and efficiency statistics we introduced in this work allow formulation and evaluation of hypotheses regarding the efficiency of free energy methods that can be verified meaningfully with the standard tools of statistical inference. We applied this framework to seven free energy methodologies and compared their efficiency and their level of agreement on a set of three host-guest systems parametrized by the same force field. The analysis highlighted significant and system-dependent differences in the methods’ convergence properties that depend on both the sampling strategies and the free energy estimator used. Overall, the study shows that PMF and alchemical absolute binding free energy calculations can converge within reasonable computing time for this type of system.

Surprisingly, we observed significant differences in the converged free energies for the different methods ranging from 0.3 to 1.0 kcal/mol. These discrepancies are small enough that they would not have aroused suspicion without the comparison of multiple independent methods, which stresses the utility and efficacy of this type of study in detecting methodological problems. While we were able to isolate the origins of some of these discrepancies, further work will be required to track down the causes of remaining discrepancies, which might be attributable to small differences in the model (e.g. treatment of long-range interactions, ionic strength), sampling issues of some of the methods, software package, or any combination of the above. Notably, the discrepancies between methods are roughly half the size of the current reported inaccuracies of leading free energy methods compared to experiment (roughly 1 kcal/mol). Eliminating these discrepancies would therefore be very useful for the field to make further progress.

Although we decided to accept non-blinded submissions to increase the value of the study, future rounds of the challenge should ideally be limited to blind predictions, in line with the other challenges within the SAMPL series. The lessons learned while organizing this first round of the challenge will be useful to address the problems identified during the analysis. In particular, we hope to adopt a slightly different measure of computational cost based on the number of force/energy evaluations that also takes into account the system size, and increase the size and variety of the test set. Although an aspirational goal, running on the same dedicated hardware would allow a meaningful comparison of the performance of the different methods also in terms of CPU/GPU time, and analyze more closely the speedups obtained with parallelization. Workflow-ized tools (e.g., Orion workflows, BioSimSpace workflows, HTBAC) could be helpful in pursuing this direction.

## 6 Detailed methods

### 6.1 Preparation of coordinates and parameters files

The protonation states of host and guest molecules were determined by Epik 4.0013 [119, 120] from the Schrödinger Suite 2017-2 at pH 7.4 for CB8-G3 and pH 11.7 for OA-G3 and OA-G6. These values correspond to the pH of the buffer adopted for the experimental measurements performed for the SAMPL6 host-guest binding affinity challenge. For each host-guest system, 5 docked complexes were generated with rigid docking using FRED [66, 67] in the OpenEye Toolkit 2017.6.1. Binding poses with a root mean square deviation less than 0.5 Å with respect to any of the previously generated binding poses were discarded. Hosts and guests were parameterized with GAFF v1.8 [70] and antechamber [121]. AM1-BCC [68, 69] charges were generated using OpenEye’s QUACPAC toolkit through OpenMolTools 0.8.1. The systems were solvated in a 12 Åbuffer of TIP3P [71] water molecules using tleap in AmberTools16 [122] shipped with ambermini 16.16.0. In order to make relative free energy calculations between OA-G3 and OA-G6 possible, ParmEd 2.7.3 was used to remove some of the molecules from the OA systems and reduce the solvation box to the same number of waters. This step was not performed for the CB8-G3 system, and the 5 replicate calculations where simulated in boxes containing a different number of waters. The systems’ net charge was neutralized with Na+ and Cl- ions using Joung-Cheatham parameters [123]. More Na+ and Cl- ions were added to reach the ionic strength of 60 mM for OA-G3/G6 systems and 150 mM for CB8. Note that this ionic strength is likely to be different from the one used for the experimental measurements, which was estimated to be 41 mM and 58 mM respectively. Systems were minimized with the L-BFGS optimization algorithm and equilibrated by running 1 ns of Langevin dynamics (BAOAB splitting [22], 1 fs time step) at 298.15 K with a Monte Carlo barostat set at 1 atm using OpenMM 7.1.1 [75] and OpenMMTools [124]. Particle Mesh Ewald (PME) was used for long-range electrostatic interactions with a cutoff of 10 Å. Lennard-Jones interactions used the same 10 Å cutoff and a switching function with a switching distance of 9 Å. After the equilibration, the systems were serialized into the OpenMM XML format. The rst7 file was generated during the equilibration using the RestartReporter object in the parmed.openmm module (ParmEd 2.7.3). The AMBER prmtop and rst7 files were then converted to PDB format by MDTraj 1.9.1 [125]. The files were converted to GROMACS, CHARMM, LAMMPS, and DESMOND using InterMol [29] (Git hash f691465, May 24,2017) and ParmEd (Git hash 0bab490, Dec 11, 2017).

### 6.2 Free energy methodologies

#### AMBER/APR

We used the attach-pull-release (APR) [93, 94] method to calculate absolute binding free energies of each host-guest complex. We used 14 “attach” umbrella sampling windows, during which time host-guest complex restraints are gradually applied, and 46 “pull” umbrella sampling windows to separate the host and guest. A final, analytic “release” phase was applied to adjust the effective guest concentration to standard conditions (1 M). Since CB8 has two symmetrically equivalent openings, and the APR method only pulls the guest out of one opening, we have added an additional −RT ln(2) = −0.41 kcal/mol to the calculated binding free energy to adjust for this additional equivalent entropic state.

The restraints were setup using our in-development Python package: pAPRika 0.0.3 (commit hash e69f053). Six restraints (1 distance, 2 angles, and 3 dihedrals) were used to restrain the translational and orientational degrees of freedom of the host relative to three positionally restrained dummy anchor atoms. These restraints, which were constant throughout all APR windows, did not perturb the internal degrees of freedom of the host. The distance force constant was set to 5.0 kcal/mol-Å^2^ and the angle force constant to 100.0 kcal/mol-rad^2^. Three additional restraints were added, during the attach phase of APR, between the dummy atoms and two guest atoms in order to orient the guest relative to the host and then separate the two molecules by 18 Å, which was sufficient for reaching a plateau in the potential of mean force. The distance and angle force constants for these restraints were the same as before.

All equilibration and production simulations were carried out with the GPU-capable pmemd.cuda MD engine in the AMBER 18 package [72]. The OA systems were re-solvated with 3000 waters and the CB8 systems were re-solvated with 2500 waters in a orthorhombic box elongated in the pulling direction to enable distances between the host and guest necessary to carry out the potential of mean force calculation. Force field parameters and charges of the host-guest systems were not altered in the operation. Equilibration consisted of 500 steps of energy minimization and enough NPT simulation such that 1 ns could be completed without the simulation box dimensions changing beyond AMBER limits (up to 10 ns total). All simulations used a time step of 2 fs, with a Langevin thermostat and a Monte Carlo barostat. The nonbonded cutoff was set to 9.0 Å, and the default AMBER PME parameters were employed.

For the OA-G3 simulations, we performed 10 ns of sampling per window. For the OA-G6 simulations, we performed 15 ns of sampling per window. For the CB8-G3 simulations, we performed 70 ns of sampling per window. In all cases, we used thermodynamic integration to compute the binding free energies. To compute the uncertainties, we used blocking analysis to calculate the mean and standard error of *dU* /*dλ* in each window, where *U* is the potential energy and *λ* is the reaction coordinate. We then created 1000 bootstrapped splines through points sampled off the distribution determined by the *dU* /*dλ* mean and standard error of the mean for each window, used trapezoidal integration for the total free energy for each spline, and computed the mean and standard deviation of the free energies from the bootstrap samples.

#### GROMACS/NS-DS/SB and GROMACS/NS-DS/SB-long

The estimates were obtained with alchemical nonequilibrium free energy calculations using GROMACS 2018.3 [73] as described in [90]. Briefly, both legs of the thermodynamic cycle were carried out in the same box: i.e. one guest molecule was decoupled from the solvent while another copy was coupled while in the host binding pocket. The two guest molecules were placed 2.5 nm apart and restrained with a single position restraint on one of their heavy atoms. For the guest molecule bound to the host, a set of restraints as described by Boresch [91] (1 distance, 2 angles, 3 dihedrals) was applied. A force constants of 10 kcal/mol-Å^2^ was applied to the distance, and constants of 10 kcal/mol-rad^2^ were applied to the angles.

First, both end-states (A: bound guest coupled and unrestrained, unbound guest decoupled; B: bound guest decoupled and restrained, unbound guest coupled) were simulated using 10 simulations of 20 ns each (20.2 ns for CB8), for a total of 400 ns of equilibrium sampling (404 ns for CB8). Each of these 20 simulation boxes had been previously built from the input files provided by the organizer by re-solvating the host-guest systems and randomly placing ions in the box at a concentration of 0.1 M, followed by minimization with 10000 steps of steepest descent. The re-solvation was a necessary step to enable sufficient distance between the host and guest in the unbound state and did not alter the force field parameters of hosts and guests. However, differently from the challenge input files, Cl- and Na+ ions were added to the simulation to reach a 100 mM concentration.

For the OA systems, 50 frames were extracted from each of the equilibrium simulations at an interval of 400 ps. Thus, in total 500 frames were extracted from the equilibrium simulations of each of the two end-states. For the CB8 systems, 100 frames were extracted from each of the equilibrium simulations every 200 ps, for a total of 1000 frames. The extracted snapshots were used to spawn rapid nonequilibrium alchemical transitions between the end-states. In the nonequilibrium trajectories, the Hamiltonian between the two end states was constructed by linear interpolation.

The alchemical transitions were performed in both directions (A->B and B->A) in 500 ps per simulation for the OA systems, and in 1000 ps for the CB8 systems. A second submission identified by GROMACS/NS-DS/SB-long used a 2000 ps nonequilibrium trajectory instead and only for CB8-G3. For the unbound guest, charges were annihilated (i.e. intra-molecular electrostatics was turned off) and Lennard-Jones interactions were decoupled (i.e. intra-molecular sterics was left untouched) at the same time, using a soft-core potential for both. The same protocol was used for the bound guest except that also the Boresch restraints were switched on/off during the nonequilibrium transitions by linearly scaling the force constants. The two positional restraints attached to the two copies of the guest were left activated throughout the calculation. All simulations used Langevin dynamics with a 2 fs time step with constrained hydrogen bonds. Periodic boundary conditions and Particle Mesh Ewald were employed with a cutoff of 10 Å, interpolation order of 5, and tolerance of 10^−4^. A cutoff of 10 Å with a switching function between 9 Å and 10 Å was used for the Lennard-Jones interactions. An analytical dispersion correction for energy and pressure was also used to account for the dispersion energy. The Langevin thermostat was set at 298.15 K and a Parrinello-Rahman barostat [126] was employed to maintain the pressure at 1 atm.

The binding free energy was estimated with pmx [127] from the set of nonequilibrium work with the BAR [128, 129] estimator after pooling all the data from the ten independent calculations. Uncertainties were instead estimated by computing the standard error of the ten individual BAR estimates.

#### GROMACS/EE and GROMACS/EE-fullequil

The free energy of bindings were obtained with the double decoupling method [15] using the expanded ensemble enhanced-sampling methodology [21] implemented in GROMACS 2018.3 [73]. Charges were turned off completely before removing Van der Waals interactions in both the complex and the solvent phase. Both Coulomb and Lennard-Jones interactions were annihilated (i.e. intra-molecular interactions were turned off). Two restraints were used during the complex phase of the calculation: a flat-bottom restraint with radius 1.5 nm and spring constant 1000 kJ/mol-nm^2^, and a harmonic restraint with spring constant 1000 kJ/mol-nm^2^. Both restraints were attached to the centers of mass of host and guest, but while the flat-bottom restraint remained throughout the simulation, the harmonic restraint was incrementally activated while the Lennard-Jones interactions were removed. In the bound state, the flat-bottom distance between the centers of mass remained always smaller than the 1.5 nm radius necessary to have a non-zero potential.

Because of instabilities and bias introduced by the Berendsen barostat during the expanded ensemble calculation, all the simulations were performed in NVT using the average volume sampled by the OpenMM/HREX calculations performed with YANK. V-rescale temperature was used to keep the temperature at 298.15 K, and and bonds to hydrogen atoms were constrained using the SHAKE algorithm. We used the md-vv integrator, a velocity Verlet integrator, with time steps of 2 fs. Metropolized Gibbs Monte Carlo moves between all intermediate states [82] were performed every 100 time steps based on weights calculated with the Wang–Landau (WL) algorithm as described below. The metropolized Gibbs move in state space proposes jumps to all states except the current state, with a rejection step to satisfy detailed balance. An equal number of time steps were allocated to production simulations of complex and solvent systems for each free energy estimate. A cutoff of 10Å was used for nonbonded interactions with a switching function between 9 Å and 10 Å for Lennard-Jones forces. Particle Mesh Ewald used an interpolation order of 5 and a tolerance of 10^−5^. A sample .mdp file can be found in the submission at https://github.com/samplchallenges/SAMPL6/blob/master/host_guest/Analysis/Submissions/SAMPLing/NB006-975-absolute-EENVT-1.txt.

The expanded ensemble calculation was divided into two stages: an equilibration stage, in which the expanded ensemble weights were adaptively estimated, and a production stage that generated the data used to compute the submitted free energy estimates and in which the weights were kept fixed. In the equilibration stage, the weights are adaptively estimated using the Wang-Landau algorithm [83, 84]. For all systems an absolute value of the initial Wang–Landau incrementor was set to 2 k_*B*_ T. Weights were updated at each step, and the increment amount was reduced by a factor of 0.8 each time a flat histogram was observed, meaning that the ratio between the least visited and most visited states since the last change in the weight increment was less than 0.7. The process of updating the weights was halted when the incrementing amount fell below 0.001 k_*B*_ T. Equilibration of the weights was only ran on a single starting conformation out of five for each host-guest pair. The weight of the fully coupled state is normalized to zero, meaning that the weight of the uncoupled state corresponds to the free energy of the process. The last stage of the simulation, during which period the expanded ensemble weights were no longer updated, was termed the “production” stage since it was the only part of the trajectory used to calculate the final free energy change. Once the Wang–Landau incrementor reached a value of 0.001 k_*B*_ T the simulation was stopped, MBAR was ran on simulation data obtained while the Wang–Landau incrementor was between values of 0.01 and 0.001 k_*B*_ T, and the resulting free energies were used to set the weights for the production simulations for all starting conformation of a host-guest pair.

Reported values were obtained by running MBAR on production simulation data. The submissions GROMACS/EE and GROMACS/EE-fullequil differ only in whether the computational cost of the equilibration is added in its entirety to each of the five replicate calculations (GROMACS/EE-fullequil) or whether it is ammortized over the replicates (GROMACS/EE).

#### NAMD/BAR

The alchemical free energy calculations were performed using the double decoupling method as implemented in NAMD 2.12 [74]. The NAMD protocol utilized a total number of 32 equidistant *λ* windows, that are simulated independently for 20 ns/window with Langevin dynamics using a 2 fs time step and coupling coefficient of 1.0 ps^−1^. The Lennard-Jones interactions are linearly decoupled from the simulation in equidistant windows between 0 and 1, while the charges were turned off together with LJ over the *λ* values 0-0.9 for CB8-G3 and 0-0.5 for OA-G3 and OA-G6. During the complex leg of the simulation a flat-bottom restraint with a wall constant of 100 kcal/mol/Å^2^ was applied to prevent the guest from drifting away from the host. A non-interacting particle having the same charge of the guest was created during the annihilation of the Coulomb interactions in order to maintain the charge neutrality of the box [65, 89]. Before collecting samples for the free energy estimation, each window was equilibrated for 2 ns. The pressure was maintained at 1 atm using a modified Nosé–Hoover method implemented in NAMD, in which Langevin dynamics is used to control fluctuations in the barostat [130, 131]. The Langevin piston utilized an oscillation period of 100 fs and a damping time scale of 50 fs. Long range electrostatic interactions were treated with the following PME parameters: PME tolerance = 10^−6^, PME spline order 4, and PME grid = 48×48×48. The cutoff for both Lennard-Jones and PME was set to 10 Å, and the switching distance was set to 9 Å. The free energy of each replicate calculation and their uncertainties were computed with BAR using ParseFEP [132] Tcl plugin (version 2.1) for VMD 1.9.4a29.

#### OpenMM/HREX

The free energy calculations and analysis were performed with YANK 0.20.1 [79, 80] and OpenMMTools 0.14.0 [124] powered by OpenMM 7.2.0 [75]. The protocol followed the double decoupling methodology [15] using the thermodynamic cycle in SI Figure 4. In both phases, we first annihilated the guest charges (i.e. intra-molecular electrostatics was turned off) and then decoupled the soft-core (1-1-6 model) Lennard Jones interactions [81] (i.e. intra-molecular sterics was left untouched). The spacing and number of intermediate states was determined automatically for the three systems by the trailblaze algorithm implemented in YANK [79]. This resulted in a protocol with a total of 69 and 62 intermediate states for the complex and solvent phase respectively of CB8-G3, 59 and 54 states for OA-G3, and 55 and 52 states for OA-G6. Since all guests had a net charge, a counterion of opposite charge was decoupled with the guest to maintain the box neutrality at each intermediate state and avoid artifacts introduced by finite-size effects with Particle Mesh Ewald.

Hamiltonian replica exchange [20] was used to enhance sampling of the binding modes. Each iteration of the algorithm was composed by a metropolized rigid translation, using a Gaussian proposal of mean 0 and standard deviation 1 nm, and a random rotation of the ligand followed by 1 ps of Langevin dynamics (BAOAB splitting [22], 2 fs timestep, 10/ps collision rate). A Monte Carlo barostat step was performed every 25 integration steps to maintain a pressure of 1 atm. All hydrogen bonds were constrained. The Hamiltonian exchange step was carried out after each iteration by performing *K*^4^ metropolized Gibbs sampling steps [82], where *K* is the number of intermediate states in the protocol. At the beginning of each iteration, velocities for all replicas were randomly re-sampled from the Boltzmann distribution. In all calculations, we ran 40000 iterations of the algorithm (i.e. 40 ns of MD per replica) for both the complex and solvent calculation for a total MD propagation of 5.24 *μ*s, 4.52 *μ*s, and 4.28 *μ*s for each of the five replicates of CB8-G3, OA-G3, and OA-G6 respectively. An analytical dispersion correction for the long-range Lennard-Jones interactions was added during the simulation for all atoms except the alchemically-softened atoms for optimization reason. The contribution of the guest to the dispersion correction was instead found by reweighting the end states.

The analysis of the samples was performed with the MBAR estimator [78] with PyMBAR 3.0.3. We computed an estimate of the statistical inefficiency of the sampling process in order to decorrelate the HREX samples. The statistical inefficiency was estimated from the correlation function of the time series of the traces of the *K* × *K* MBAR energy matrix *U* (*i*) computed at each iteration *i*, where the matrix element *U_jl_* (*i*) is the reduced potential of the sample generated by state *j* at iteration *i* and evaluated in state *l*. The resulting statistical inefficiencies were 2.74 ± 0.03 ps, 2.9 ± 0.3 ps, and 2.84 ± 0.3 ps for CB8-G3, OA-G3, and OA-G6 respectively (uncertainties are given as the standard deviation of the statistical inefficiencies over replicates). The statistical inefficiency was then used to discard the burn-in data by maximizing the number of effective samples as described in [133] and to subsample the data before running MBAR. In the complex phase, the guest was restrained throughout the calculation into the binding site through a single harmonic restraint connecting the center of mass of the heavy atoms of host and guest with a spring constant of 0.2 kcal/(mol · Å^2^) for CB8-G3 and 0.17 (mol · Å^2^) for OA-G3/G6. Following the double decoupling approach, an analytical correction was added to bring the affinity in units of standard concentration and correct for the restraint volume in the decoupled state. However, because the restraint was activated in the bound state as well, we also used MBAR to reweight the samples to remove the bias introduced by the harmonic potential. Samples whose restrained distance (i.e. the distance between the host and guest centers of mass) was above a specific threshold were discarded. This is equivalent to reweighting the data to a state having a restraint following a square well potential, where the energy is either zero or infinity, with a radius equal to the distance threshold. The distance threshold was determined by selecting the 99.99-percentile distance sampled in the bound state, which resulted in 4.5830673 Åfor CB8-G3, 5.773037 Åfor OA-G3, and 6.0628217 Å for OA-G6. The YANK input file used for the calculation can be found at https://github.com/samplchallenges/SAMPL6/blob/master/host_guest/SAMPLing/YANK_input_script.yaml. The number of energy evaluations used to determine the computational cost of the method was computed for each iteration as MD_*cost*_ + MC_*cost*_ + MBAR_*cost*_, where MD_*cost*_ is the number of force evaluations used to propagate the system (i.e. 1 ps/2 fs = 500 force evaluations), MC_*cost*_ are the number of energy evaluations performed for acceptance/rejection of the MC rotation and translation (4 energy evaluations), and MBAR_*cost*_ is the number of energy evaluations necessary to compute the MBAR free energy matrix at each iteration. We set MBAR_*cost*_ = *K* × *K*, where *K* is both the number of states and the number of replicas. This is an overestimation as YANK computes the energies of each replica for all states by recomputing only the parts of the Hamiltonian that change from state to state.

### 6.3 Estimation of the relative efficiency

We considered the standard deviation, absolute bias, and RMSE error statistics in (Eq. 2, 4) to compute respectively the relative efficiencies *e*_std_, *e*_bias_, *e*_RMSE_. The relative efficiencies of all methods were estimated with respect to OpenMM/HREX, which was the longest calculation and could provide free energy predictions at all the computational costs intervals required to estimate the statistics. We used a uniform weight *w*(*c*) = const. for all methods, and, because we have data available for only 100 computational costs over the interval [*c*_*min,X*_, *c*_*max,X*_], we interpolated the error statistic for the other values of *c* and approximated the average over the number of energy evaluations with

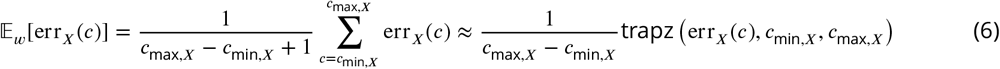

where trapz(·) represent the quadrature integral of the error function performed with the trapezoidal rule over the considered interval of *c*. The denominator does not affect the relative efficiency as it cancels out in Eq. (5).

The population mean 𝔼[Δ*G*(*c*)] and standard deviation std(Δ*G*(*c*)) of the binding free energy predictions at computational cost *c* were estimated as usual with the sample mean 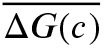 and the sample standard deviation *S*(*c*) respectively calculated using the five independent replicates

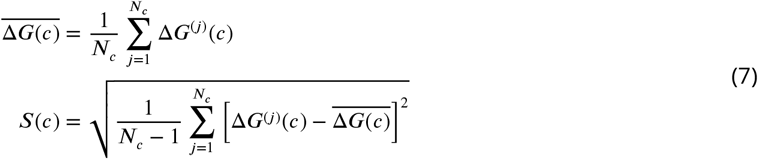

where *N_c_* = 5 is the number of independent measures at computational cost *c*.

However, estimating the error statistics defined in (Eq. 2, 4) requires estimates of the asymptotic free energy Δ*G_θ_*, which is necessary for the bias. This is problematic due to the different levels of convergence and the lack of agreement between methods. We estimated the bias assuming 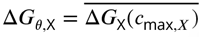, where is the total computational cost of the calculation for method *X*, which is equivalent to assuming that the free energy estimate has converged. As a consequence, the bias is generally underestimated, and longer calculations are penalized in computing the relative absolute bias and RMSE efficiency.

To estimate 95% confidence intervals for the relative efficiency measures we used the arch 4.6.0 Python library [134] to run the bias-corrected and accelerated (BCa) bootstrap method by resampling free energy trajectories with replacement. The acceleration parameter was estimated with the jackknife method.

## Code and data availability

- Input files and setup scripts: https://github.com/samplchallenges/SAMPL6/tree/master/host_guest/SAMPLing/
- Analysis scripts: https://github.com/samplchallenges/SAMPL6/tree/master/host_guest/Analysis/Scripts/
- Analysis results: https://github.com/samplchallenges/SAMPL6/tree/master/host_guest/Analysis/SAMPLing/
- Participant submissions: https://github.com/samplchallenges/SAMPL6/tree/master/host_guest/Analysis/Submissions/SAMPLing/

## Author Contributions

Conceptualization: DLM, AR, JDC, MS, JM; Data Curation: AR; Formal Analysis: AR; Funding Acquisition: JDC, DLM, MRS, MKG, AD, BLdG, JM, ZC; Investigation: AR, TJ, DRS, MA, VG, AD, DN, SB, NMH, MP; Methodology: AR, DLM, MKG, JM, JDC; Project Administration: AR, DLM, JDC; Resources: JDC, MRS, MKG, ZC, JM, AD, BLdG; Software: AR; Supervision: JDC, MRS, MKG, DLM, JM, ZC, AD, BLdG; Visualization: AR, VG, TJ; Writing – Original Draft: AR; Writing – Review & Editing: AR, MKG, JDC, DLM, DRS, MRS, MA, VG, JM, DN, AD, ZC, BLdG.

## Acknowledgments

AR and JDC acknowledge support from the Sloan Kettering Institute. JDC acknowledges support from NIH grant P30 CA008748. AR acknowledges partial support from the Tri-Institutional Program in Computational Biology and Medicine. DLM appreciates financial support from the National Institutes of Health (1R01GM108889-01 and 1R01GM124270-01A1) and the National Science Foundation (CHE 1352608). AR and JDC are grateful to OpenEye Scientific for providing a free academic software license for use in this work. MA was supported by a Postdoctoral Research Fellowship of the Alexander von Humboldt Foundation. VG and BLdG were supported by BioExcel CoE, a project funded by the European Union contract H2020-INFRAEDI-02-2018-823830. MKG thanks NIGMS (NIH) for partial support of this project (GM061300). The contents of this publication are solely the responsibility of the authors and do not necessarily represent the official views of the NIH. AD acknowledges support from the National Institutes of Health (R01GM130794) and the National Science Foundation (DMS 1761320). ZC and DN would like to thank Stamatia Zavitsanou, Michail Papadourakis and Chris Chipot for useful discussions. This work was further supported by computational time granted from the Greek Research & Technology Network (GRNET) in the National HPC facility - ARIS-under project ID vspr001005/arp2/3. ZC acknowledges support of this work by the project “An Open-Access Research Infrastructure of Chemical Biology and Target-Based Screening Technologies for Human and Animal Health, Agriculture and the Environment (OPENSCREEN-GR)” (MIS 5002691), which is implemented under the Action “Reinforcement of the Research and Innovation Infrastructure”, funded by the Operational Programme “Competitiveness, Entrepreneurship and Innovation” (NSRF 2014-2020) and co-financed by Greece and the European Union (European Regional Development Fund).

