## Supplementary Material for "The SAMPL6 SAMPLing challenge: Assessing the reliability and efficiency of binding free energy calculations"

### Appendix 1 - The relative efficiency is a robust statistic when data span different ranges of computational cost

Given two free energy methods, A and B, we want to use our efficiency statistic to accept or reject the hypothesis that A is more efficient than B, or *vice versa*. When the data available for methods A and B cover the same range of computational cost, then $\mathbb{E}_w[\mathrm{err}_A(c)]$ and $\mathbb{E}_w[\mathrm{err}_B(c)]$ can be directly compared, and standard statistical inference tools can be applied to the statistic defined in Eq. (3) in the main text. In this challenge, however, each submission provided data spanning very different ranges of computational cost. For example, GROMACS/EE does not provide free energy predictions during the initial part of the calculation corresponding to the equilibration stage, which is used to calibrate the expanded ensemble weights. Mean errors computed over different ranges of $c$ (i.e. different weight functions $w(c)$) cannot be meaningfully compared, and the analysis thus requires some attention. To understand why this is a problem, consider two runs of the same method, A' and A", for which the RMSE decays as a function of the computational cost according to a standard unbiased Monte Carlo model

$$\mathrm{RMSE}(c) = \frac{\alpha}{\sqrt{c}} \tag{1}$$

where $\alpha > 0$. The two calculations are identical, but the data available for A' and A" covers different intervals $[c_{\mathrm{min},A'}, c_{\mathrm{max},A'}]$ and $[c_{\mathrm{min},A''}, c_{\mathrm{max},A''}]$ respectively as sketched in SI Figure 1. A reasonable property to expect from our statistic is to assign the same inefficiency to data generated by the same method. However, using Eq. (1) in the main text and (1), we can compute the mean RMSE as

$$\mathbb{E}_w[\mathrm{RMSE}(c)] = \frac{\int_{c_{\mathrm{min}}}^{c_{\mathrm{max}}} \mathrm{RMSE}(c) dc}{c_{\mathrm{max}} - c_{\mathrm{min}}} = 2\alpha \frac{\sqrt{c_{\mathrm{max}}} - \sqrt{c_{\mathrm{min}}}}{c_{\mathrm{max}} - c_{\mathrm{min}}} \tag{2}$$

which implies that the mean RMSE of methods A and B will generally be different unless computed over the same cost interval. This is also evident from the submitted data, as can be seen in SI Figure 4.

In order to arrive at a statistic that can be properly compared, there are at least three solutions. The first, and simplest, is to compute the inefficiency statistic in the range $[c_{\mathrm{min}}, c_{\mathrm{max}}]$ for which there are data for all the methods and discard all data points outside the interval. Alternatively, when a robust model of the error decay is available, as with the example in Eq. (1), it may be possible to remove the dependency on the range of computational costs by appropriate scaling. In the example above, this could be achieved by comparing $\mathbb{E}_w[\mathrm{RMSE}(c)] \cdot \frac{c_{\mathrm{max}} - c_{\mathrm{min}}}{\sqrt{c_{\mathrm{max}}} - \sqrt{c_{\mathrm{min}}}}$ instead of $\mathbb{E}_w[\mathrm{RMSE}(c)]$. However, both these strategies are impractical here due to the very different ranges of $c$, which would require discarding between 50% and 75% of the data points for several methods, and the difficulty of finding a model for the error functions that could satisfactorily fit the data from all methodologies.

Instead, we decided to report and compare the inefficiency relative to a common reference method through the relative efficiency in Eq. 5 in the main text. We

**SI Figure 1: Relative efficiency is robust to differences in computational cost ranges.** Examples of RMSE trajectories for two hypothetical methods (Z and A) with decay proportional to $c^{-1/2}$. The mean error, $\mathbb{E}[RMSE]$, of two runs of an identical method (A' and A") is affected by the range of computational cost considered. On the other hand, the inset plot shows that the relative efficiencies with respect to the reference method Z computed from the two runs are identical. To be used as a reference, a method must span the full range of computational costs covered by the data.

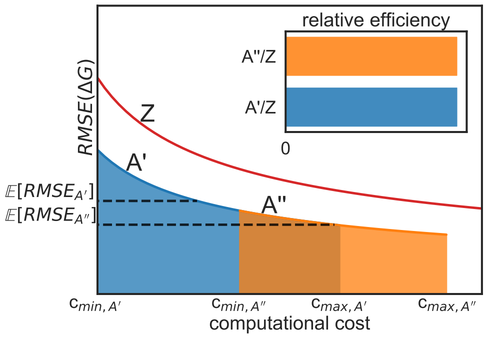


expect this statistic to be more robust with respect to the range of $c$ than the simple mean error. In particular, assuming the error function to obey the general model $\text{err}(c) = \alpha_X f(c)$ (e.g., $f(c) = c^{-1/2}$ in the example in Eq. (1)), with the constant $\alpha_X$ characterizing the decay rate of the method X, then the relative efficiency does not depend on the range of the computational costs

$$e_{\text{err},X/Z} = -\log_{10}\left(\frac{\alpha_X}{\alpha_Z}\right) \tag{3}$$

and the relative efficiency of different methods can be directly compared to each other even if their data spans different intervals (SI Figure 1). A meaningful comparison still requires the methods to obey the same decay model, but the key advantage here is that an explicit expression for $f(c)$ is not required. In practice, the relative efficiency and the ranking it produces seem to be relatively robust to differences in computational cost ranges for most methods (SI Figure 5) with fluctuations that are within the statistical uncertainty of the estimates (SI Figure 6)

### Appendix 2 - Sensitivity analysis of HREX calculations

In order to obtain insights into the origin of these differences, we focused on APR and HREX. The choice of focusing on these two methods was mainly due to technical feasibility as we considered it possible to run further HREX calculations after minimizing the differences in setups and other simulation parameters. This option was not available for investigating the differences between HREX and SOMD, for example, due to the lack of support for reaction field in YANK. Differences between HREX and other methods were not statistically significant or, in the case of CB8-G3 predictions from NAMD/BAR, likely the result of uncoverged calculations.

**SI Table 1: Summary of the free energy calculations run for the sensitivity analysis.** Average binding free energy predictions computed from five independent OpenMM/HREX calculations with 95% t-based confidence intervals under different simulation conditions. The AMBER/APR results are also reported in the first row to facilitate the comparison. All OpenMM/HREX binding free energies are reported after 20 ns/replica, including the one computed by the reference calculations after 40 ns/replica (second row). The HREX calculations were run by varying the Lennard-Jones cutoff, the exact value of the Coulomb constant, the restraint applied between host and guest, the Langevin discretization algorithm, and the box size and ion concentration. The only statistically significant difference in binding free energy was obtained after changing the cutoff from using a switching function between 9 Å and 10 Å to a 9 Å truncated cutoff.

| **method** | **LJ cutoff** | **Coulomb constant** | **Restraint** | **Langevin integrator discretization** | **Complex box size / ionic strength** | $\Delta$G [**kcal/mol**] |
|---|---|---|---|---|---|---|
| AMBER/APR | 9 Å truncated | AMBER | multiple | AMBER leap-frog | 44x43x67 Å$^3$ 73.9 mM | -6.3 ± 0.1 |
| OpenMM/HREX | 9–10 Å switched | OpenMM | harmonic | BAOAB | 43x43x43 Å$^3$ 64.3 mM | -6.7 ± 0.1 |
| OpenMM/HREX | 9 Å truncated | OpenMM | harmonic | BAOAB | 43x43x43 Å$^3$ 64.3 mM | -7.0 ± 0.1 |
| OpenMM/HREX | 9 Å truncated | AMBER | harmonic | BAOAB | 43x43x43 Å$^3$ 64.3 mM | -7.1 ± 0.1 |
| OpenMM/HREX | 9 Å truncated | AMBER | flat-bottom | BAOAB | 43x43x43 Å$^3$ 64.3 mM | -6.98 ± 0.08 |
| OpenMM/HREX | 9 Å truncated | AMBER | harmonic | OpenMM leap-frog | 43x43x43 Å$^3$ 64.3 mM | -7.14 ± 0.08 |
| OpenMM/HREX | 9 Å truncated | AMBER | harmonic | BAOAB | 44x43x67 Å$^3$ 73.9 mM | -7.1 ± 0.1 |

Moreover, we observed a systematic and statistically distinguishable difference of 0.3–0.4 kcal/mol in the final free energies from APR and HREX for all systems, which we found particularly curious. We verified by manual inspection that the distance between host and guest in the unbound state of APR was sufficient for the PMF to reach a plateau.

The conditions and the results of the additional HREX calculations are summarized in SI Table 1. All the new OpenMM/HREX calculations were run for 20 ns/replica (i.e. half the duration of the calculations in the original conditions), and we thus report in the table the original HREX binding free energy obtained at the same computational cost, which is statistically indistinguishable from the mean $\Delta G$ after 40 ns/replica. Surprisingly, simulating with a 9 Å truncated cutoff instead of using a switching function between 9 Å and 10 Å decreased the original OpenMM/HREX $\Delta G$ prediction by 0.3 kcal/mol, widening the difference between the two methods.

The source of this sensitivity may be connected to the central role of the van der Waals interactions in stabilizing the host-guest complex, and the size of the host, whose diameter is in the order of the cutoff. Changing the other parameters did not alter the binding free energy significantly. In particular, the predictions proved insensitive to the exact value of the Coulomb constant, which is slightly different in AMBER and OpenMM [6], and to the specific restraint used to restrict

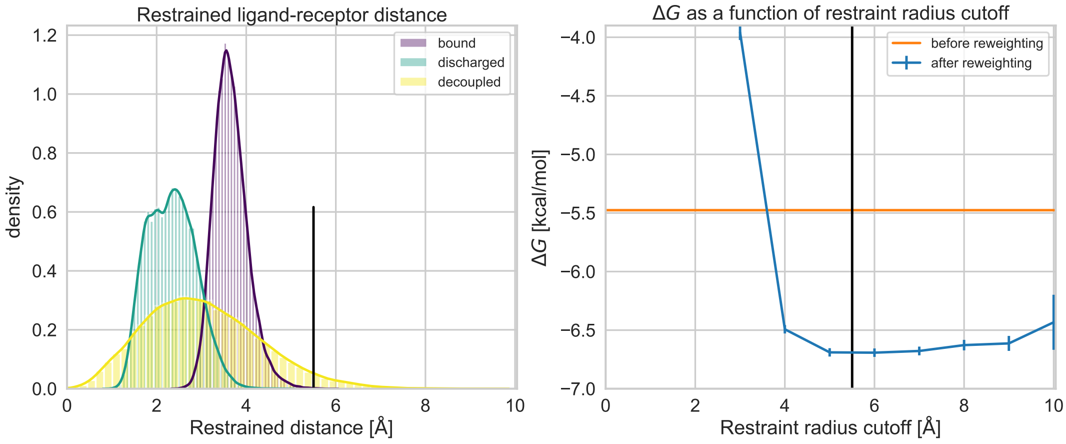


**SI Figure 2: OA-G3 restraint radius distributions, and binding free energy dependency on the binding site definition.** Distribution of the harmonic restraint radius (left) in the bound (purple), discharged (green), and decoupled state (yellow) for OA-G3-0, and predicted binding free energy as a function of the restraint radius cutoff (right). The black vertical line represents the threshold used during the reweighting analysis. The orange horizontal line in the right-bottom plot is the MBAR-predicted free energy of OA-G3-0 that did not undergo the reweighting procedure. The binding affinity is insensitive to the restraint cutoff radius between a large range of values that include most of the bound state distribution.

the conformational space available to the guest in the HREX calculation, which used a relatively tight harmonic potential (spring constant 0.17 kcal/mol/Å$^2$) in the original calculation and a more permissive flat-bottom potential (well radius 7.5 Å, spring constant 5 kcal/mol/Å$^2$) in the second case. We also investigated the impact of using a leapfrog Langevin integrator instead of a BAOAB discretization scheme, but this proved to be statistically insignificant as well with a timestep of 2 fs. It should be noted that the two leapfrog integration schemes provided in OpenMM [2] and AMBER [3] still have differences so it is still theoretically possible for the discretization error to contribute to the differences in free energy obtained by the two methods. We then re-ran OpenMM/HREX complex phase using the same input files generated for AMBER/APR. These solvation boxes were bigger to allow sampling long distances between host and guest, and the ionic strengths were slightly different. Again, the HREX binding free energy did not change significantly.

Finally, we examined the robustness of the reweighting step in YANK's analysis pipeline. Using a harmonic restraint in the bound state of the HREX calculation introduced a bias that was corrected by reweighting the data with MBAR to a state using a restraint following a square-well potential of a specific radius, which effectively defined the binding site (see also Detailed Methods). We thus looked into whether the binding free energy was sensitive to the radius of the binding site, and whether the reweighting procedure was statistically robust, or if an eventual poor overlap between the sampled and reweighted distribution could introduce significant statistical error. The results of the analysis represented in SI Figure 2 for OA-G3 show that very little statistical error is introduced in the reweighting process, and that the binding free energy is robust to the square well radius (i.e. the radius of the defined binding site), as expected from a tight binder [1]. Moreover, comparing the distributions of the restraint radius sampled in the bound and decoupled states, with the latter distribution having much larger support than the former (SI Figure 2), suggests that the spring constant of the harmonic potential was appropriate and did not limit the exploration of the binding site in the bound state.

### Appendix 3 - Incorrect initialization of intermediate states with long correlation time can bias MBAR estimates

We show here, with the help of a toy example, how initializing all the replicas with a bound-state conformation can introduce bias of negative sign in the MBAR binding free energy estimates when intermediate states decorrelate very slowly.

This phenomenon is evident by inspecting the expression of the MBAR estimator [4, 5], which first constructs the mixture distribution from all the available samples

$$p_M(\mathbf{x}) = \sum_{k=1}^{K} \frac{N_k}{N} \frac{e^{-u_k(\mathbf{x})}}{\hat{Z}_k} \tag{4}$$

where $K$ is the number of states, $N_k$ is the number of samples from state $k$, $N$ is the total number of samples from all states, and $\hat{Z}_k$ is the partition function of state $k$, and then it computes the free energy as a weighted average of the Boltzmann weight of state $i$ over all samples as

$$\hat{f}_i = -\ln \left( \frac{1}{N} \sum_{j=1}^{K} \sum_{n=1}^{N_j} \frac{e^{-u_i(\mathbf{x}_{jn})}}{p_M(\mathbf{x}_{jn})} \right) \tag{5}$$

If the sampling of any intermediate state starts from a bound-state conformation, and the state is affected by long correlation times, the samples collected at the beginning of the simulation will be biased towards the bound state, and the average Boltzmann weight of the bound state could be overestimated. This would result in binding free energies that are favorable towards the bound state, or in other words, it will introduce a bias of negative sign and result in more negative binding free energies. Trivially, this can be understood directly from Eq. (5) by recognizing that the denominators of each addend (i.e. $p_M(\mathbf{x}_{jn})$) does not depend on $i$ while, if the sampling of intermediate states is biased towards the bound state, the numerators for the bound state will be on average greater than they should be if the intermediate states were sampled from the correct equilibrium distributions.

We can verify this numerically with a very simple model. The Boltzmann distribution of a harmonic oscillator with spring constant $K$ and equilibrium length $\mu$ is equivalent to the Gaussian distribution

$$p(x|\mu, \sigma) = \frac{1}{\sqrt{2\pi}\sigma} \int e^{-\frac{(x-\mu_1)^2}{2\sigma^2}} dx \tag{6}$$

where $\sigma = \sqrt{K^{-1}}$. Consider now a transformation of a harmonic oscillator from $(\mu_0, \sigma_0)$ to $(\mu_1, \sigma_1)$ to $(\mu_2, \sigma_2)$. With the values of $\mu$ and $\sigma$ in Table 2, the distributions have, in some sense, similar entropic characteristics of the $\lambda$ windows of a typical absolute alchemical free energy calculation (note the resemblance to the restraint distance distribution in SI Figure 2) so, with the sole intention of making the example more immediate, we refer to $(\mu_0, \sigma_0)$, $(\mu_1, \sigma_1)$, and $(\mu_2, \sigma_2)$ as the bound, intermediate, and decoupled state respectively. The dimensionless free energy difference between states $i$ and $j$ can be computed analytically as $\Delta F_{ij} = F_j - F_i = -\ln(\sigma_j/\sigma_i)$, and, with perfect sampling, the overlap between the distributions is sufficient for both BAR and MBAR to estimate the difference in free energy between the states correctly (Table 2). We can simulate the scenario in which the sampling of the intermediate state are biased towards the bound state but the decoupled state decorrelate fast simply by collecting the intermediate state samples from the bound state distribution with $\mu_1 = \mu_0 = 0.0$ and $\sigma_1 = \sigma_0 = 2.0$. In this case, even if both the bound and decoupled states are sampled correctly, all the MBAR predictions are biased towards free energies that are favorable to the bound state (Table 2).

**SI Table 2:** MBAR predictions for harmonic oscillators of different equilibrium lengths and standard deviation with perfect and biased sampling. Five million samples were collected for each Gaaussian distribution. Uncertainties of the predicted free energies are given as two times the MBAR uncertainty estimate. With biased sampling, state 1 was sampled from distribution 0 to simulate a slowly decorrelating intermediate state from an initial conformation typical of state 0. When the sampling of an intermediate state is biased towards one of the end states, the free energy prediction will be favorable towards that state.

| state | $\mu$ | $\sigma$ | true $\Delta F_{i1}$ | BAR $\Delta F_{i1}$ perfect sampling | MBAR $\Delta F_{i1}$ perfect sampling | MBAR $\Delta F_{i1}$ biased sampling |
|---|---|---|---|---|---|---|
| 0 | 0.0 | 2.0 | 0.0 | 0.0 | 0.00 | 0.0 |
| 1 | 2.0 | 2.0 | 0.0 | 0.0005 ± 0.0006 | -0.0001 ± 0.0004 | -0.2954 ± 0.0004 |
| 2 | 2.0 | 5.0 | 0.9163 | 0.9151 ± 0.0008 | 0.9155 ± 0.0006 | 0.7486 ± 0.0006 |

**SI Figure 3:** Schematics of the strategy adopted to simulated perfect sampling (left) and sampling of the "intermediate" state (state 1, green) biased towards the "bound" state (state 0, blue). In both cases, the "decoupled" state (state 2, orange) is assumed to decorrelate quickly.

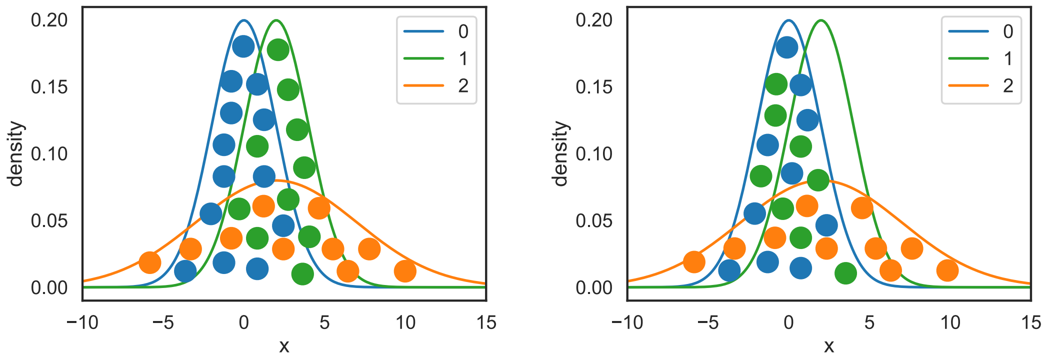

### Supporting Figures

**SI Figure 4: Mean error of all submissions as a function of the calculation length.** Mean standard deviation, absolute bias, and RMSE computed were computed with Eq. (8) in the main text, considering an increasing number of energy evaluations. The dash line is used for the method used as a reference in the calculation of the relative efficiencies. Because of the strong dependence on the number of energy evaluations, a meaningful comparison between methods using this statistic should be restricted to the range of computational cost for which data is available for all method.

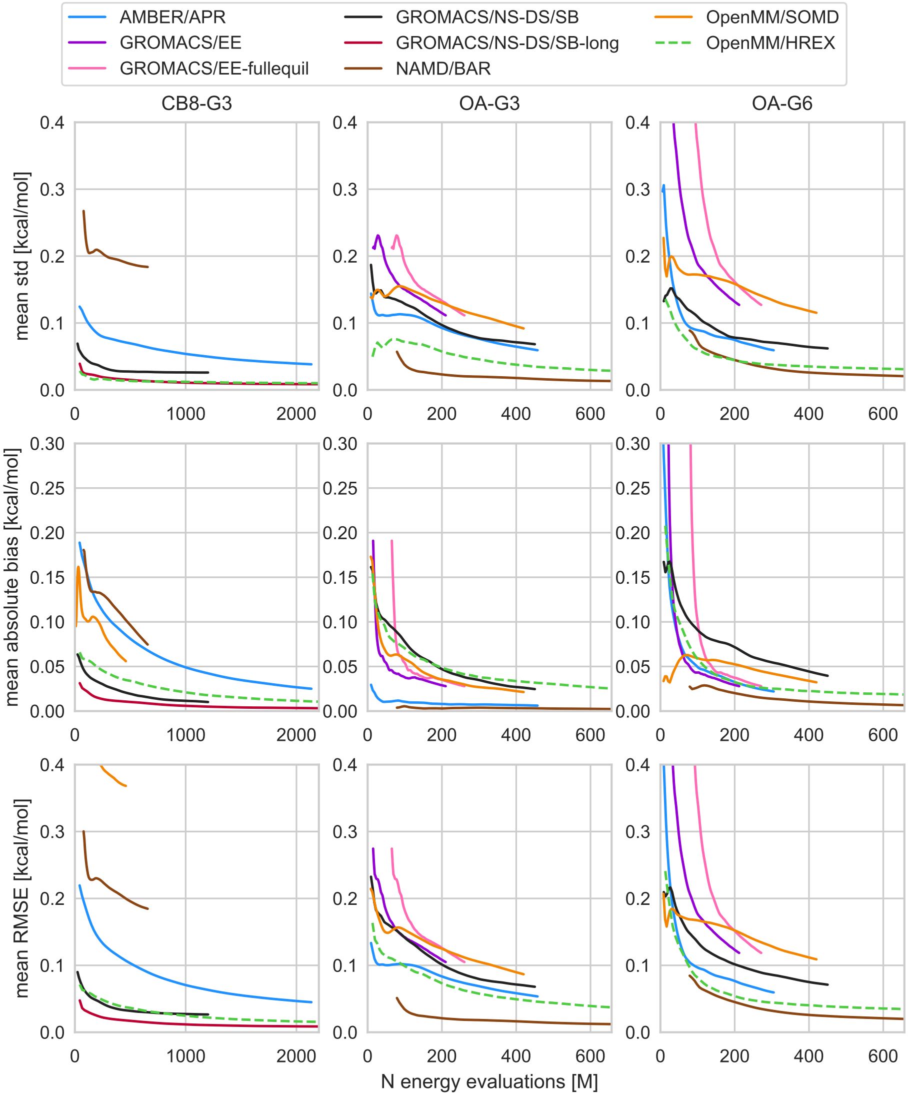

**SI Figure 5: Relative efficiencies with respect to OpenMM/HREX as a function of the calculation length.** Ratio of mean standard deviation, absolute bias, and RMSE computed considering an increasing number of energy evaluations. After an initial transient, the ranking of different methods based on the relative efficiency is fairly independent on the calculation lengths.

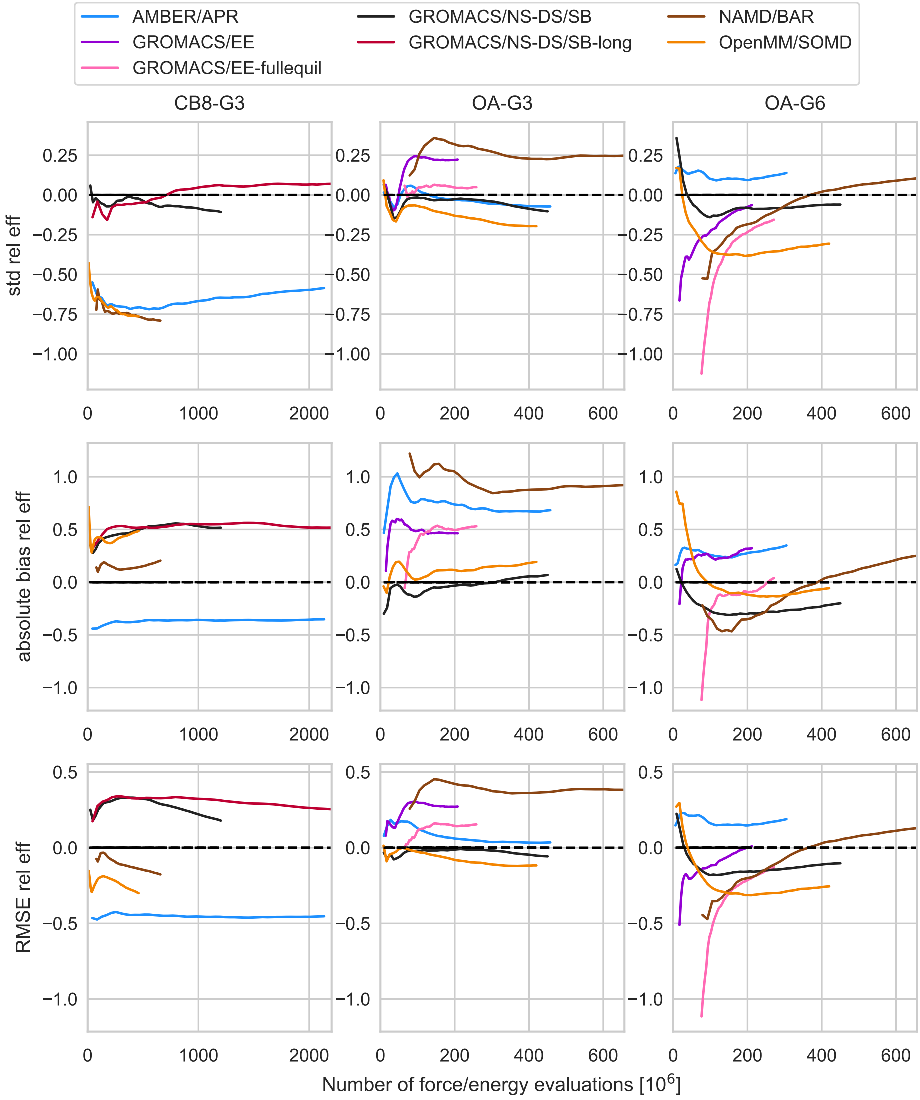

**SI Figure 6: Relative efficiency as a function of the computational cost with bootstrap 95% confidence interval for all methods.** Ratio of mean standard deviation, absolute bias, and RMSE computed considering an increasing number of energy evaluations. The confidence interval were computed with bias-corrected and accelerated bootstrap. A confidence interval entirely above/below 1 is evidence of a better performance of APR/HREX.

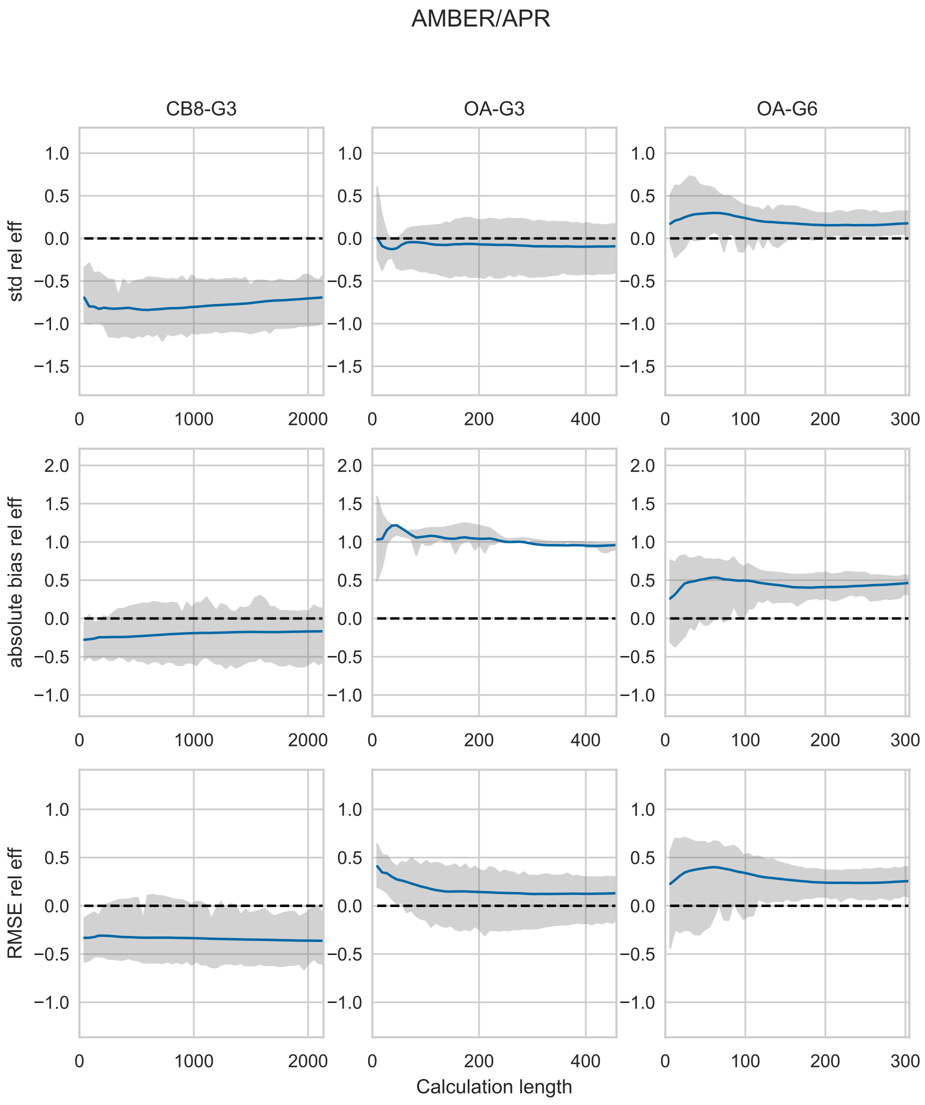

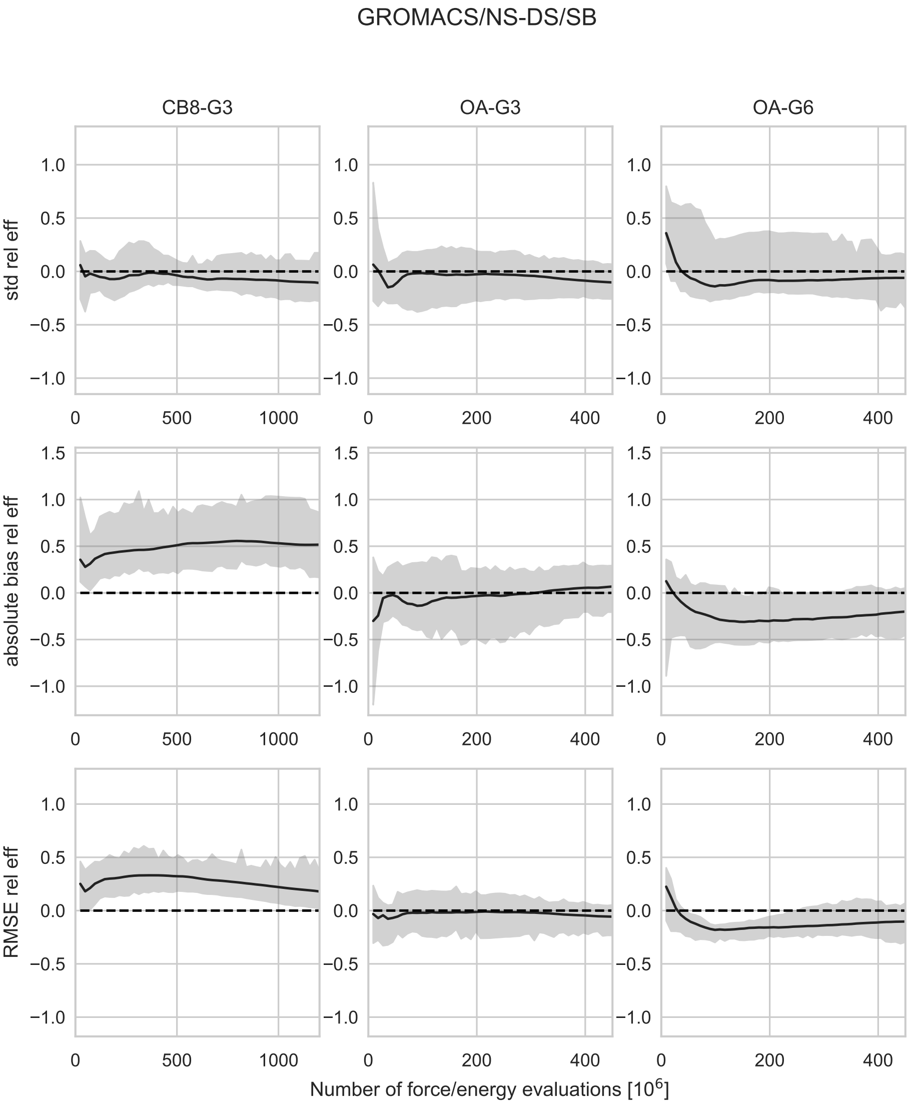

GROMACS/NS-DS/SB
CB8-G3
OA-G3
OA-G6
std rel eff
absolute bias rel eff
RMSE rel eff
Number of force/energy evaluations [10^6]

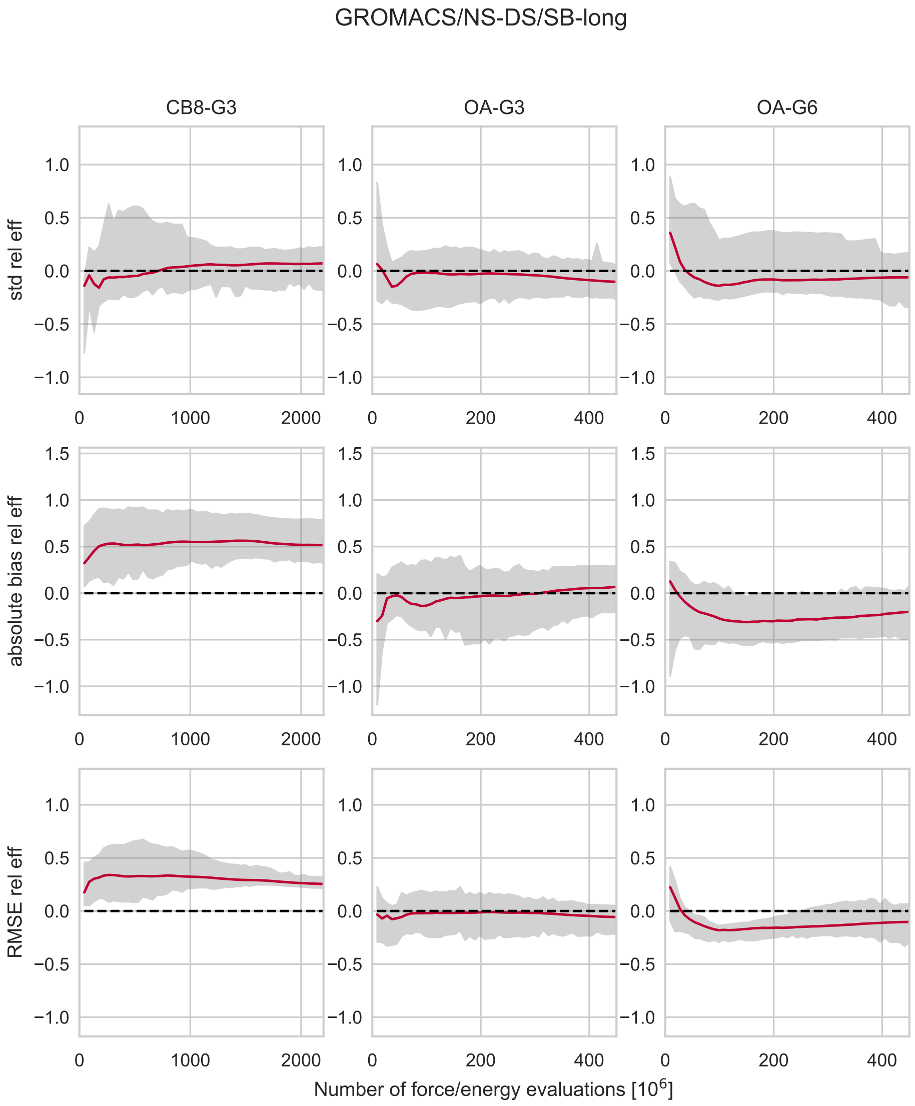

GROMACS/NS-DS/SB-long
CB8-G3
OA-G3
OA-G6
std rel eff
absolute bias rel eff
RMSE rel eff
Number of force/energy evaluations [10^6]

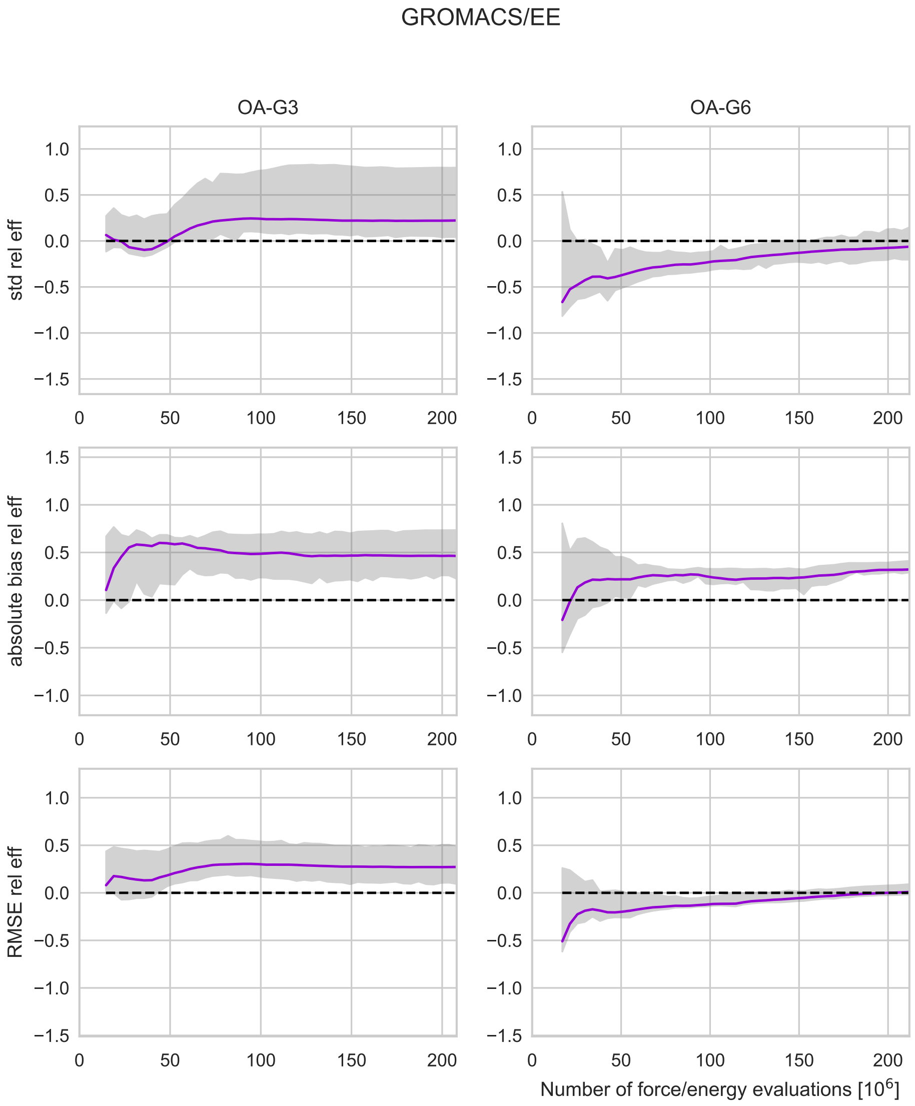
GROMACS/EE
OA-G3
OA-G6
std rel eff
absolute bias rel eff
RMSE rel eff
Number of force/energy evaluations [$10^6$]

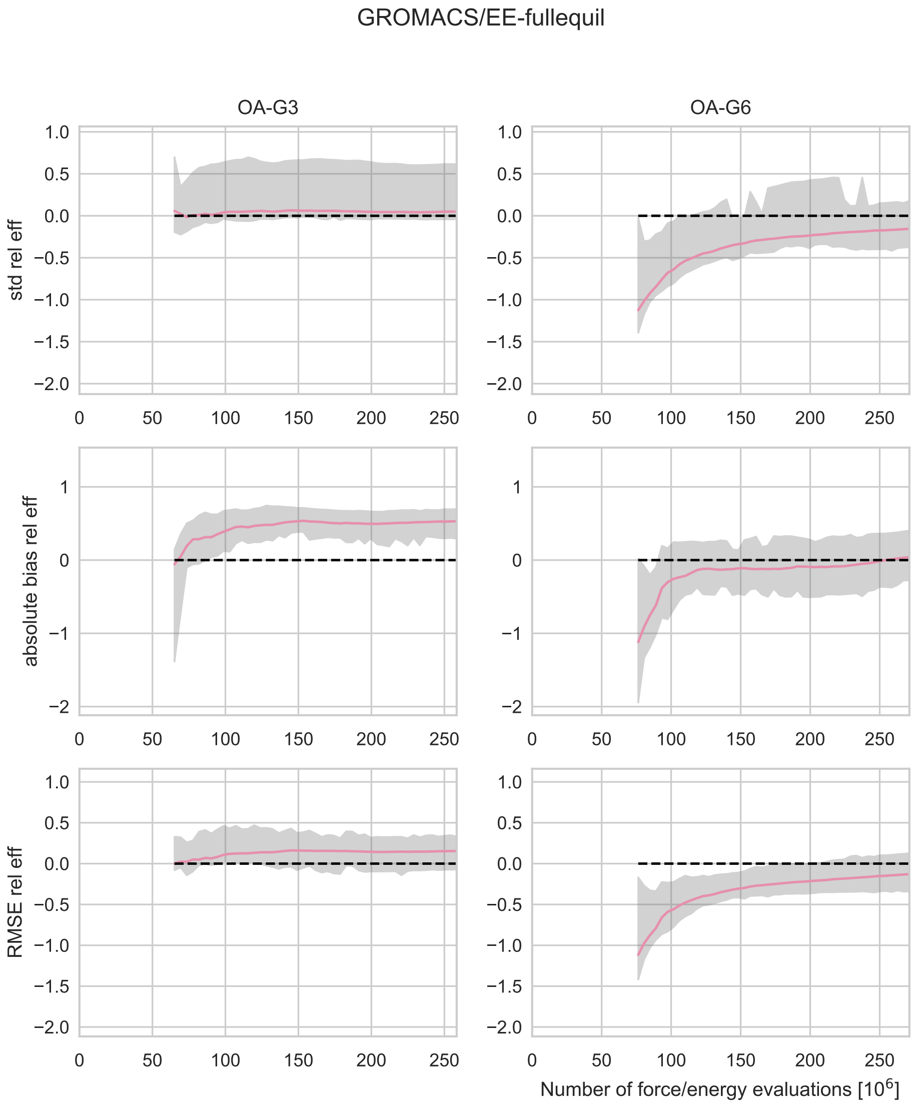

GROMACS/EE-fullequil
OA-G3
OA-G6
std rel eff
absolute bias rel eff
RMSE rel eff
Number of force/energy evaluations [10^6]

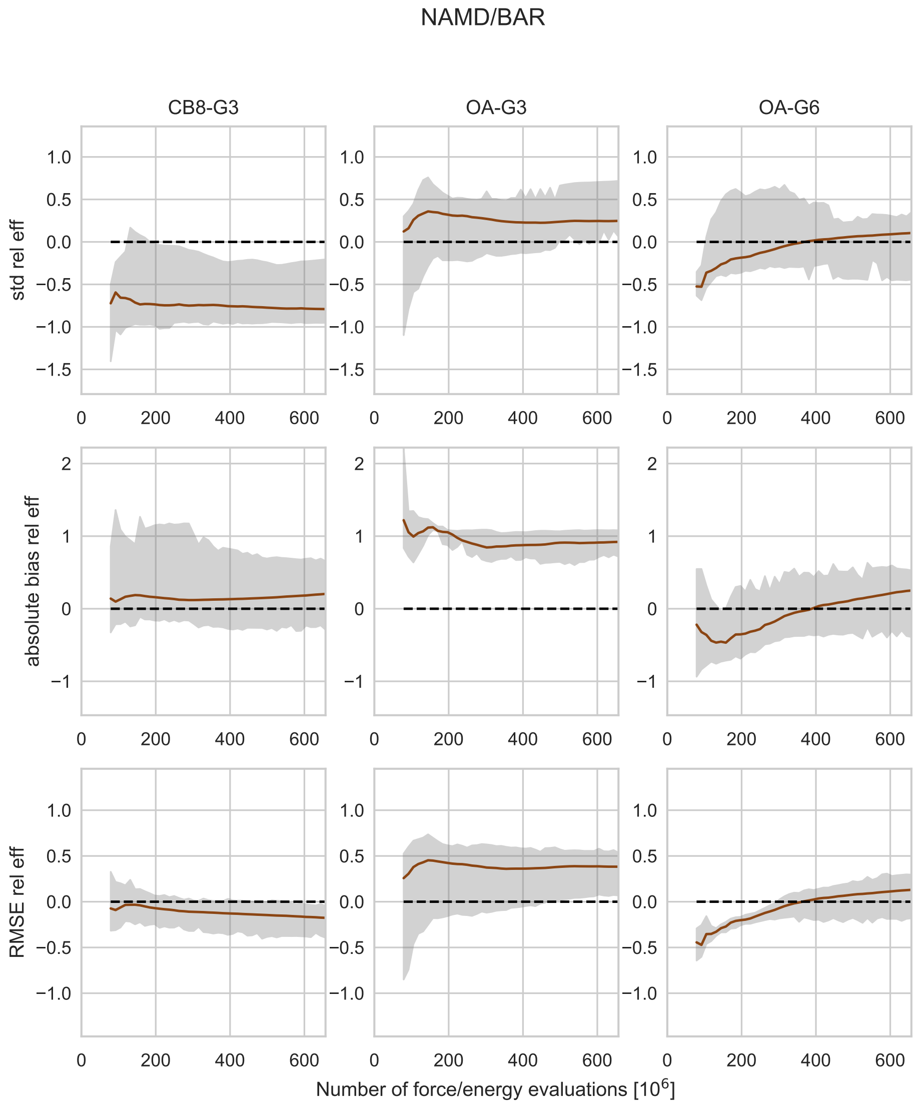
NAMD/BAR
CB8-G3
OA-G3
OA-G6
std rel eff
absolute bias rel eff
RMSE rel eff
Number of force/energy evaluations [10^6]

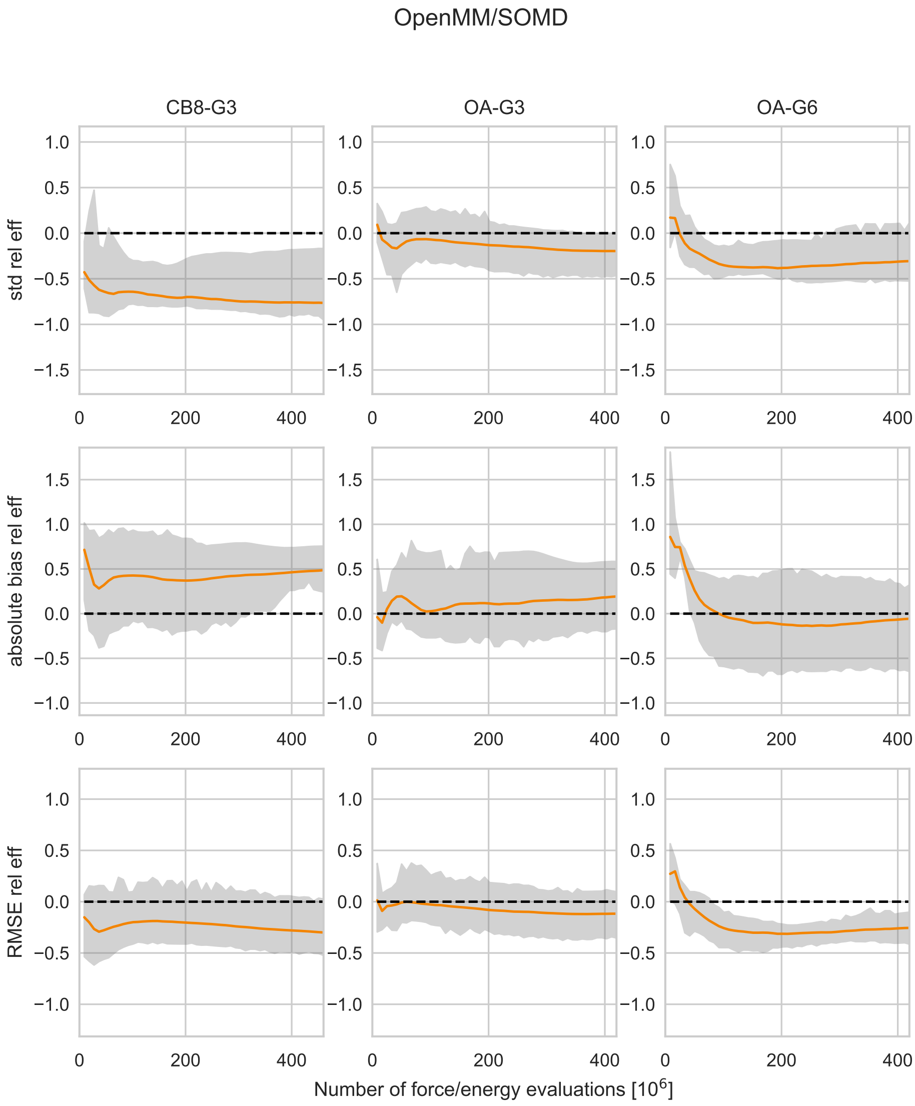

OpenMM/SOMD
CB8-G3
OA-G3
OA-G6
std rel eff
absolute bias rel eff
RMSE rel eff
Number of force/energy evaluations [10^6]

**SI Figure 7: Free energy, standard deviation, and bias as a function of computational cost.** The trajectories and shaded areas in the top row represent the mean binding free energies and 95% t-based confidence intervals computed from the 5 replicate predictions for CB8-G3 (left column), OA-G3 (center), and OA-G6 (right) for all submissions. The second and third row show as a function of the computational effort the standard deviation and the bias respectively.

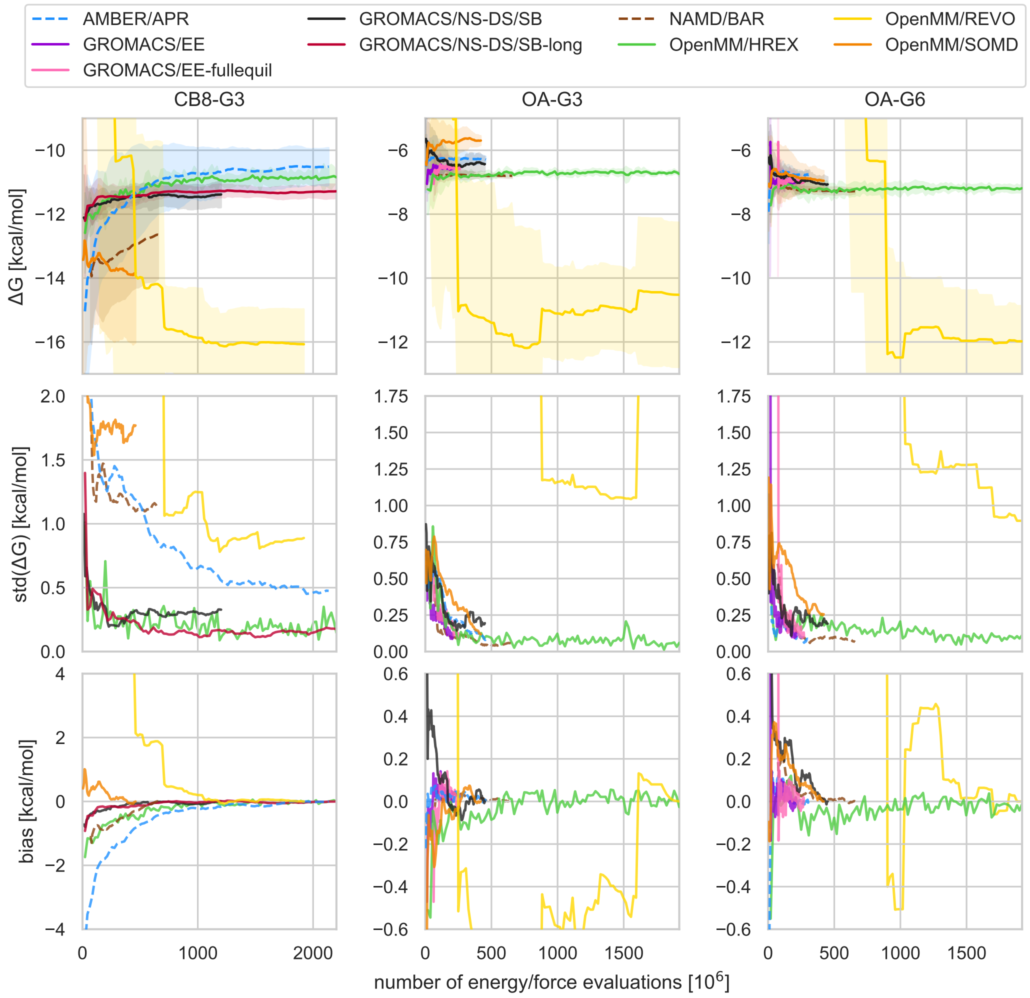

**SI Table 3: Final binding free energy predictions of the single replicate calculations.** Estimates of binding free energy predictions with uncertainty submitted by all participants for the individual replicate calculations. The results for GROMACS/EE-fullequil are identical to GROMACS/EE.

| | | | **CB8-G3** | | | | | **OA-G3** | | | | | **OA-G6** | | |
|---|---|---|---|---|---|---|---|---|---|---|---|---|---|---|---|
| **Method** | CB8-G3-0 | CB8-G3-1 | CB8-G3-2 | CB8-G3-3 | CB8-G3-4 | OA-G3-0 | OA-G3-1 | OA-G3-2 | OA-G3-3 | OA-G3-4 | OA-G6-0 | OA-G6-1 | OA-G6-2 | OA-G6-3 | OA-G6-4 |
| AMBER/APR | -10.9 ± 0.4 | -9.9 ± 0.4 | -10.1 ± 0.3 | -11.0 ± 0.4 | -10.7 ± 0.3 | -6.4 ± 0.1 | -6.2 ± 0.1 | -6.3 ± 0.1 | -6.2 ± 0.1 | -6.3 ± 0.1 | -6.8 ± 0.2 | -6.9 ± 0.2 | -6.8 ± 0.2 | -6.8 ± 0.2 | -6.6 ± 0.2 |
| GROMACS/NS-DS/SB | -11.5 ± 0.3 | -11.1 ± 0.2 | -11.8 ± 0.2 | -11.0 ± 0.2 | -11.4 ± 0.1 | -6.2 ± 0.2 | -6.4 ± 0.2 | -6.4 ± 0.1 | -6.6 ± 0.1 | -6.6 ± 0.2 | -7.3 ± 0.3 | -7.2 ± 0.2 | -7.0 ± 0.2 | -7.0 ± 0.1 | -6.8 ± 0.1 |
| GROMACS/NS-DS/SB-long | -11.4 ± 0.2 | -11.2 ± 0.2 | -11.4 ± 0.1 | -11.4 ± 0.1 | -11.0 ± 0.1 | | | | | | | | | | |
| GROMACS/EE | | | | | | -6.49 ± 0.06 | -6.76 ± 0.05 | -6.57 ± 0.05 | -6.56 ± 0.05 | -6.54 ± 0.05 | -6.96 ± 0.05 | -6.98 ± 0.05 | -6.97 ± 0.05 | -7.16 ± 0.05 | -6.87 ± 0.05 |
| GROMACS/NS-Gauss-F | 5.0 ± 2.0 | -2.0 ± 1.0 | -5.0 ± 1.0 | -3.2 ± 0.8 | 13.0 ± 1.0 | -6.7 ± 0.3 | -5.1 ± 0.6 | -6.3 ± 0.2 | -5.1 ± 0.6 | -6.2 ± 0.4 | -6.2 ± 0.4 | -5.4 ± 0.4 | -5.8 ± 0.4 | -5.1 ± 0.6 | -2.5 ± 0.6 |
| GROMACS/NS-Gauss-R | -14.3 ± 0.5 | -13.3 ± 0.6 | -12.2 ± 0.2 | -14.7 ± 0.8 | -14.5 ± 0.4 | -17.0 ± 2.0 | -30.0 ± 20.0 | -50.0 ± 20.0 | -11.0 ± 1.0 | -11.6 ± 1.0 | -22.0 ± 2.0 | -14.4 ± 1.0 | -21.0 ± 3.0 | -17.0 ± 2.0 | -16.0 ± 1.0 |
| GROMACS/NS-Jarz-F | -13.0 ± 0.3 | -12.3 ± 0.3 | -12.4 ± 0.3 | -13.1 ± 0.3 | -12.4 ± 0.2 | -8.1 ± 0.2 | -7.9 ± 0.3 | -7.7 ± 0.3 | -6.8 ± 0.3 | -6.7 ± 0.3 | -5.4 ± 0.6 | -7.7 ± 0.3 | -8.5 ± 0.3 | -7.7 ± 0.3 | -7.3 ± 0.4 |
| GROMACS/NS-Jarz-R | -10.5 ± 0.4 | -10.3 ± 0.3 | -10.8 ± 0.3 | -10.1 ± 0.3 | -10.2 ± 0.3 | -5.0 ± 0.4 | -6.0 ± 0.4 | -5.3 ± 0.2 | -5.8 ± 0.3 | -5.9 ± 0.4 | -6.7 ± 0.5 | -7.1 ± 0.4 | -5.3 ± 0.3 | -6.2 ± 0.3 | -5.6 ± 0.2 |
| NAMD/BAR | -11.7 ± 0.3 | -13.9 ± 0.3 | -11.4 ± 0.3 | -13.7 ± 0.2 | -12.4 ± 0.2 | -6.85 ± 0.05 | -6.82 ± 0.05 | -6.78 ± 0.05 | -6.71 ± 0.05 | -6.85 ± 0.05 | -7.17 ± 0.06 | -7.31 ± 0.05 | -7.33 ± 0.05 | -7.28 ± 0.06 | -7.32 ± 0.05 |
| OpenMM/REVO | -17.3 ± 0.8 | -15.8 ± 0.8 | -16.0 ± 0.8 | -14.9 ± 0.8 | -16.4 ± 0.8 | -11.0 ± 2.0 | -11.0 ± 2.0 | -13.0 ± 2.0 | -10.0 ± 2.0 | -8.0 ± 2.0 | -11.6 ± 0.8 | -12.1 ± 0.8 | -12.0 ± 0.8 | -10.9 ± 0.8 | -13.3 ± 0.8 |
| OpenMM/SOMD | -15.0 ± 0.1 | -14.03 ± 0.09 | -11.0 ± 0.1 | -13.71 ± 0.09 | -15.54 ± 0.09 | -5.75 ± 0.03 | -5.85 ± 0.03 | -5.55 ± 0.03 | -5.72 ± 0.03 | -5.61 ± 0.03 | -6.73 ± 0.03 | -7.04 ± 0.03 | -6.91 ± 0.03 | -6.74 ± 0.03 | -7.35 ± 0.03 |
| OpenMM/HREX | -10.82 ± 0.07 | -10.97 ± 0.07 | -10.88 ± 0.07 | -10.56 ± 0.07 | -11.0 ± 0.07 | -6.75 ± 0.04 | -6.65 ± 0.04 | -6.74 ± 0.04 | -6.71 ± 0.04 | -6.73 ± 0.06 | -7.19 ± 0.05 | -7.18 ± 0.05 | -7.12 ± 0.05 | -7.15 ± 0.05 | -7.24 ± 0.05 |

**SI Table 4: Relative efficiency of unidirectional nonequilibrium estimators in comparison to BAR.** Relative efficiencies of a method $X$ are reported with respect to GROMACS/NS-DS/SB-long as $e_{\mathrm{err},X/\mathrm{GROMACS/NS\text{-}DS/SB\text{-}long}}$ as defined by Eq. (6) in the main text. The lower and upper bound of the 95% confidence intervals bootstrap estimates for the relative efficiencies are reported as subscript and superscript respectively. The theoretical free energy used to compute the bias was estimated as the final average free energy of the 5 replicates predictions of GROMACS/NS-DS/SB-long.

| Method | CB8-G3 | | | OA-G3 | | | OA-G6 | | |
|---|---|---|---|---|---|---|---|---|---|
| | $e_{\mathrm{std}}$ | $e_{\lvert\mathrm{bias}\rvert}$ | $e_{\mathrm{RMSE}}$ | $e_{\mathrm{std}}$ | $e_{\lvert\mathrm{bias}\rvert}$ | $e_{\mathrm{RMSE}}$ | $e_{\mathrm{std}}$ | $e_{\lvert\mathrm{bias}\rvert}$ | $e_{\mathrm{RMSE}}$ |
| GROMACS/NS-Gauss-F | $-1.5^{-0.7}_{-1.8}$ | $-2.0^{-1.5}_{-2.3}$ | $-1.7^{-1.4}_{-2.0}$ | $-0.5^{-0.3}_{-0.7}$ | $-0.5^{0.2}_{-0.8}$ | $-0.5^{-0.2}_{-0.6}$ | $-0.7^{-0.4}_{-0.8}$ | $-0.9^{-0.8}_{-1.0}$ | $-0.8^{-0.6}_{-0.8}$ |
| GROMACS/NS-Gauss-R | $-0.7^{-0.4}_{-1.0}$ | $-1.4^{-1.1}_{-1.7}$ | $-1.1^{-0.9}_{-1.3}$ | $-1.7^{-0.8}_{-2.0}$ | $-2.0^{-1.5}_{-2.5}$ | $-1.8^{-1.1}_{-2.1}$ | $-1.1^{-0.9}_{-1.4}$ | $-1.6^{-1.2}_{-2.0}$ | $-1.4^{-1.2}_{-1.6}$ |
| GROMACS/NS-Jarz-F | $-0.2^{0.2}_{-0.4}$ | $-1.0^{-0.7}_{-1.4}$ | $-0.8^{-0.5}_{-1.0}$ | $-0.4^{-0.2}_{-0.9}$ | $-0.8^{-0.2}_{-1.2}$ | $-0.5^{0.1}_{-0.8}$ | $-0.6^{-0.4}_{-0.8}$ | $-0.1^{0.4}_{-0.6}$ | $-0.5^{-0.0}_{-0.8}$ |
| GROMACS/NS-Jarz-R | $-0.3^{-0.1}_{-0.7}$ | $-0.8^{-0.4}_{-1.2}$ | $-0.6^{-0.3}_{-0.8}$ | $-0.3^{0.0}_{-0.5}$ | $-0.9^{-0.7}_{-1.2}$ | $-0.6^{-0.3}_{-0.8}$ | $-0.3^{-0.1}_{-0.8}$ | $-0.7^{-0.4}_{-1.0}$ | $-0.5^{-0.4}_{-0.7}$ |
| **GROMACS/NS-DS/SB-long** | 0.0 | 0.0 | 0.0 | 0.0 | 0.0 | 0.0 | 0.0 | 0.0 | 0.0 |

**SI Figure 8: Volume fluctuations sampled by the Berendsen barostat in expanded ensemble calculations by state.** Average volume sampled in each intermediate state in the complex stage of the double decoupling calculation. State 0 is the bound state and state 39 is the decoupled state. Error bars are standard deviations of the volume distribution.

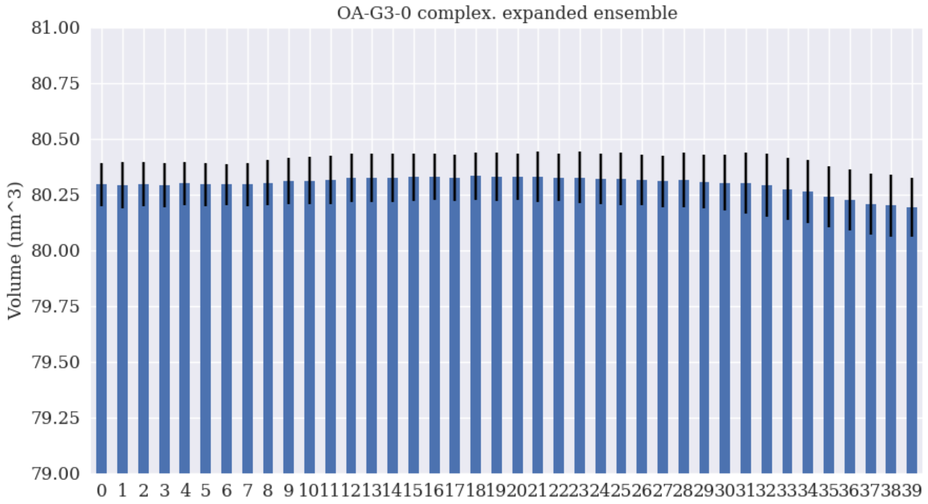

**SI Figure 9: Comparison of single-replicate uncertainty estimates and 5-replicate standard deviation.** Binding free energy trajectories (top row) and their uncertainty estimates (bottom row) as a function of the number of force/energy evaluations. The estimates for the individual replicate calculations were submitted by the challenge participants, and they are plotted in pastel colors. The best available estimates (in black) for $\Delta G$ and std($\Delta G$) were taken to be respectively the mean and standard deviation of 5 individual $\Delta G$ trajectories. A 95% chi-based confidence interval is plotted as a gray shaded area around the best estimate of the standard deviation.

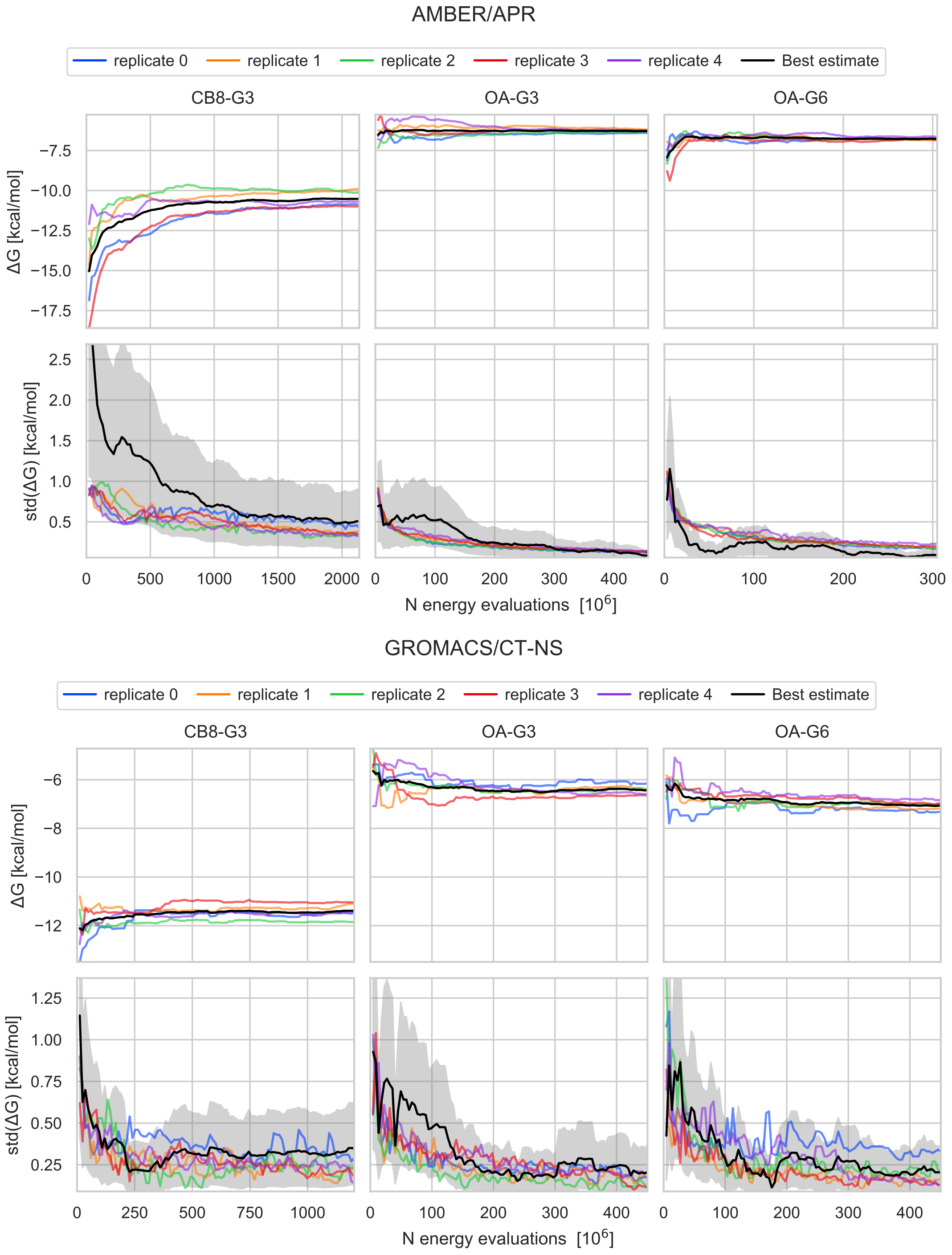

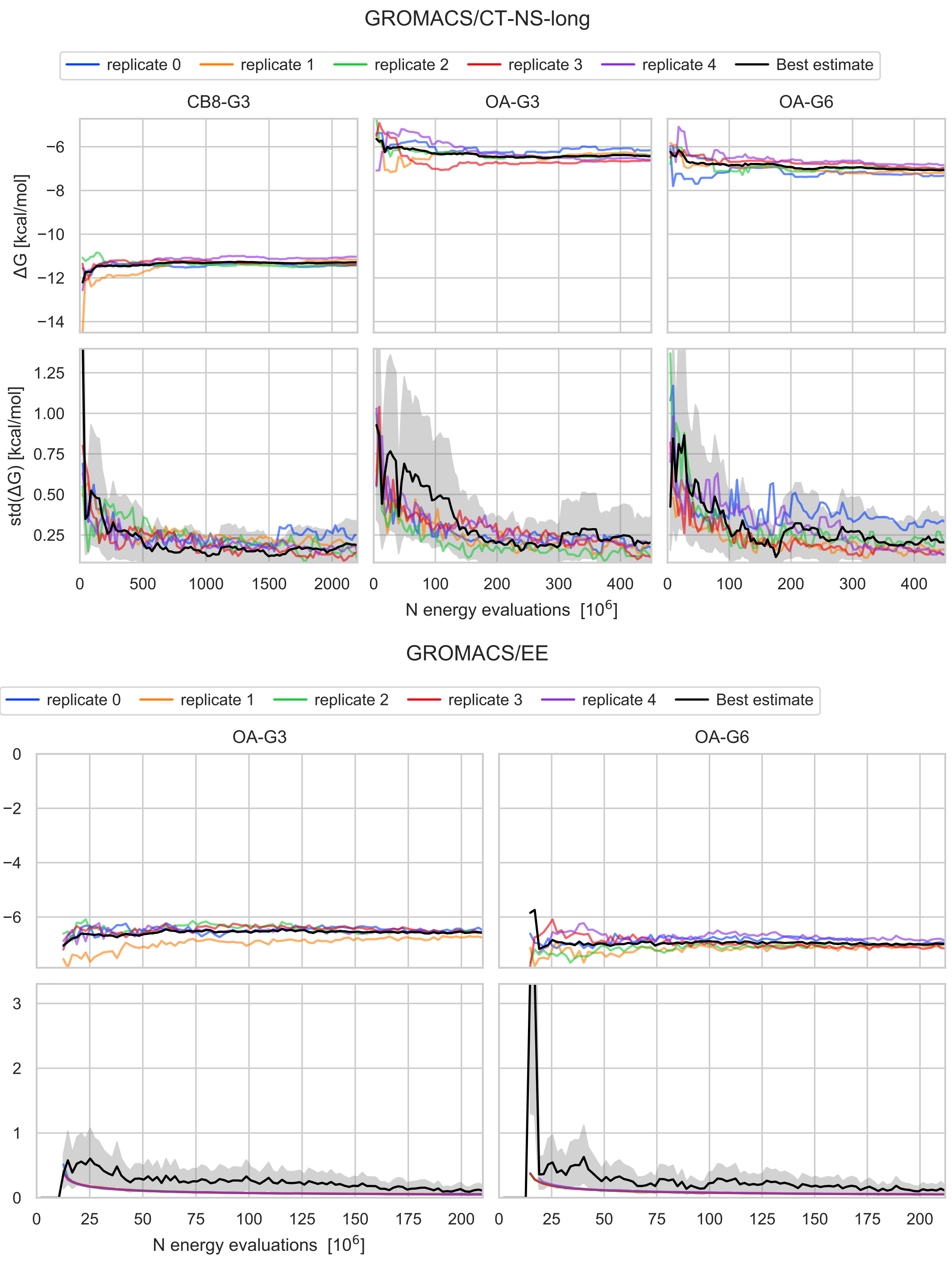
GROMACS/CT-NS-long
replicate 0
replicate 1
replicate 2
replicate 3
replicate 4
Best estimate
CB8-G3
OA-G3
OA-G6
ΔG [kcal/mol]
std(ΔG) [kcal/mol]
N energy evaluations [10^6]
GROMACS/EE
replicate 0
replicate 1
replicate 2
replicate 3
replicate 4
Best estimate
OA-G3
OA-G6
N energy evaluations [10^6]

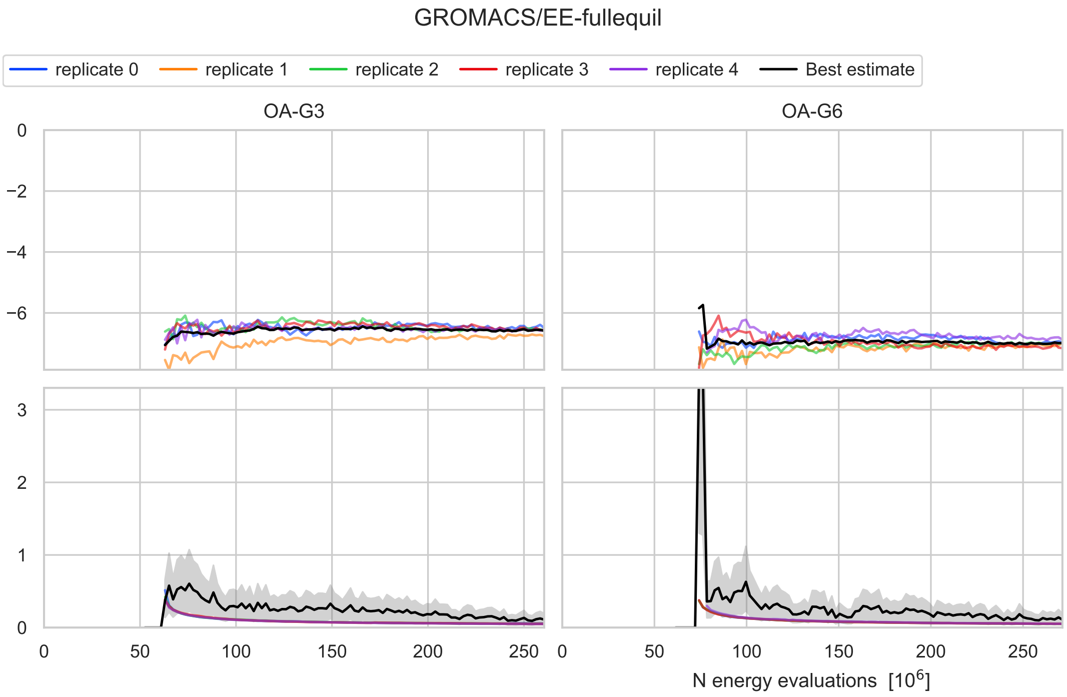
GROMACS/EE-fullequil
replicate 0
replicate 1
replicate 2
replicate 3
replicate 4
Best estimate
OA-G3
OA-G6
N energy evaluations [10^6]

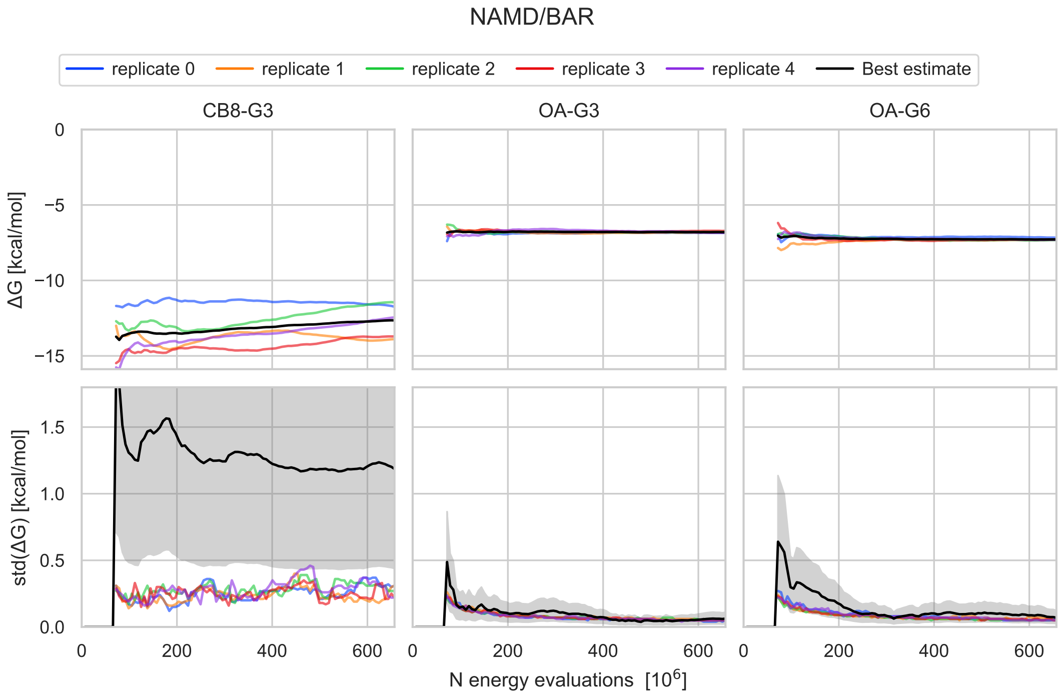
NAMD/BAR
replicate 0
replicate 1
replicate 2
replicate 3
replicate 4
Best estimate
CB8-G3
OA-G3
OA-G6
ΔG [kcal/mol]
std(ΔG) [kcal/mol]
N energy evaluations [10^6]

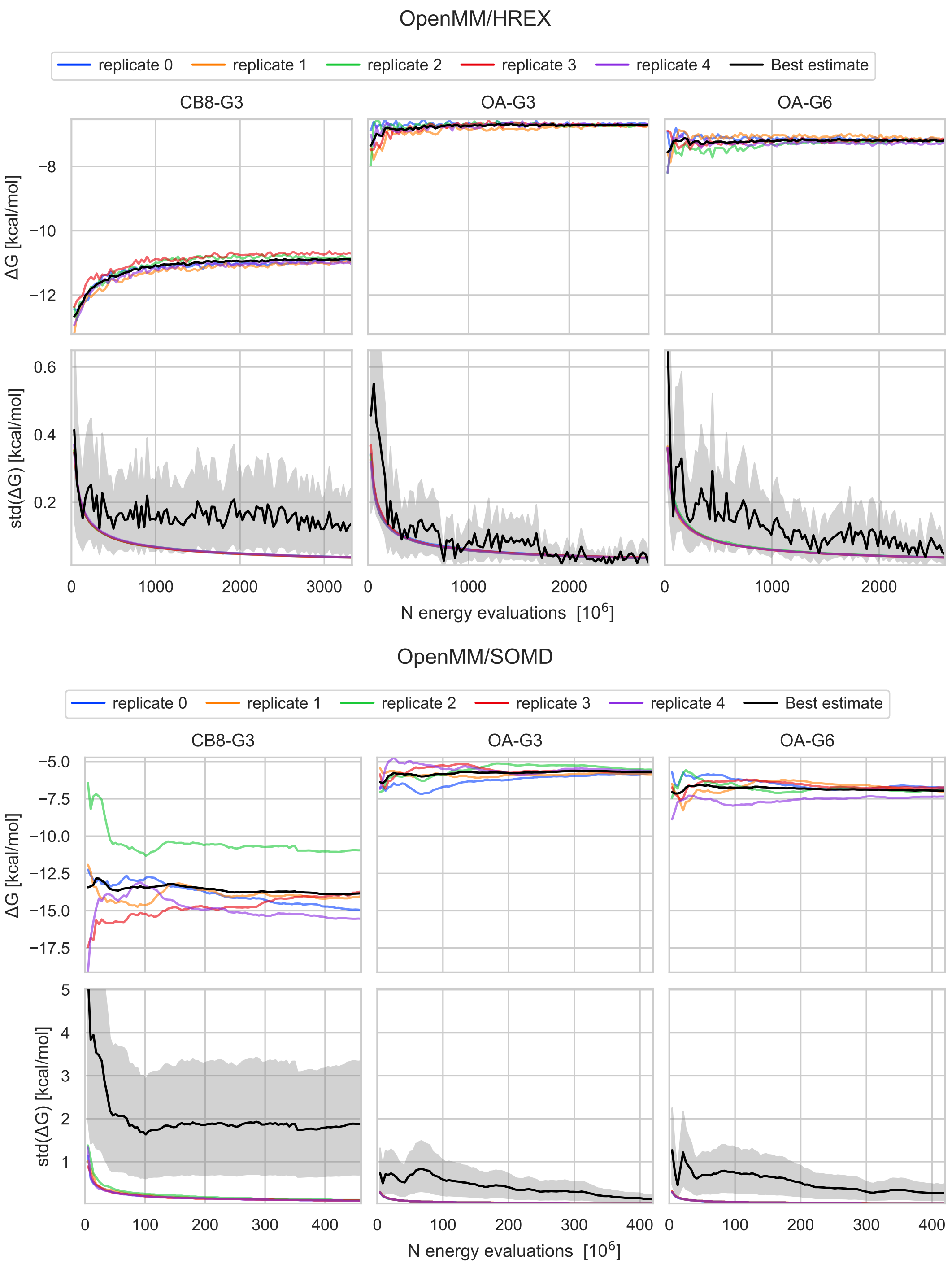
OpenMM/HREX
replicate 0
replicate 1
replicate 2
replicate 3
replicate 4
Best estimate
CB8-G3
OA-G3
OA-G6
ΔG [kcal/mol]
std(ΔG) [kcal/mol]
N energy evaluations [10^6]
OpenMM/SOMD
replicate 0
replicate 1
replicate 2
replicate 3
replicate 4
Best estimate
CB8-G3
OA-G3
OA-G6
ΔG [kcal/mol]
std(ΔG) [kcal/mol]
N energy evaluations [10^6]

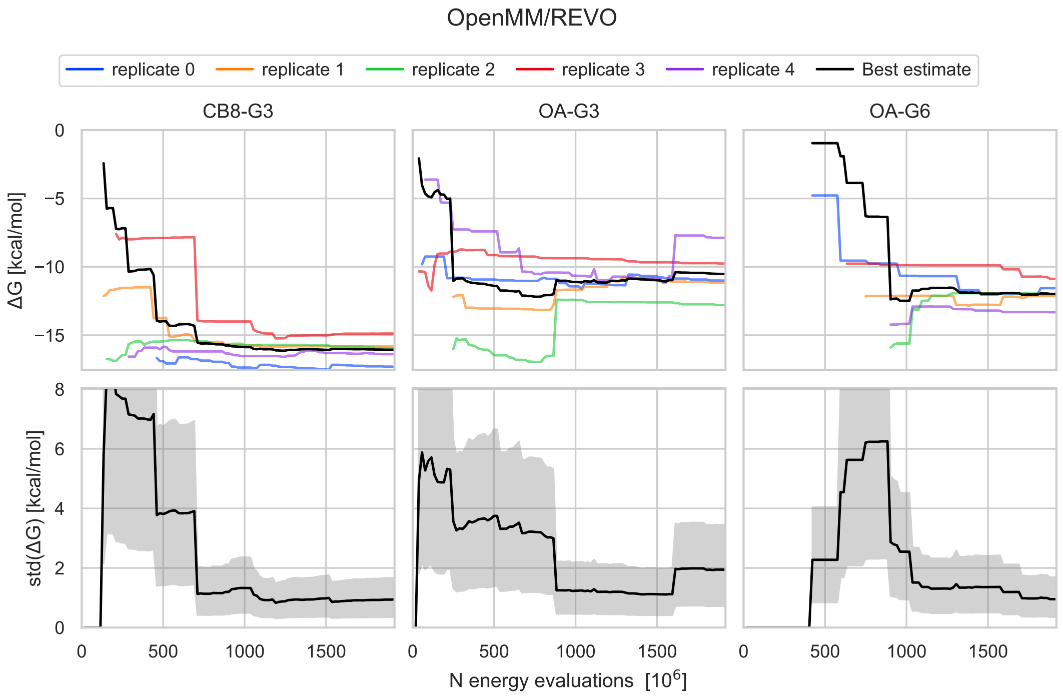
OpenMM/REVO
replicate 0
replicate 1
replicate 2
replicate 3
replicate 4
Best estimate
CB8-G3
OA-G3
OA-G6
ΔG [kcal/mol]
std(ΔG) [kcal/mol]
N energy evaluations [10^6]

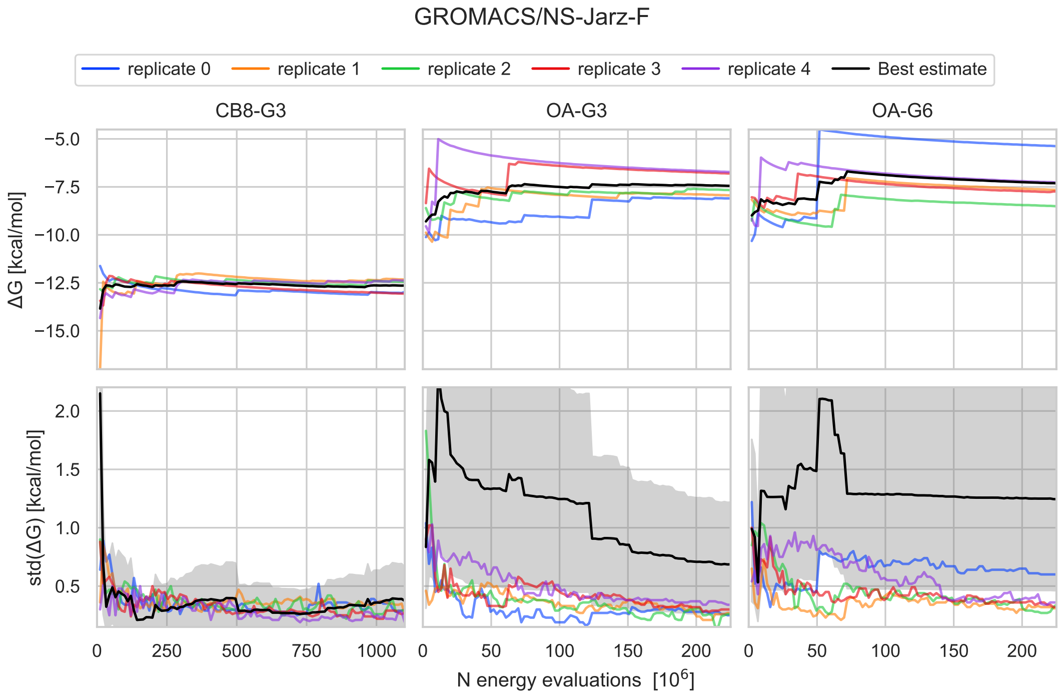
GROMACS/NS-Jarz-F
replicate 0
replicate 1
replicate 2
replicate 3
replicate 4
Best estimate
CB8-G3
OA-G3
OA-G6
ΔG [kcal/mol]
std(ΔG) [kcal/mol]
N energy evaluations [10^6]

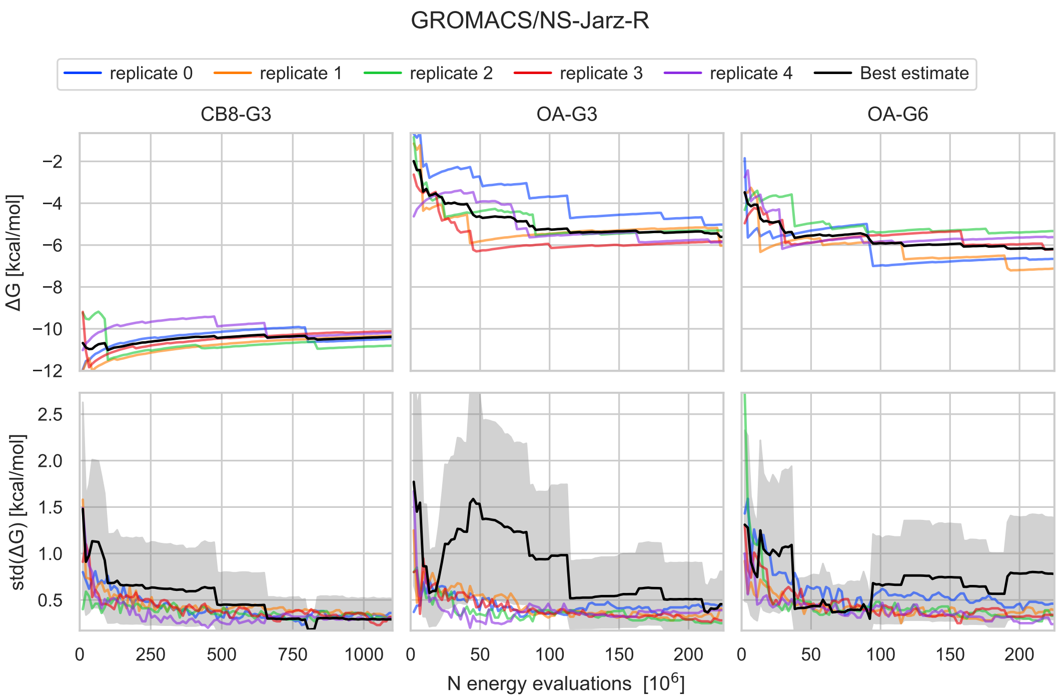
GROMACS/NS-Jarz-R
replicate 0
replicate 1
replicate 2
replicate 3
replicate 4
Best estimate
CB8-G3
OA-G3
OA-G6
ΔG [kcal/mol]
std(ΔG) [kcal/mol]
N energy evaluations [10^6]

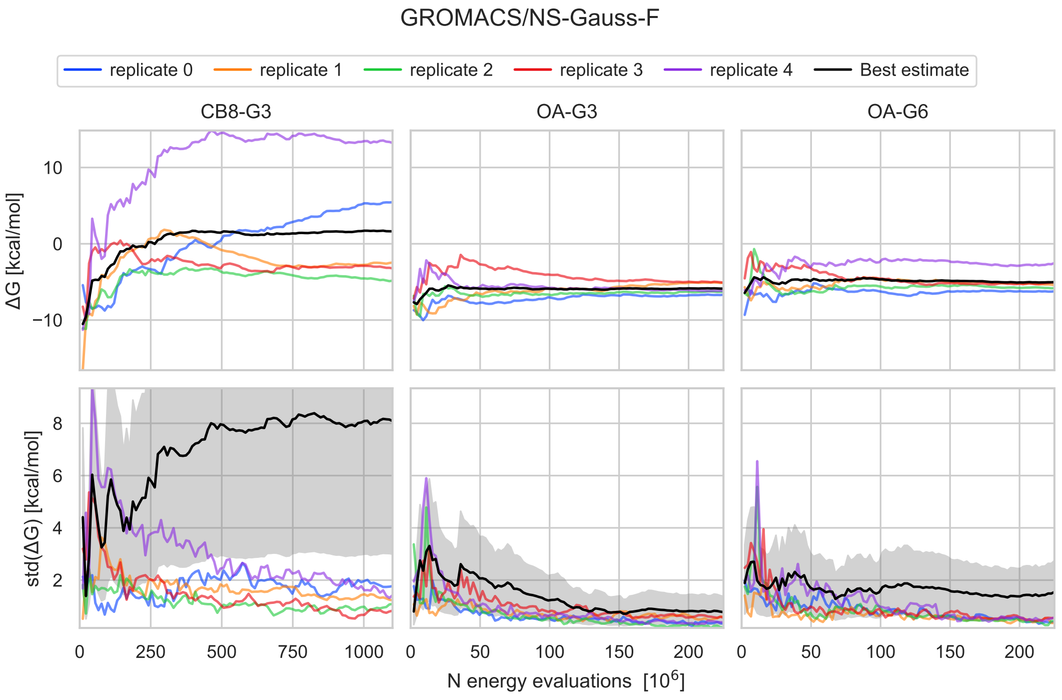
GROMACS/NS-Gauss-F
replicate 0
replicate 1
replicate 2
replicate 3
replicate 4
Best estimate
CB8-G3
OA-G3
OA-G6
ΔG [kcal/mol]
std(ΔG) [kcal/mol]
N energy evaluations [10^6]

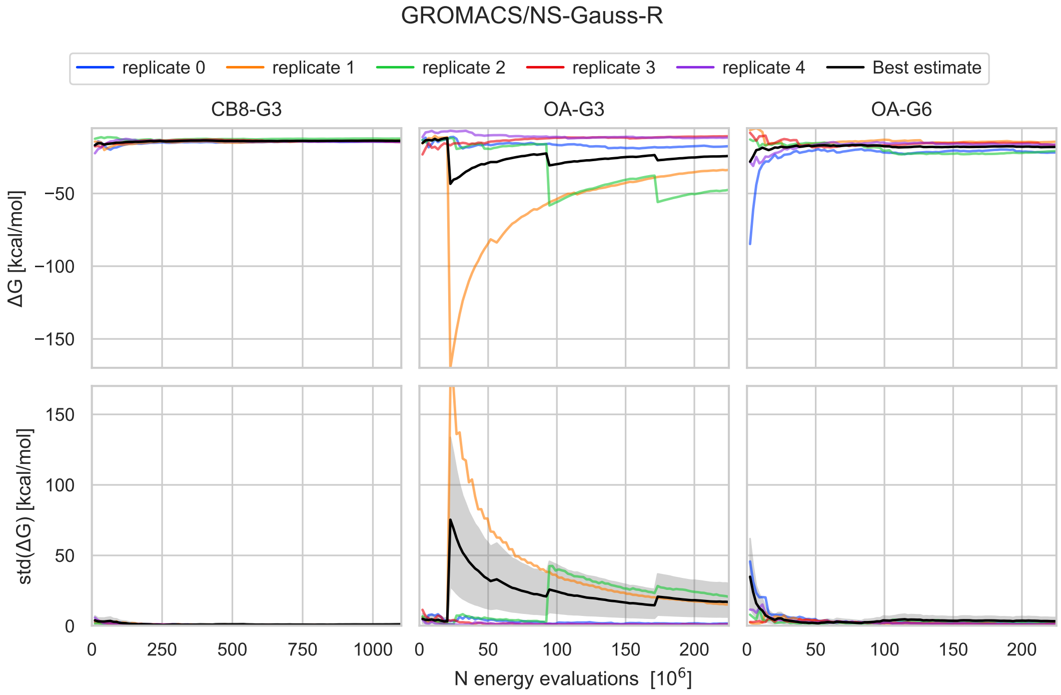


**SI Figure 10: Rare events with large dissipated work impact the free energy.** Work values underlying the $\Delta G$ estimates are depicted together with the free energy values based on the Jarzynski's Gaussian approximation. The orange lines denote work values for the 10 independent simulations that were performed for each initial pose (OA-G3-2 on the left and OA-G3-3 on the right). The work values from the simulations that cause the largest jumps in $\Delta G$ are shown in thick orange lines. The figure illustrates that this estimator is highly sensitive to rare events where large work dissipation is encountered during an alchemical transition.

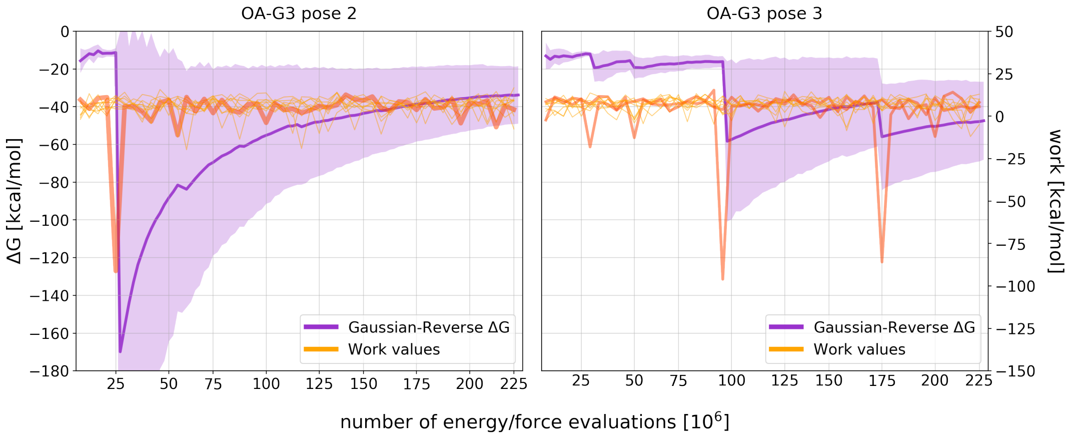

**SI Figure 11: HREX uncertainties as a function of the statistical inefficiency.** Binding free energy trajectories and their uncertainty estimates as a function of the number of force/energy evaluations and statistical inefficiency. The estimates for the individual replicate calculations were computed with MBAR, and they are plotted in pastel colors. The best available estimates (in black) for $\Delta G$ and std($\Delta G$) were taken to be respectively the mean and standard deviation of 5 individual $\Delta G$ trajectories. A 95% chi-based confidence interval is plotted as a gray shaded area around the best estimate of the standard deviation. The MBAR uncertainties are within the confidence interval of the true standard deviation, but they are insensitive to the specific free energy trajectory.

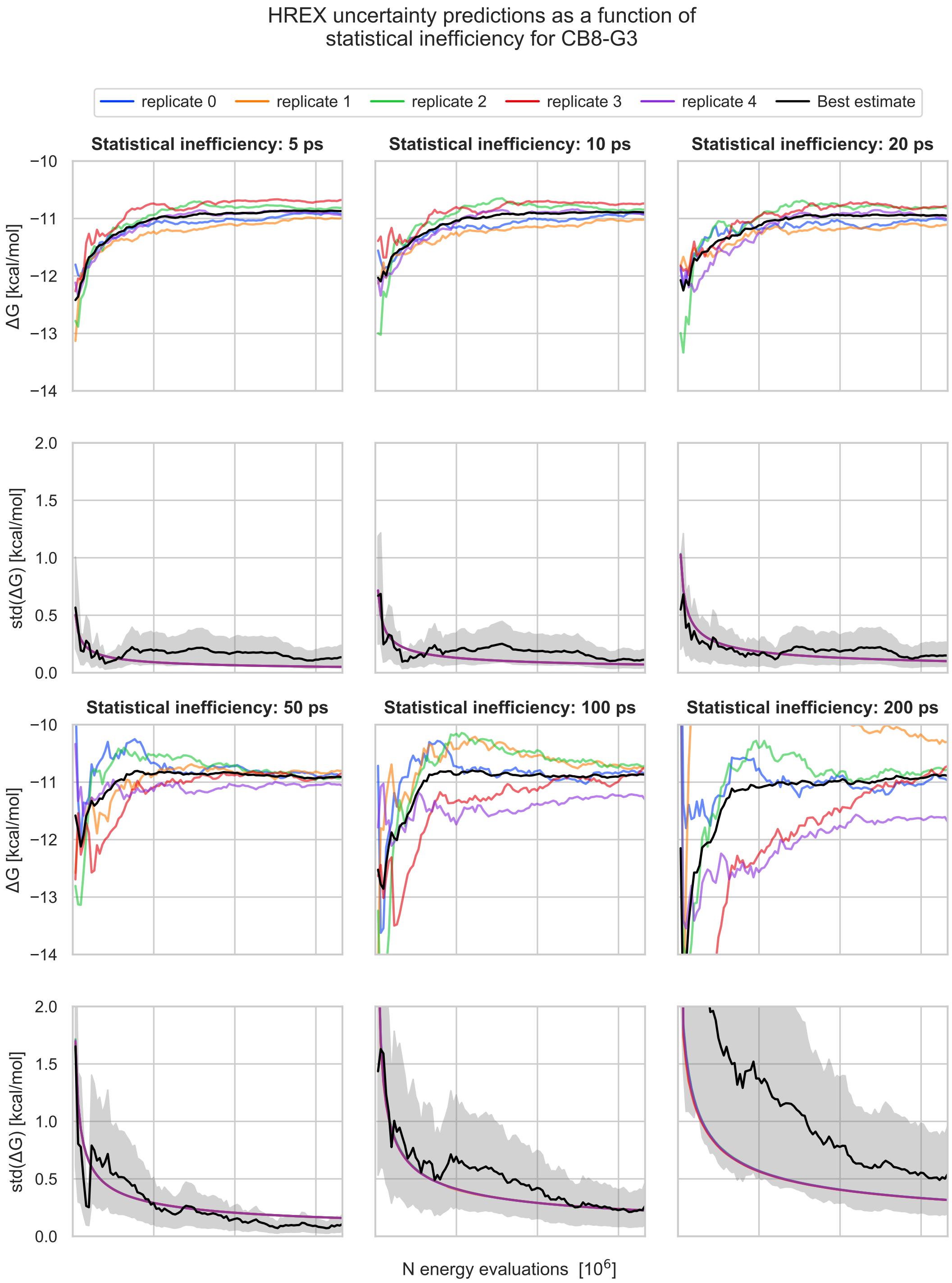

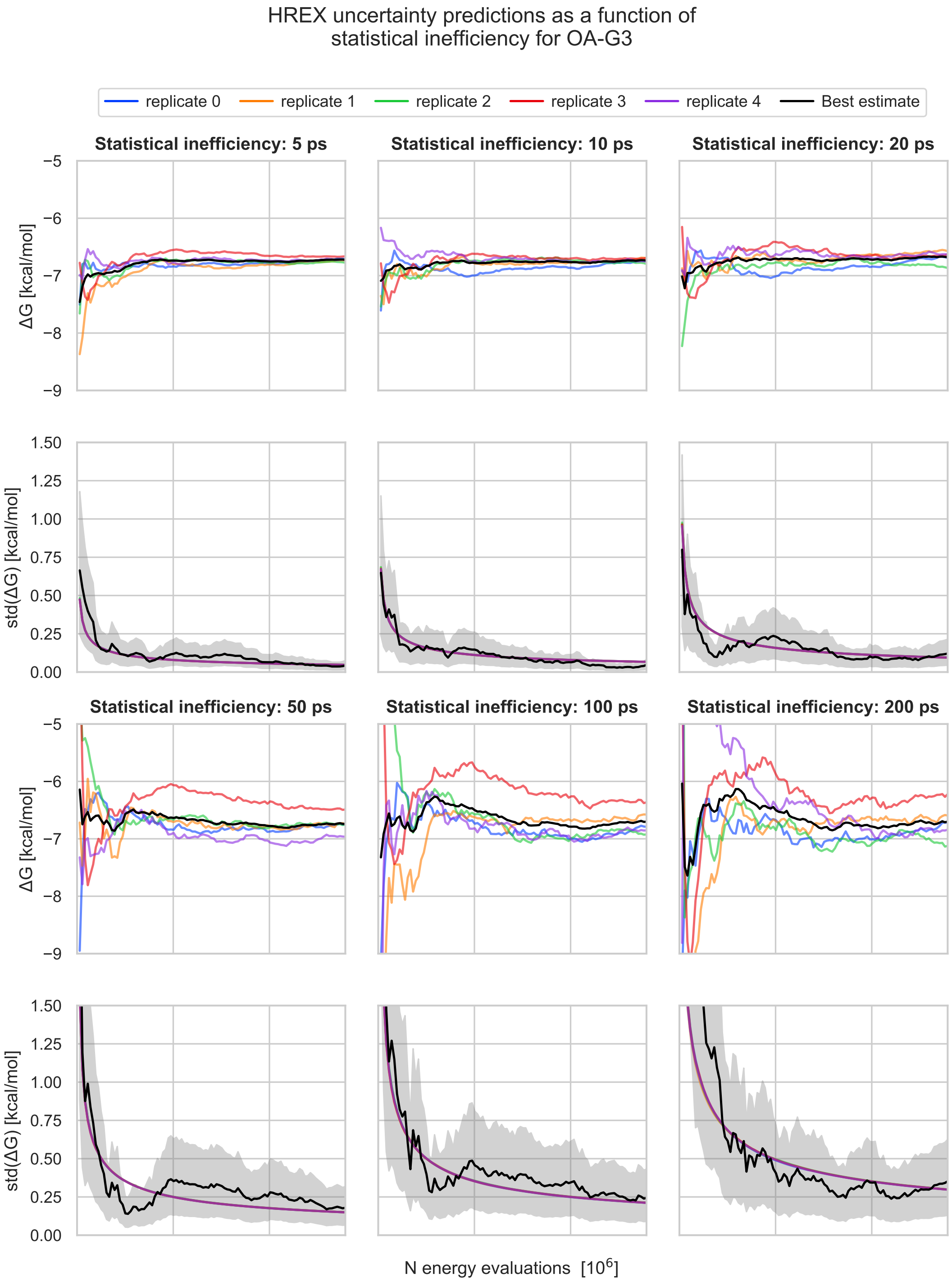
HREX uncertainty predictions as a function of
statistical inefficiency for OA-G3
replicate 0
replicate 1
replicate 2
replicate 3
replicate 4
Best estimate
Statistical inefficiency: 5 ps
Statistical inefficiency: 10 ps
Statistical inefficiency: 20 ps
Statistical inefficiency: 50 ps
Statistical inefficiency: 100 ps
Statistical inefficiency: 200 ps
ΔG [kcal/mol]
std(ΔG) [kcal/mol]
−5
−6
−7
−8
−9
1.50
1.25
1.00
0.75
0.50
0.25
0.00
N energy evaluations [10^6]

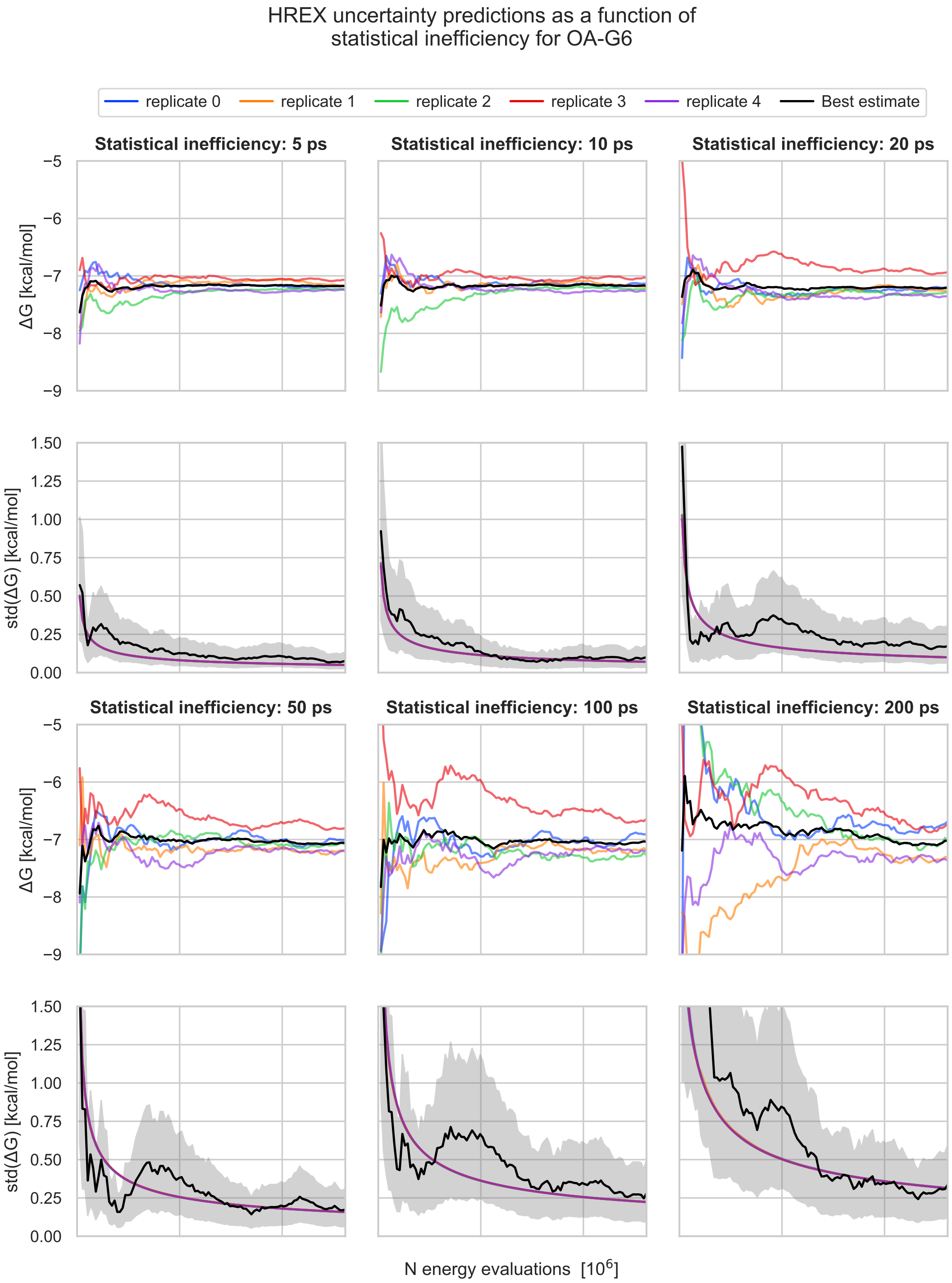

HREX uncertainty predictions as a function of
statistical inefficiency for OA-G6
replicate 0
replicate 1
replicate 2
replicate 3
replicate 4
Best estimate
Statistical inefficiency: 5 ps
Statistical inefficiency: 10 ps
Statistical inefficiency: 20 ps
Statistical inefficiency: 50 ps
Statistical inefficiency: 100 ps
Statistical inefficiency: 200 ps
ΔG [kcal/mol]
std(ΔG) [kcal/mol]
N energy evaluations [10^6]

**SI Table 5: Free energy of replicate SOMD calculations.** Average binding free energy predictions computed from three replicate calculations of each of the five initial conformations of the CB8-G3 system (i.e., CB8-G3-0, ..., CB8-G3-4) plus/minus the standard error of the average. CB8-G3-2 agrees within statistical uncertainty to the other four initial conformations, which suggests that the different free energy trajectory submitted was the result of long correlation time, and that, given a sufficiently long simulation, SOMD would be able to overcome the energy barrier associated to the corresponding slow degree of freedom.

| **initial conformation** | **Average $\Delta G$ [kcal/mol]** |
|---|---|
| CB8-G3-0 | -13.2 ± 0.8 |
| CB8-G3-1 | -14.3 ± 0.3 |
| CB8-G3-2 | -13 ± 1 |
| CB8-G3-3 | -13.7 ± 0.9 |
| CB8-G3-4 | -14 ± 1 |

**SI Table 6: Summary of the expanded ensemble and HREX calculations in NVT and NPT.** Average binding free energy predictions computed from five independent replicate calculations with 95% t-based confidence intervals under different simulation conditions. The barostat column reports whether the simulation was run in NVT (/) or in NPT with a Monte Carlo or Berendsen barostat. The number of grid point for the PME mesh is reported for both the complex and the solvent phase, in this order, when they differ or only once when the same grid was used for both. The simulations in NVT were performed at the average volume sampled by the NPT simulations. Removing the Berendsen barostat caused the free energy of GROMACS/EE to change by 0.6 ± 0.3, while the free energy obtained by YANK with and without the Monte Carlo barostat are identical. Varying the PME parameters did not alter significantly the predictions. The rows labeled with an asterisk refers to the calculations that were used as the final submissions.

| **system** | **method** | **barostat** | **PME FFT grid** | **PME spline order/ error tolerance** | **$\Delta$ G [kcal/mol]** |
|---|---|---|---|---|---|
| OA-G3 | GROMACS/EE | Berendsen | 48x48x48 | 4 / $10^{-5}$ | -6.0 ± 0.2 |
| | *OpenMM/YANK | Monte Carlo | 54x54x54 - 40x40x40 | 5 / $10^{-4}$ | -6.70 ± 0.02 |
| OA-G6 | GROMACS/EE | Berendsen | 48x48x48 | 4 / $10^{-5}$ | -6.9 ± 0.2 |
| | *OpenMM/YANK | Monte Carlo | 54x54x54 - 40x40x40 | 5 / $10^{-4}$ | -7.17 ± 0.05 |
| OA-G3 | GROMACS/EE | / | 48x48x48 | 4 / $10^{-5}$ | -6.6 ± 0.2 |
| | OpenMM/YANK | / | 54x54x54 - 40x40x40 | 5 / $10^{-4}$ | -6.7 ± 0.1 |
| OA-G6 | GROMACS/EE | / | 48x48x48 | 4 / $10^{-5}$ | -7.0 ± 0.2 |
| | OpenMM/YANK | / | 54x54x54 - 40x40x40 | 5 / $10^{-4}$ | -7.15 ± 0.09 |
| OA-G3 | *GROMACS/EE | / | 48x48x48 | 5 / $10^{-5}$ | -6.6 ± 0.1 |
| | OpenMM/YANK | / | 48x48x48 | 5 / $10^{-5}$ | -6.64 ± 0.07 |
| OA-G6 | *GROMACS/EE | / | 48x48x48 | 5 / $10^{-5}$ | -7.0 ± 0.1 |
| | OpenMM/YANK | / | 48x48x48 | 5 / $10^{-5}$ | -7.1 ± 0.1 |

**SI Figure 12: Double decoupling thermodynamic cycle used by YANK for the OpenMM/HREX submission.** In both the complex (top) and solvent (bottom) phases of the thermodynamic cycle, the Coulomb charges were annihilated completely before decoupling the Lennard-Jones interactions. A counterion with charge equal and opposite the net charge of the guest is also decoupled to maintain the box neutrality. In the complex phase, a harmonic restraint was kept activated throughout the calculation, and the end states were reweighted to a state in which the restraint was substituted by a square well restraint that robustly defined the binding site.

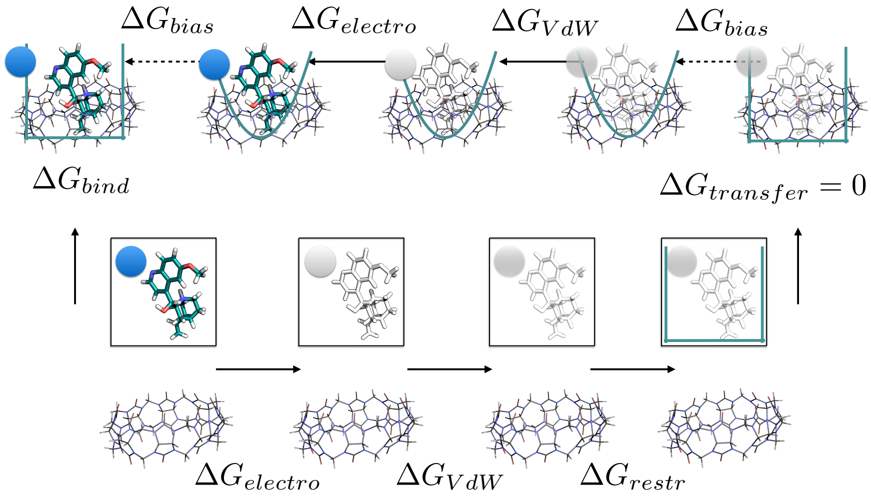


**SI Figure 13: Decomposition of the OpenMM/HREX free energy prediction by phase for CB8-G3.** Complex (orange), solvent (blue), and total (black) mean free energy trajectory computed from the five replicates as a function of the computational cost. Shaded areas represent the standard deviation of the mean free energy. The three axes used to plot the three trajectories are shifted to ease the comparison of the components, but they have the same scale.

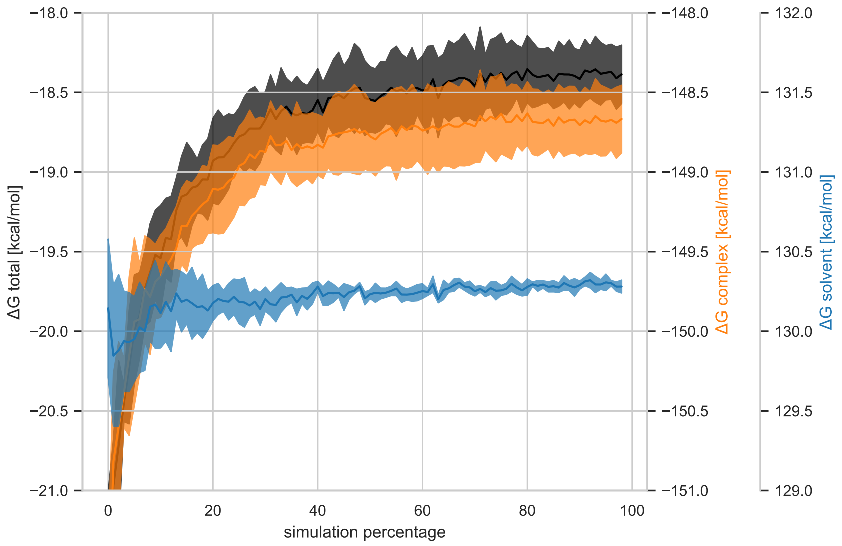

**SI Figure 14: Decomposition of the OpenMM/HREX free energy prediction in entropy and enthalpy for CB8-G3.** Mean enthalpy (orange), entropy (blue), and free energy (black) of binding as a function of the computational cost. The mean is computed from the 5 replicates, and the shaded areas represent the standard deviation of the mean. The axes of entropy and $-T\Delta S$ have the same scale, while the free energy is plotted over a smaller range of values to ease the comparison.

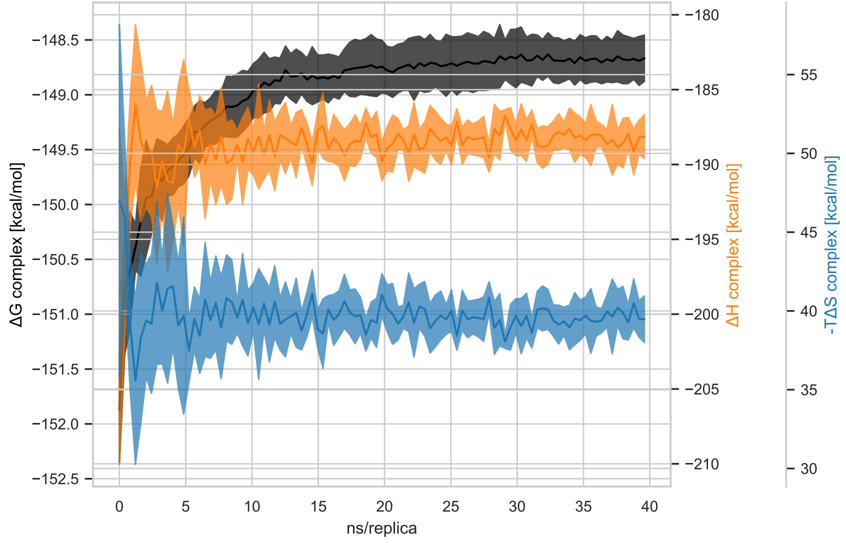

**SI Figure 15: Trajectories of the number of bound waters for each state in CB8-G3-0.** State 0 is the interacting state, while the last state is the decoupled state. The trajectory is plotted against the Hamiltonian replica exchange iteration. The number of bound waters were computed from counting the water molecules with at least one atom within the convex hull of the heavy atoms of CB8.

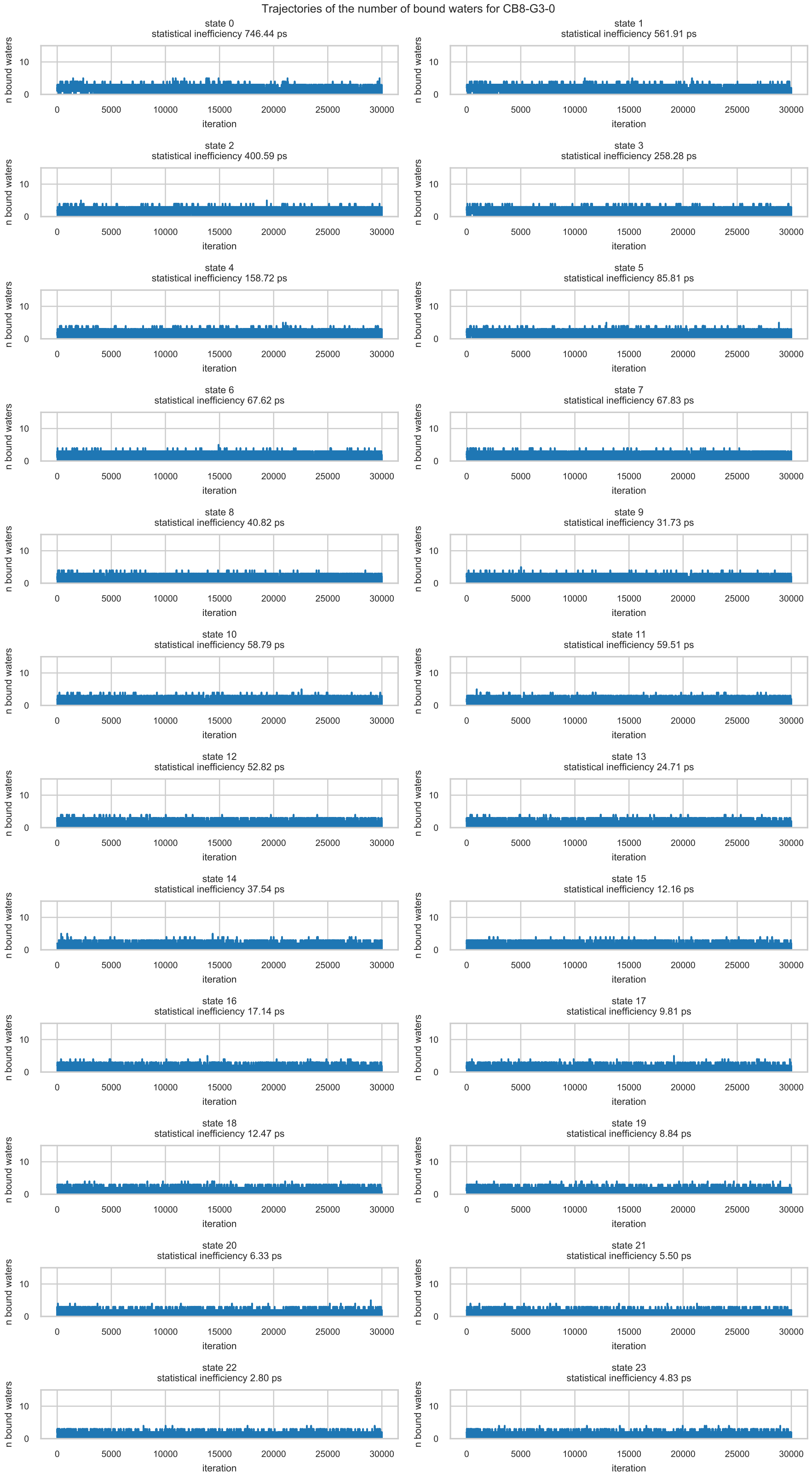

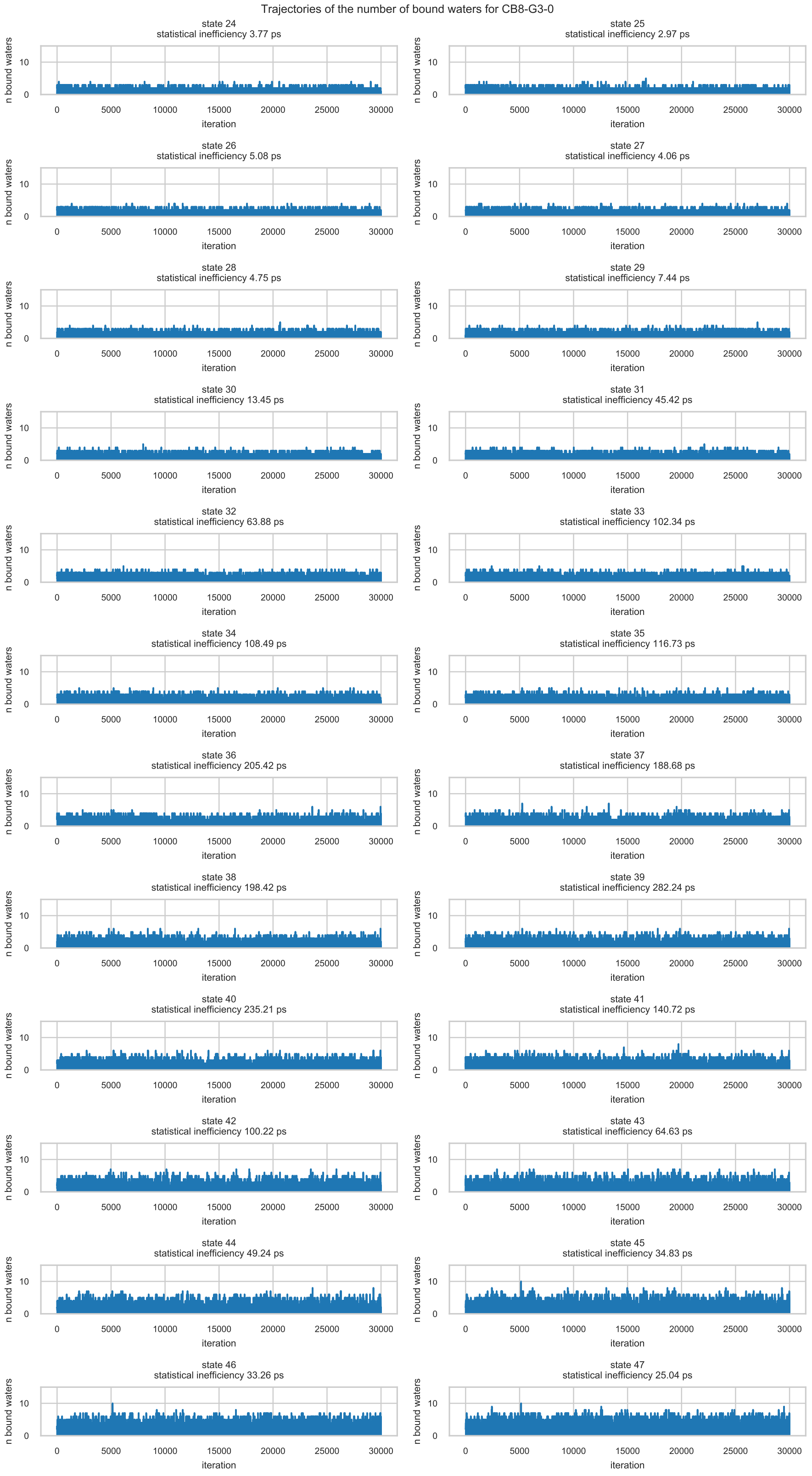
Trajectories of the number of bound waters for CB8-G3-0
state 24
statistical inefficiency 3.77 ps
state 25
statistical inefficiency 2.97 ps
state 26
statistical inefficiency 5.08 ps
state 27
statistical inefficiency 4.06 ps
state 28
statistical inefficiency 4.75 ps
state 29
statistical inefficiency 7.44 ps
state 30
statistical inefficiency 13.45 ps
state 31
statistical inefficiency 45.42 ps
state 32
statistical inefficiency 63.88 ps
state 33
statistical inefficiency 102.34 ps
state 34
statistical inefficiency 108.49 ps
state 35
statistical inefficiency 116.73 ps
state 36
statistical inefficiency 205.42 ps
state 37
statistical inefficiency 188.68 ps
state 38
statistical inefficiency 198.42 ps
state 39
statistical inefficiency 282.24 ps
state 40
statistical inefficiency 235.21 ps
state 41
statistical inefficiency 140.72 ps
state 42
statistical inefficiency 100.22 ps
state 43
statistical inefficiency 64.63 ps
state 44
statistical inefficiency 49.24 ps
state 45
statistical inefficiency 34.83 ps
state 46
statistical inefficiency 33.26 ps
state 47
statistical inefficiency 25.04 ps
n bound waters
iteration
0
10
0
5000
10000
15000
20000
25000
30000

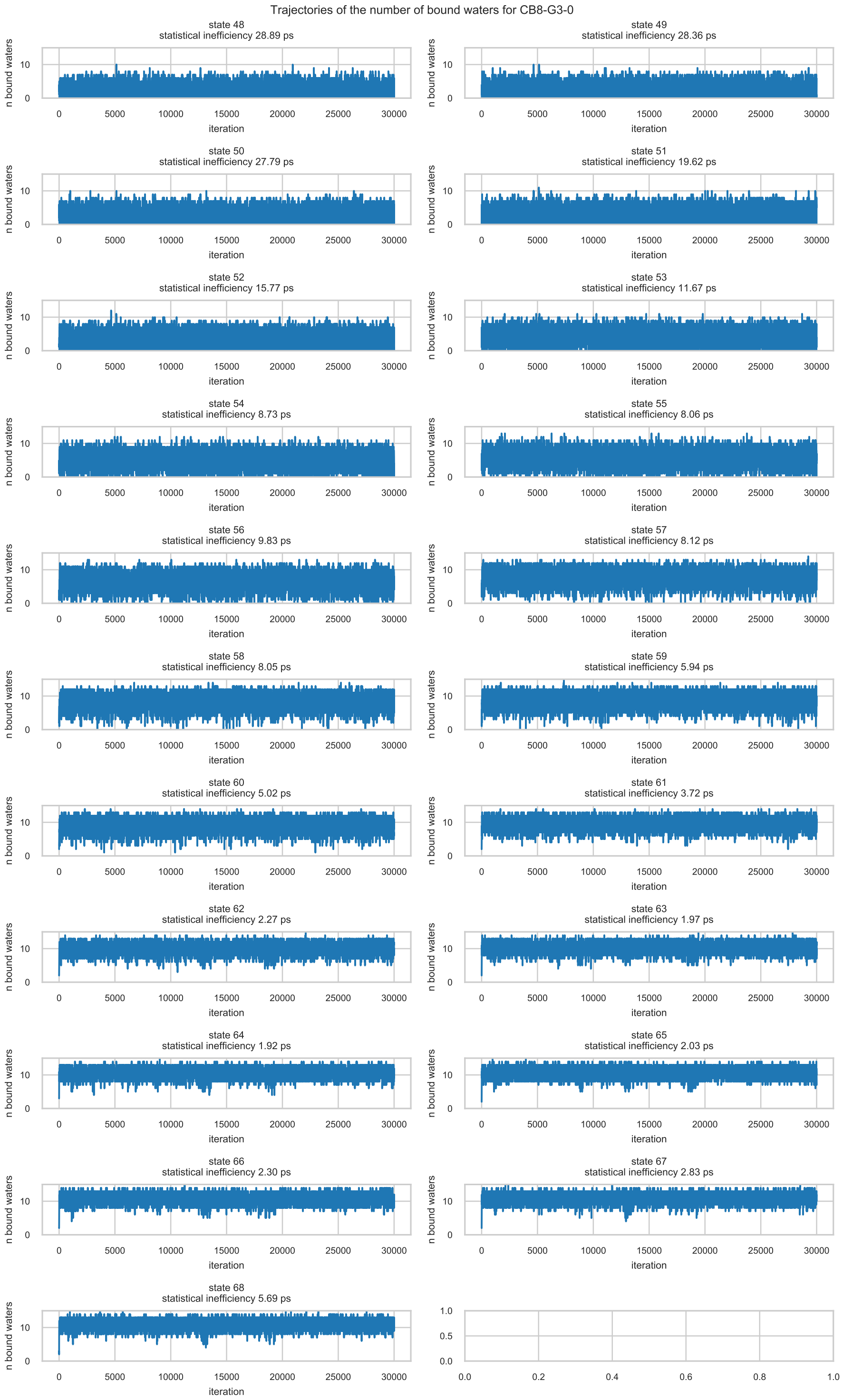
Trajectories of the number of bound waters for CB8-G3-0
state 48
statistical inefficiency 28.89 ps
state 49
statistical inefficiency 28.36 ps
state 50
statistical inefficiency 27.79 ps
state 51
statistical inefficiency 19.62 ps
state 52
statistical inefficiency 15.77 ps
state 53
statistical inefficiency 11.67 ps
state 54
statistical inefficiency 8.73 ps
state 55
statistical inefficiency 8.06 ps
state 56
statistical inefficiency 9.83 ps
state 57
statistical inefficiency 8.12 ps
state 58
statistical inefficiency 8.05 ps
state 59
statistical inefficiency 5.94 ps
state 60
statistical inefficiency 5.02 ps
state 61
statistical inefficiency 3.72 ps
state 62
statistical inefficiency 2.27 ps
state 63
statistical inefficiency 1.97 ps
state 64
statistical inefficiency 1.92 ps
state 65
statistical inefficiency 2.03 ps
state 66
statistical inefficiency 2.30 ps
state 67
statistical inefficiency 2.83 ps
state 68
statistical inefficiency 5.69 ps
n bound waters
iteration

**SI Figure 16: Histograms of the number of bound water by thermodynamic state in the complex phase of OA-G3-0 and OA-G6-0.** The color maps the progression of the alchemical protocol from the bound state (purple, lower part of the histogram) to the discharged state (blue, middle part), where all the charges are turned off but Lennard-Jones interactions are still active, and decoupled state (yellow, upper part). Only the histograms of every other state in the protocol is plotted for clarity.

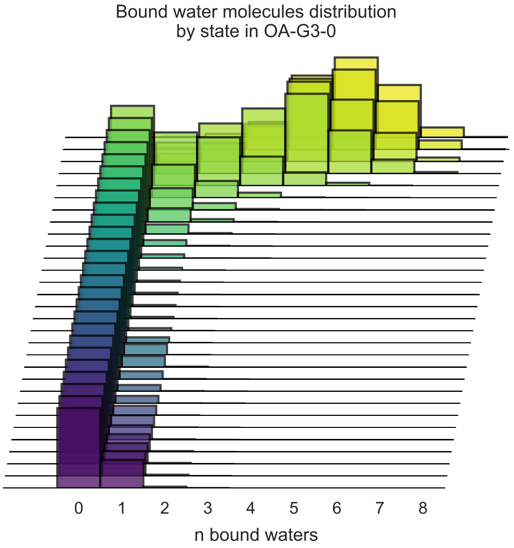


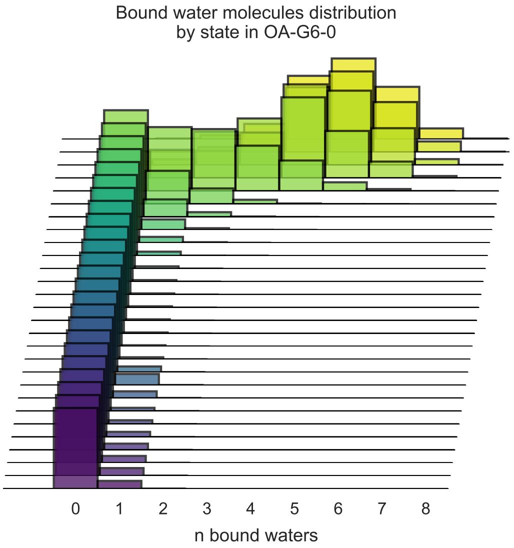
